# Somitic Change Drives Changes in Vertebral Regionalisation in African Cichlids Despite Strong Canalisation of Somite Number

**DOI:** 10.1101/2025.05.23.655800

**Authors:** Callum V Bucklow, Emanuell Duarte Ribeiro, Roger Benson, Berta Verd

**Author notes:** Co-corresponding authors: Callum V Bucklow; Berta Verd.

## Abstract

Vertebrae arise from somites, transient embryonic segments that rhythmically bud from the presomitic mesoderm during axial elongation. The number and identity of vertebrae are ultimately determined by somitogenesis and subsequent anterior-posterior regionalisation, largely governed by *hox* gene expression. Interspecific variation in vertebral count and regionalisation therefore reflects evolutionary changes in somite number and homeotic identity following species divergence. While many macroevolutionary studies have examined homeotic and non-homeotic changes in the vertebral column, few have explored these dynamics in teleosts, despite their exceptional species richness. Using African cichlids as a model, we show that shifts in vertebral regionalisation can arise through modifications to anterior-posterior patterning, but that much of the observed variation is driven by changes in somite number, with homeotic effects emerging largely as a by-product of somitic changes. Moreover, low intraspecific variation in vertebral count, lacking phylogenetic structure, suggests that somitic count variation within species is strongly canalised and has remained consistent throughout the diversification of African cichlids. In addition, we find no correlation between intraspecific variation in vertebral counts and mean vertebral counts, and this variation does not consistently scale with body aspect ratio among individuals. Therefore, intraspecific variation is decoupled from both macroevolutionary patterns of vertebral count evolution and body shape diversification. Together, our findings highlight the dynamic interplay between somitogenesis and homeotic transformations in shaping vertebral diversity and underscore the value of cichlids as a model for understanding the developmental basis of axial evolution in teleosts.

## INTRODUCTION

The vertebral column, a defining feature of all vertebrates, is critical for locomotion, providing structural support for muscle attachment and protecting the spinal cord and nerve roots that transmit motor and sensory signals (Ford, 1937). Vertebrae arise from somites, transient embryonic segments that rhythmically bud from the presomitic mesoderm during axial elongation during somitogenesis (Maroto et al., 2012). Concurrent with somite formation, anterior-posterior (AP) patterning of the somitic mesoderm is directed by overlapping expression domains of *Hox* genes, which specify vertebral identity along the axis (Iimura et al., 2009), a mechanism deeply conserved across vertebrates (Morin-Kensicki et al., 2002; Böhmer et al., 2015; Criswell et al., 2021). *Hox* genes are typically arranged in linear clusters within the genome, and their sequential activation during development reflects the serial organisation of the vertebral column (Iimura et al., 2009). This phenomenon, known as *Hox* collinearity, spatially and temporally regulates AP patterning of the vertebrate body axis (Ye and Kimelman, 2020). Thus, evolutionary changes to both vertebral number and regional identity can arise through modifications to the rate of somite formation or shifts in the expression boundaries of *Hox* genes (Cohn and Tickle, 1999).

Evolutionary modulation of both somitogenesis and anterior-posterior *Hox* patterning has been critical to the evolution of variation in the vertebrate body plan (Naganathan and Oates, 2020). Shifts in *Hox* patterning have been linked to the considerable variability present in vertebral column regionalisation in archosaurs (Böhmer et al., 2015) aligning with morphological domains present in the avian vertebral column (Marek et al., 2021). In mammals, homeotic shifts are sufficient to explain variation in lumbar and thoracic proportions (Narita and Kuratani, 2005) and indeed in the proportions of all post-cervical vertebral counts (Cerbus et al., 2024). In addition to homeotic shifts, changes in vertebral proportions can be brought about by increasing the rate of somitogenesis relative to the *hox* ‘timer’. In Python embryos, for example, an increased rate of somitic budding leads to the formation of a greater number of smaller somites relative to chicken embryos (Gomez et al., 2008; Woltering et al., 2009). In order to support their elongated axes, an anterior shift of *HoxC6* and *HoxC8* in the lateral plate mesoderm prevents forelimb formation, specifying instead the development of thoracic (chest, rib-bearing) vertebrae to support their elongate bodies (Cohn and Tickle, 1999).

Since the number and type of vertebrae is ultimately determined by somitogenesis and subsequent *Hox* regionalisation, differences in the number and type of vertebrae among adults (and other post-embryonic stages) of different species provide evidence of somitic and homeotic changes in development since divergence from their most recent common ancestor. Extrapolation of this onto a larger scale can provide informative abstractions and unique insights into the macroevolutionary dynamics and evolutionary modification of somitogenesis and regionalisation within and between clades (Narita and Kuratani, 2005; Mehta et al., 2010; Soul and Benson, 2017). Whilst a large number of macroevolutionary studies have examined the evolution of the structure of the vertebral column, most have focused on tetrapods (Narita and Kuratani, 2005; Jones et al., 2018b,a; Cerbus et al., 2024) and few have examined the macroevolutionary patterns of vertebral count and identity in ray-finned fish (Actinopterygii) (Ward and Brainerd, 2007; Mehta et al., 2010).

Ray-finned fish (Actinopterygii) are the most speciose class of extant vertebrates, accounting for over 50% of described species, of which 99% are teleosts (Near and Thacker, 2024). Understanding the macroevolutionary and developmental dynamics of the teleost vertebral column is therefore central to elucidating morphological and functional diversification in a highly diverse group. Unlike in tetrapods, that are defined by the presence of five well defined vertebral types: cervical, thoracic, lumbar, sacral and caudal (tail) (Cerbus et al., 2024), the teleostean vertebral column has traditionally been divided into two domains: the precaudal and caudal. Precaudal vertebrae are characterised by ventrolateral basapophyses that serve as attachment sites for ribs, which protect the viscera and provide anchorage for muscles and tendons. Caudal vertebrae, in contrast, are defined by the presence of a closed haemal arch formed by haemal spines (see Figure 1A), which provide articulation points for the anal and caudal fins required for swimming (Ford, 1937). Therefore, changing proportions of precaudal and caudal vertebrae are a useful proxy to investigate the evolutionary importance of homeotic transformations in teleost evolution.

**Figure 1.**
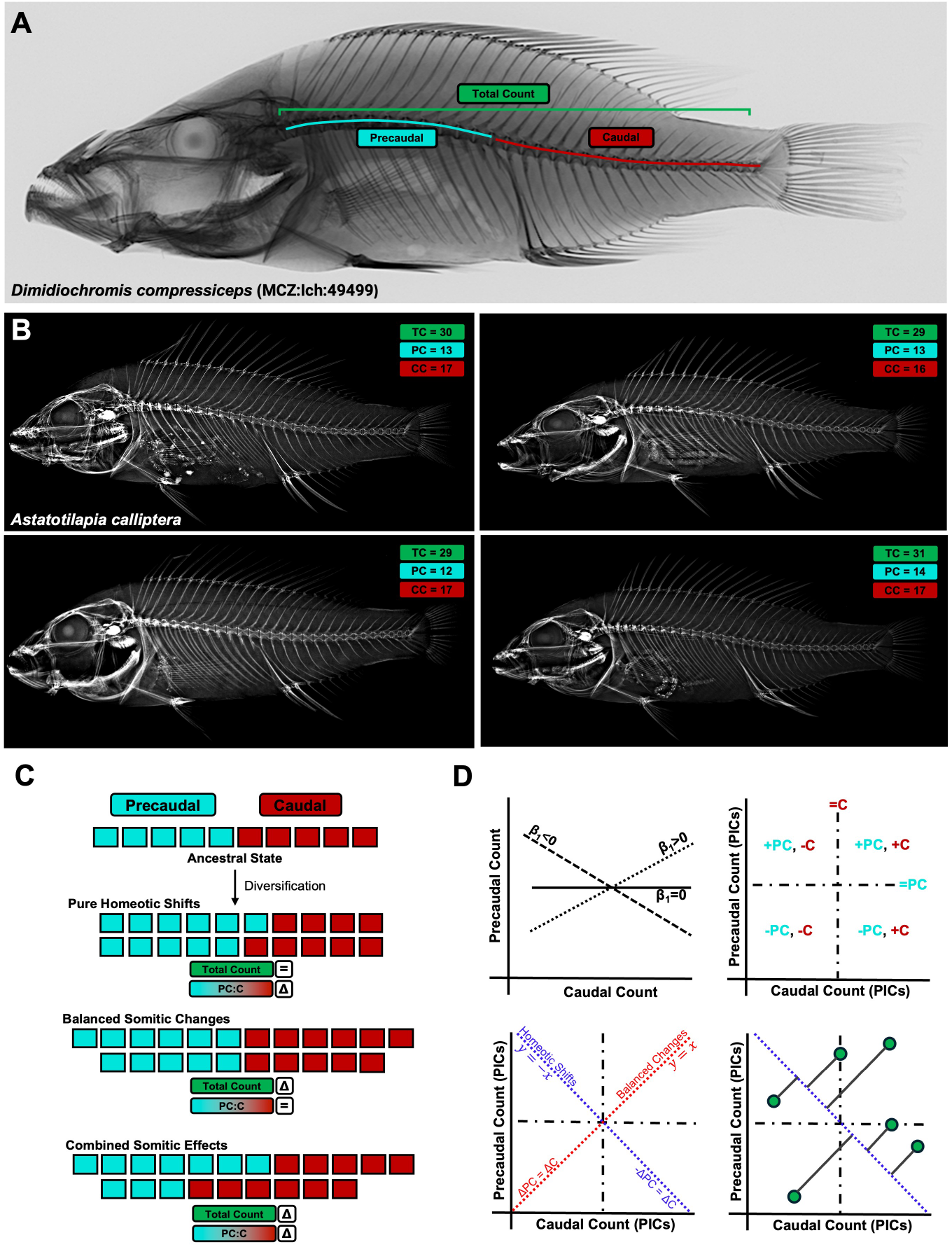
Vertebral column structure in cichlids. (A) X-ray of *Dimidiochromis compressiceps*, showing precaudal, caudal, and total vertebral counts. (B) X-rays of four individuals of *Astatotilapia calliptera*, illustrating intraspecific variation in total vertebral count and proportions of precaudal and caudal vertebrae. (C) Evolutionary changes in vertebral number and regionalisation can arise through three primary modes: Pure Homeotic Shifts, in which the relative proportions of precaudal and caudal vertebrae change without altering the total count, consistent with changes to anterior-posterior patterning independent of somitogenesis; Balanced Somitic Changes, where precaudal and caudal counts increase or decrease proportionally, suggesting changes in somite number without affecting regional identity boundaries; and Combined Somitic Effects, where changes in precaudal and caudal counts are disproportionate, reflecting shifts in regional proportions that arise as a consequence of altered somitogenesis, without necessarily involving changes to anterior-posterior patterning. (D) Schematics clarifying Figure 2. Top left: if precaudal and caudal counts have evolved independently they do not co-vary (*β*_1_ = 0). Top right: the position of nodes in different quadrants reflects changes in precaudal and caudal proportions relative to the ancestral state. Bottom left: nodes falling along the *y* = −*x* and *y* = −*x* lines represent pure homeotic shifts and balanced somitic changes, respectively. Bottom right: nodes deviating perpendicularly from the *y* = − *x* line indicate changes in total vertebral count accompanied by a shift in regionalisation; the greater the perpendicular distance, the greater the magnitude of the somitic change.

While most studies of vertebral evolution have focused on interspecific patterns across broad phylogenetic scales, relatively few, if any, have examined patterns of intraspecific variation in vertebral number, despite widespread evidence of such variation across vertebrates (Slijepčević et al., 2015; Hu et al., 2016; Sosa and Hospitaleche, 2024), including in teleosts (Yamahira et al., 2006; Tibblin et al., 2016; Oliver, 2024) (see Figure 1B). Intraspecific variation in total vertebral count necessarily reflects underlying variation in somitogenesis, since the number of vertebrae corresponds to the number of embryonic somite pairs formed during early development (Morin-Kensicki et al., 2002). In teleosts, this relationship is particularly direct: because resegmentation does not occur following somite specification, the total number of vertebrae corresponds to the number of somitic pairs minus three, with the anterior-most somites contributing to the basioccipital region of the skull (Woltering et al., 2018). Examining the phylogenetic distribution and structure of intraspecific variation in vertebral counts therefore offers a powerful but unexplored means to investigate how axial patterning and developmental constraint have evolved in vertebrates. However, despite its potential significance, the extent, developmental basis, and evolutionary implications of intraspecific variation in precaudal, caudal and total vertebral counts in teleosts remains poorly understood and represents a key gap in our understanding of vertebrate axial evolution.

African cichlids (Ovalentaria, Cichliformes, Cichlidae, Pseudocrenilabrinae) represent one of the most species-rich and morphologically diverse groups of teleost fishes, found across the rivers and lakes of Africa and the Middle East (Astudillo-Clavijo et al., 2023). Much of this diversity has arisen within the adaptive radiations of the East African Great Lakes Tanganyika, Malawi, and Victoria, which together represent hundreds of endemic species (McGee et al., 2020). The radiations of Lakes Malawi and Victoria, in particular, have diverged rapidly, with estimated ages of approximately 800,000 and 100,000 years, respectively (Malinsky et al., 2018; Meier et al., 2017). Among their many axes of phenotypic diversity, African cichlids exhibit remarkable variation in axial skeletal morphology, including in total vertebral counts, the relative proportions of precaudal and caudal vertebrae, and in elongation of the body (Oliver, 2024; Bucklow et al., 2025). We previously showed that while total vertebral counts evolve under lineage-specific stochastic rates within each lake radiation, these rates are highest among riverine lineages, contributing to their especially diverse axial morphologies. Moreover, increases in total vertebral counts, often driven by increased caudal counts, are associated with elongation of the body axis, a trait linked to pelagic, demersal, and piscivorous lifestyles (Bucklow et al., 2025). However, despite investigating the macroevolutionary dynamics of total count evolution, we did not make any inferences about the evolution of the developmental mechanisms that may underlie these evolutionary patterns. Understanding the developmental basis of vertebral variation is therefore essential to fully understand the evolution of axial morphology in African cichlids.

Here we show that across African cichlids, precaudal and caudal vertebral counts evolve largely independently of each other. While variation in total vertebral count is often accompanied by changes in regionalisation, the relationship between precaudal-caudal proportions and total vertebral count is weak, non-linear, and likely highly lineage-specific. We define three distinct modes of evolutionary change in vertebral column evolution (Figure 1C). First, pure homeotic shifts involve proportional changes in precaudal and caudal vertebral counts, shifting the boundary between regions without altering total vertebral number. Second, balanced somitic effects reflect the gain or loss of vertebrae, with precaudal and caudal counts changing in the same direction and proportion. Finally, combined somitic effects involve unbalanced changes in precaudal and caudal counts, altering both total vertebral number and regional identity. We detect examples of all three modes across the African cichlid phylogeny, but find that evolutionary changes to vertebral column structure most often involve combined somitic effects, whereas pure homeotic shifts and balanced somitic changes are comparatively rare. These patterns are especially pronounced within the haplochromine radiations of Lakes Victoria and Malawi, where the highest rates of combined change are observed. In addition, despite the large diversity of total counts across species, intraspecific variation is low, lacks phylogenetic structure, and has remained consistent during the diversification of African cichlids. Furthermore, vertebral count variability within species does not scale with body elongation, nor does it differ between lineages with divergent total counts. Together, these results indicate that while vertebral numbers and regionalisation have evolved dynamically and extensively across cichlid lineages, intraspecific variability in somitic counts remains strongly constrained and decoupled from macroevolutionary patterns of body shape and vertebral diversification.

## METHODS AND MATERIALS

All analyses were conducted in R (v4.2.0) (R Core Team, 2022). Phylogenetic analyses used the cichlid family tree constructed by McGee et al. (2020) as the tree is ultrametric, time-calibrated and includes all three of the major lacustrine radiations of African cichlids (Lake Tanganyika, Lake Malawi and Lake Victoria), as well as representatives from every currently recognised tribe in the African subfamily. All vertebral count data is from a previous study, which includes all necessary information about how the counts were conducted and includes landmarked images of all the specimens (Bucklow et al., 2025). All code and data have been deposited alongside this manuscript as a self-contained R project.

### Testing modularity of precaudal and caudal domains

To explore the macroevolutionary relationship between vertebral column regionalisation and somitogenesis, we first needed to confirm that changes in precaudal and caudal vertebrae could serve as reliable proxies for homeotic and somitic changes. If counts in either domain have changed independently of the other, then changes in the relative numbers of precaudal and caudal vertebrae can be used as a proxy for homeotic shifts (and co-occurring somitic changes). We also examined the relationship between the ratio of precaudal:caudal vertebrae (ln[Precaudal]-ln[Caudal]) and the ln[Total Count], to evaluate whether increases in total vertebral count were associated with shifts in the relative allocation of precaudal versus caudal vertebrae. We used phylogenetic generalised least squares (PGLS) to evaluate the relationship between the univariate traits, using the *pgls* function in the R package *caper* (v1.0.1) (Orme et al., 2018). To account for phylogenetic signal, we estimated *λ* (Pagel, 1999) for the covariance matrix, with *δ* and *κ* fixed at 1. We added a quadratic term to the PGLS model for the ratio of ln[Precaudal] - ln[Caudal] as a function of ln[Total Count], as both the OLS and linear PGLS model fits did not visually capture the data structure well (see Figure 2B). The linear term was marginally non-significant (*β*_1_ = 0.304, P = 0.055) but the quadratic term was significant (*β*_2_ = 0.460, P *<* 0.001) and the model overall provided a considerably better fit than the linear PGLS model (ΔAIC *>* 3).

**Figure 2.**
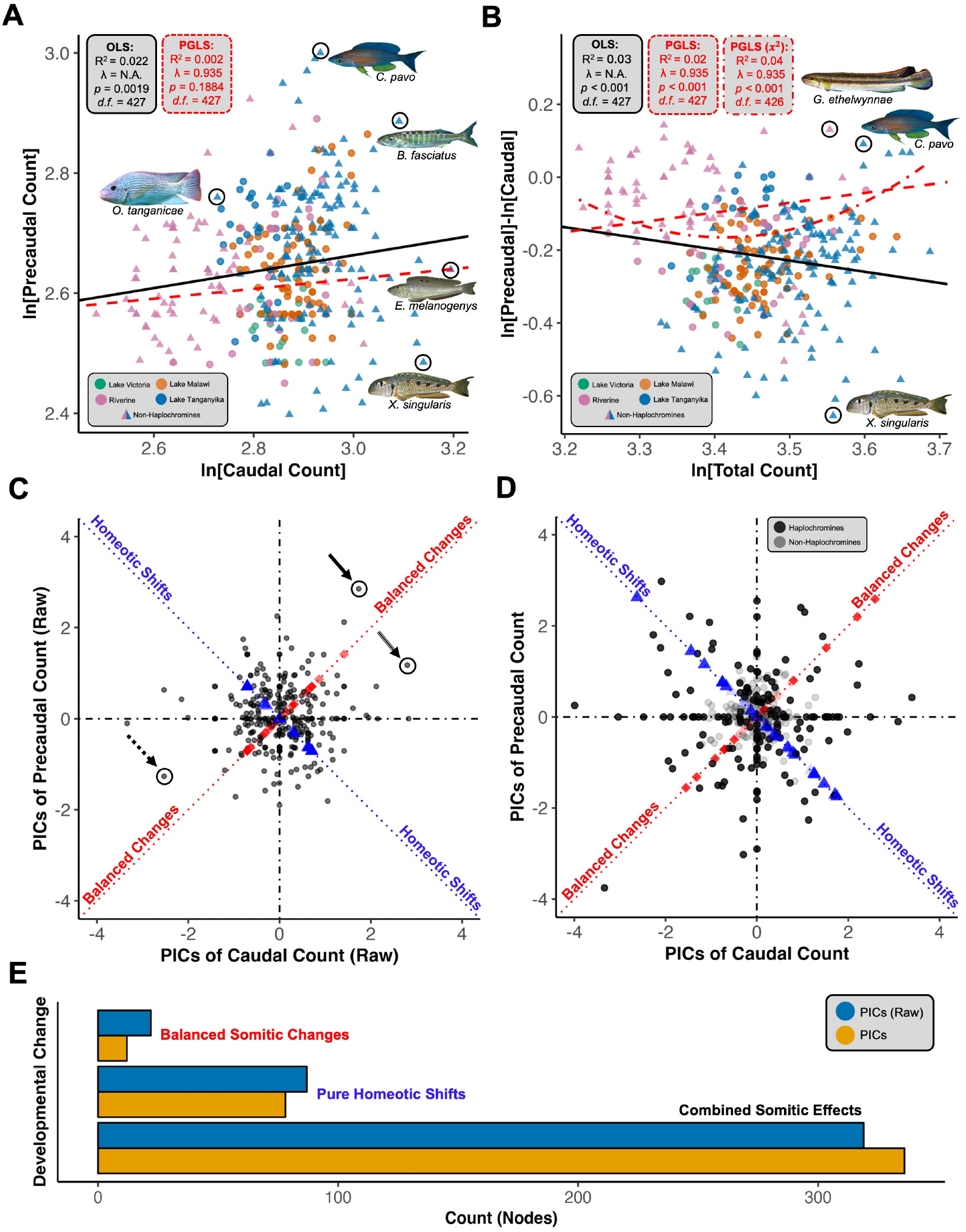
Homeotic and somitic changes in African cichlids. (A) Scatterplot of ln[Precaudal] and ln[Caudal] vertebral counts and (B) ln[Precaudal]-ln[Caudal] (PC:C ratio) with regression lines: black (OLS) and red, dashed (PGLS). Scatterplot of modal, raw (C) and time-calibrated (D) precaudal and caudal vertebral phylogenetic independent contrasts (PICs). Points (blue triangles) on the *y* = −*x* line (blue, dotted) indicate pure homeotic shifts, points (red diamonds) on the *y* = *x* line (red, dotted) indicate balanced changes, perpendicular deviations from either line represent co-occurring somitic changes. Arrows match nodes exemplified in Figure 3. (E) Bar chart showing the number of nodes where each developmental change (defined in Figure 1) are predicted to have occurred. Blue represents the number of nodes where each developmental change is predicted to have occurred, based on PICs where the branch length is equal to 1 (i.e. the raw magnitude of change) and in orange are the predictions based on the time-corrected PICs. *B, Bathybates*; *C, Cyprichromis*; *E, Enantiopus*; *G, Gobiocichla*; *L, Lamprologus*; *O, Oreochromis*; *X, Xenotilapia*. See Supplementary Materials for image credits.

### Identification of pure homeotic shifts, balanced somitic changes and combined somitic effects

To assess the phylogenetic scale of homeotic shifts and test whether their rates varied across the tree, we used standardised phylogenetic independent contrasts (PICs) (Felsenstein, 1985) to quantify changes in precaudal and caudal vertebral counts. PICs were calculated as the difference in the modal raw counts between nodes, standardised by their expected standard deviation, using the R package *ape* (v5.7.1) (Paradis and Schliep, 2019), where the contrast sign indicated the direction of change in vertebral counts at that respective node. We calculated PICs in two ways: first, using the original, time-calibrated branch lengths; and second, with all branch lengths set to 1. The time-calibrated PICs allowed us to investigate the temporal scale at which homeotic and somitic effects could occur, since PICs represent point estimates of the evolutionary rate of change in precaudal and caudal vertebral counts (Freckleton and Harvey, 2006). In contrast, using equal branch lengths enabled us to assess the raw magnitude of changes across the phylogeny, independent of divergence time.

Following Soul and Benson (2017), we defined homeotic shifts as nodes where precaudal and caudal vertebrae contrasts changed disproportionally. Specifically, instances where a predicted loss of precaudal vertebrae was accompanied by a concurrent, equal-magnitude increase in caudal vertebrae were considered ‘pure homeotic shifts’, as they did not alter the total vertebral count (and thus the number of somites). In contrast, we defined ‘balanced somitic changes’ as nodes where precaudal and caudal count contrasts were predicted to have changed, albeit proportionally, maintaining the ratio of precaudal-caudal vertebrae but not the total vertebral counts. Nodes where precaudal and caudal count contrasts changed disproportionately, implying homeotic shifts with co-occurring somitic changes, were defined as ‘combined somitic effects’. For example, if the loss of one precaudal vertebra was accompanied by the addition of two caudal vertebrae, the total vertebral count (and also the number of somites) increased, indicating a co-occurring somitic change (see Figure 1C).

In practical terms, we plotted precaudal count PICs against those of caudal vertebral counts (see Figure 1D for summary). Pure homeotic shifts were identified as nodes falling on the *y* = − *x* line, indicating equal and opposite changes in precaudal and caudal counts. Balanced changes were nodes located on the *y* = −*x* line, where precaudal and caudal counts changed proportionally, preserving their ratio. Nodes deviating perpendicularly from the *y* = −*x* line were considered homeotic shifts accompanied by somitic changes, as both vertebral column regionalisation and the number of either precaudal in either both or one vertebrae changed. The magnitude of deviation along this perpendicular axis represented the magnitude of the somitic change for nodes where combined somitic effects were identified. Greater perpendicular distances indicate a larger additive or subtractive somitic effect (i.e. co-occurring somitic change) (Soul and Benson, 2017). For PICs calculated using estimated divergence times, a greater perpendicular deviation indicates that substantial somitic changes occurred over relatively short evolutionary timescales. To quantify the magnitude of co-occurring somitic changes (for ‘combined somitic effect’ nodes), we calculated the perpendicular distance of each node from the *y* = −*x* line, using the formula: 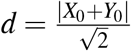, where *X*_0_ and *Y*_0_ are the standardised contrasts for caudal and precaudal vertebrae, respectively. A value of *d* = 0 corresponded to pure homeotic shifts, while *d* = 1 (aligned with *y* = *x*) represents balanced changes. A perpendicular distance 0 < *d*≠ 1, indicated homeotic shifts with co-occurring somitic changes.

### Quantification of intraspecific variation in vertebral counts

Intraspecific variation in vertebral counts is present in African cichlids (Oliver, 2024). Since the total number of vertebrae is ultimately defined by the number of somite pairs formed during somitogenesis, the presence of intraspecific variation in vertebral counts implies flexibility in somitogenesis which could be the source of interspecific variation in total vertebral counts. We had previously shown that vertebral addition is necessary for elongation of the fusiform body in African cichlids (Bucklow et al., 2025). If increased vertebral counts evolved through positive selection acting on intraspecific variation in somite number, we would expect lineages with exceptionally high vertebral counts such as *Rhamphochromis, Serranochromis*, and *Bathybates* to retain elevated levels of intraspecific variation in vertebral number. Alternatively, if higher vertebral counts arose through a directional shift followed by purifying selection against lower counts (e.g., due to ecological or functional constraints), we would expect this to result in a narrowing of the trait distribution, and thus reduced intraspecific variation within those high-count lineages. To explore this, however, we needed to characterise the intraspecific variation in African cichlids, determine whether it is linked to interspecific vertebral counts and test whether intraspecific variation has been modified over the course of their diversification.

We filtered our dataset of vertebral counts to only include species for which we had at least five individuals, leaving us with 277 species retained on the phylogeny after pruning. Interspecific variation in total counts remained normally distributed in this subset of species (Supplementary Figure 1A, Shapiro-Wilks test, W = 0.995, *p* = 0.573) and covered the full range of vertebral counts present in African cichlids (Oliver, 2024; Bucklow et al., 2025). Given that sample sizes were highly variable among species (n = 5–57 individuals), we tested whether species with larger sample sizes exhibited inflated estimates of intraspecific variation. This was particularly important because some specimens, especially from museum collections, may have been misidentified, potentially increasing apparent variation. A linear model was fit using log-transformed values of the total count standard deviation and sample size to account for potential non-linearity and heteroscedasticity (Supplementary Figure 1B). We found a weak and marginally non-significant positive correlation between intraspecific variation and sample size (*R*^2^ = 0.010, *β*_1_ = 0.04584, *p* = 0.056, *d.f.* = 267), and highly non-significant results from a non-parametric Spearman’s rank correlation test (*ρ* = 0.054, *p* = 0.37). Therefore, increased sampling does not systematically inflate observed intraspecific variation.

To quantify intraspecific variation in total vertebral counts, we used the coefficient of variation (CV; CV = *σ/μ*) rather than the standard deviation. Since the standard deviation is an absolute measure, it can be biased by species with higher mean vertebral counts, making cross-species comparisons less meaningful. In contrast, the CV expresses variation relative to the mean, enabling standardised comparisons and providing a clearer assessment of relative developmental variability in vertebral number across species. We did, however, calculate the variance (*σ* ^2^) of precaudal and caudal counts to estimate their respective contributions to total vertebral count variance. Total count is the sum of precaudal and caudal counts. Therefore, total variance can be expressed as *σ* ^2^ 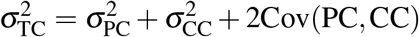. However, precaudal and caudal counts did not covary when corrected for phylogeny (Figure 2A). Thus, 2Cov(PC, CC) *≈* 0, and 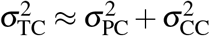. Significant differences in the distributions of precaudal and caudal variances thus indicate differential contributions to total vertebral count variation. Nonetheless, we also calculated the CVs for precaudal and caudal counts separately, since in most species the mean caudal count (*μ*_CC_) is greater than the mean precaudal count (*μ*_PC_), see Figure 2A.

### Investigating the evolution of intraspecific variation in vertebral counts

The CV is bounded at zero and sensitive to small means, therefore, the distribution of total count CV was right-skewed and not normally distributed (see Figure 4A), violating the assumption of normality required for PGLS models. In order to to examine whether intraspecific variation in total count could contribute to the evolution of the total vertebral count or had changed across the African phylogeny, we instead used a non-parametric approach. Since the range of vertebrae is 15 ( Supplementary Figure 1), we binned mean total vertebral counts into groups of 3 vertebrae to ensure that sample sizes per group were comparable. We applied a permutation-based implementation of the Kruskal-Wallis test to determine whether the medians were significantly different between groups. We simulated total count CV evolution using a single rate (*σ* ^2^) Brownian motion model, using the *fastBM* function in the R package *phytools* (v2.1.1) (Revell, 2024). The rate (*σ* ^2^) was set as the total count CV variance and CV estimates were bounded between 0 and 1 to avoid the estimation of negative CVs. Model fits were simulated 10,000 times, each of which was used as input for a Kruskal-Wallis test. *χ*^2^ values were extracted, and p-values were estimated from the null distribution using the formula: 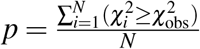, where *N* is the total number of simulations.

To assess whether somitic count variability has remained consistent across the phylogeny, we leveraged the unique evolutionary history of African cichlids. Multiple adaptive radiations are nested within Pseudocrenilabrinae, each having occurred independently (Ronco et al., 2021; Malinsky et al., 2018; Meier et al., 2017). These radiations, along with more basal riverine lineages, provide a natural experiment for investigating how trait variability evolves over time. Our previous analyses showed that interspecific variation in vertebral counts correlates with the age of a radiation, where total count evolution in each lacustrine radiation (and riverine lineages) was subject to its own stochastic rate of evolution (*σ* ^2^) (Bucklow et al., 2025). If intraspecific variation in vertebral counts evolved in tandem with shifts in vertebral count disparity, we might expect systematic differences in variability across these independent lineages. Alternatively, if somitic count variability has been maintained at consistent levels, despite shifts in mean vertebral counts, it would suggest that intraspecific variation has remained consistent throughout the diversification of African cichlids. Consequently, implying a developmental constraint, or canalisation, of the number of somite pairs formed during somitogenesis within a species across clades. To test for significant differences in the intraspecific variation between water systems, we redeployed the permutation-based Kruskal-Wallis test as described above.

As a second approach to investigate the evolution of intraspecific variation in vertebral counts, we quantified the phylogenetic signal in the total, precaudal and caudal count CVs using Pagel’s *λ* (Pagel, 1999) and Blomberg’s *K* (Blomberg et al., 2003). *These metrics assess the extent to which closely related species exhibit similar levels of intraspecific variability. If phylogenetic signal in the CV is weak or absent, it would suggest that intraspecific variation (i.e., somitic count variability) is similar between distantly related and closely related lineages, rather than being constrained by shared ancestry. Pagel’s λ* measures how much trait evolution follows a Brownian motion (BM) model by scaling internal branch lengths of the phylogenetic tree. A *λ* value of 1 indicates strong phylogenetic dependence, meaning that intraspecific variability evolves in proportion to shared ancestry. In contrast, a *λ* of 0 suggests no phylogenetic structure, implying that variability arises independently across species. Intermediate values indicate partial phylogenetic influence, where variation is shaped by both ancestry and other evolutionary factors. Blomberg’s *K* compares observed trait variation to expectations under BM. A *K* value of 0 suggests that the trait varies independently of phylogeny, whilst a value of 1 suggests that variation follows BM exactly. Values of *K <* 1 indicate that closely related species are more different than expected, suggesting weaker phylogenetic constraint. Conversely, *K >* 1 implies that related species are more similar than expected under BM, potentially due to stabilising selection or shared developmental constraints. Both metrics were estimated using the *phylogsig* function in *phytools* (v2.1.1) (Revell, 2024). Significance was assessed through 100,000 simulations. For *λ*, we examined its likelihood profile, while for *K*, we compared observed values to simulated null distributions (Supplementary Figure 2).

### Testing scaling of intraspecific variation and whole body elongation

We had previously reported that elongation of the fusiform body in African cichlids has partly been driven by the addition of vertebrae (Bucklow et al., 2025). In addition, prior evidence suggests that intraspecific variation in vertebral counts may enhance individual fitness by providing functional advantages (Swain, 1992; Tibblin et al., 2016). We therefore hypothesised that variation in vertebral count may scale with variation in body aspect ratio, and sought to examine whether this relationship is consistent both within and across African cichlids. To test this hypothesis, we fit within-species regressions (ordinary least squares) of ln[Total Vertebral Count] on ln[Length] – ln[Depth] (body aspect ratio) for species with at least 10 individuals (176 species). Individual species’ regression slopes (*β*_1_) were highly variable, generally weak, and noisy (see Results). Therefore, in order to assess the presence of an overall trend, while accounting for slope sampling variances and phylogenetic non-independence, we performed a multivariate phylogenetic meta-analysis (Lajeunesse, 2009) using restricted maximum likelihood (REML) via the *rma.mv* function in the R package *metafor* (v4.8.0) (Viechtbauer, 2010). Species-specific regression slopes were treated as effect sizes and weighted by their respective sampling variances (i.e. the squared standard errors). Species were included as a random effect, allowing each to deviate from the overall mean slope. To account for shared evolutionary history, we included a phylogenetic covariance matrix constructed using the *vcv* function from the R package *ape* (v5.7.1) (Paradis and Schliep, 2019). This approach allowed us to estimate the average relationship between vertebral number and body aspect ratio while accounting for both measurement uncertainty and phylogenetic structure.

## RESULTS

### Precaudal and caudal vertebral counts evolve independently across cichlid lineages

We found no significant correlation between ln[Precaudal Count] and ln[Caudal Count] across species (Figure 2A; *R*^2^ = 0.002, *β*_1_ = 0.082, *λ* = 0.935, *d.f.* = 427, *p* = 0.1884), suggesting that the number of vertebrae in each region has evolved largely independently. This indicates that lineages have gained or lost vertebrae in one domain without requiring corresponding changes in the other. We found a very weak but significant positive correlation between the relative proportions of precaudal and caudal vertebrae (ln[Precaudal] - ln[Caudal]) and total vertebral count, indicating that the addition of vertebrae may be driven primarily by the addition of precaudal vertebrae (Figure 2B; *R*^2^ = 0.02, *β*_1_ = 0.269, *λ* = 0.935, *p <* 0.001, *d.f.* = 427). However, this pattern appears to be influenced by specific genera, such as *Gobiocichla* and *Cyprichromis*, which exhibit relatively high total vertebral counts alongside an unusually large number of precaudal vertebrae (Oliver, 2024; Bucklow et al., 2025). Incorporating a quadratic term into the PGLS model significantly improved model fit (Δ*AIC >* 3). Only the quadratic term was significant (*β*_2_ = 0.460*, p <* 0.001), suggesting that the relationship between ln[Precaudal] - ln[Caudal] and ln[Total Count] is not constant but varies across the range of total vertebral counts, rather than following a simple power-law relationship. Consistent with previous findings (Bucklow et al., 2025), different lineages, therefore, have independently modified precaudal-to-caudal proportions and total vertebral counts. In addition, the phylogenetic signal for precaudal and caudal covariance was very high (*λ* = 0.935, 95% CI [0.910, 0.955]), indicating that while vertebral counts in each region evolve separately (Figure 2A), their variation remains strongly structured by shared ancestry. Consequently, similar vertebral patterns are found within closely related species, even though the exact number of vertebrae in each region can vary independently (Figure 2B).

### Somitic changes are usually accompanied by homeotic shifts in African cichlids

For most points of divergence in the phylogeny (i.e. nodes of the tree), changes in the total number of vertebrae are accompanied by shifts in regionalisation, suggesting a dominant role of combined somitic effects in the evolution of the vertebral column in African cichlids (Figures 2C, D points dispersed across space). Subsequently, changes to the total number of vertebrae (i.e. somitic changes) are rarely present without a concurrent shift in regionalisation. Nodes estimated to have undergone large somitic changes (and homeotic shifts) were predictably present in nodes representing divergences of clades with large differences in total vertebral count (Figure 3, black arrows). The highest rates of combined somitic effects were found exclusively in haplochromines (Figure 2D), specifically within the adaptive radiations of Lake Victoria and Lake Malawi (Supplementary Figure 3), which have undergone extremely rapid diversification in the presence of limited genetic variation (Meier et al., 2017; Malinsky et al., 2018). Therefore, evolutionary modification of vertebral column regionalisation (and co-occurring somitic changes) can occur with relatively little genetic variation and relatively quickly. We nonetheless did identify nodes where pure homeotic shifts (i.e., without co-occurring somitic changes) were predicted to have occurred (Figure 2C, D, blue triangles), indicating that changes in vertebral column regionalisation can occur independently of changes in vertebral count. Balanced somitic changes too were identified (Figures 2C, D, red squares), albeit at notably fewer nodes that either combined somitic effects or pure homeotic shifts (Figure 2E), suggesting that balanced somitic changes have played a minor role in the evolution of the vertebral column. Nonetheless, combined somitic effects, where changes in precaudal and caudal proportions have occurred as a consequence of additive or subtract somitic changes were identified at the vast majority of nodes in the phylogeny (Figure 2E), suggesting that the majority of changes in vertebral column regionalisation arise as a result of somitic, rather than homeotic changes.

**Figure 3.**
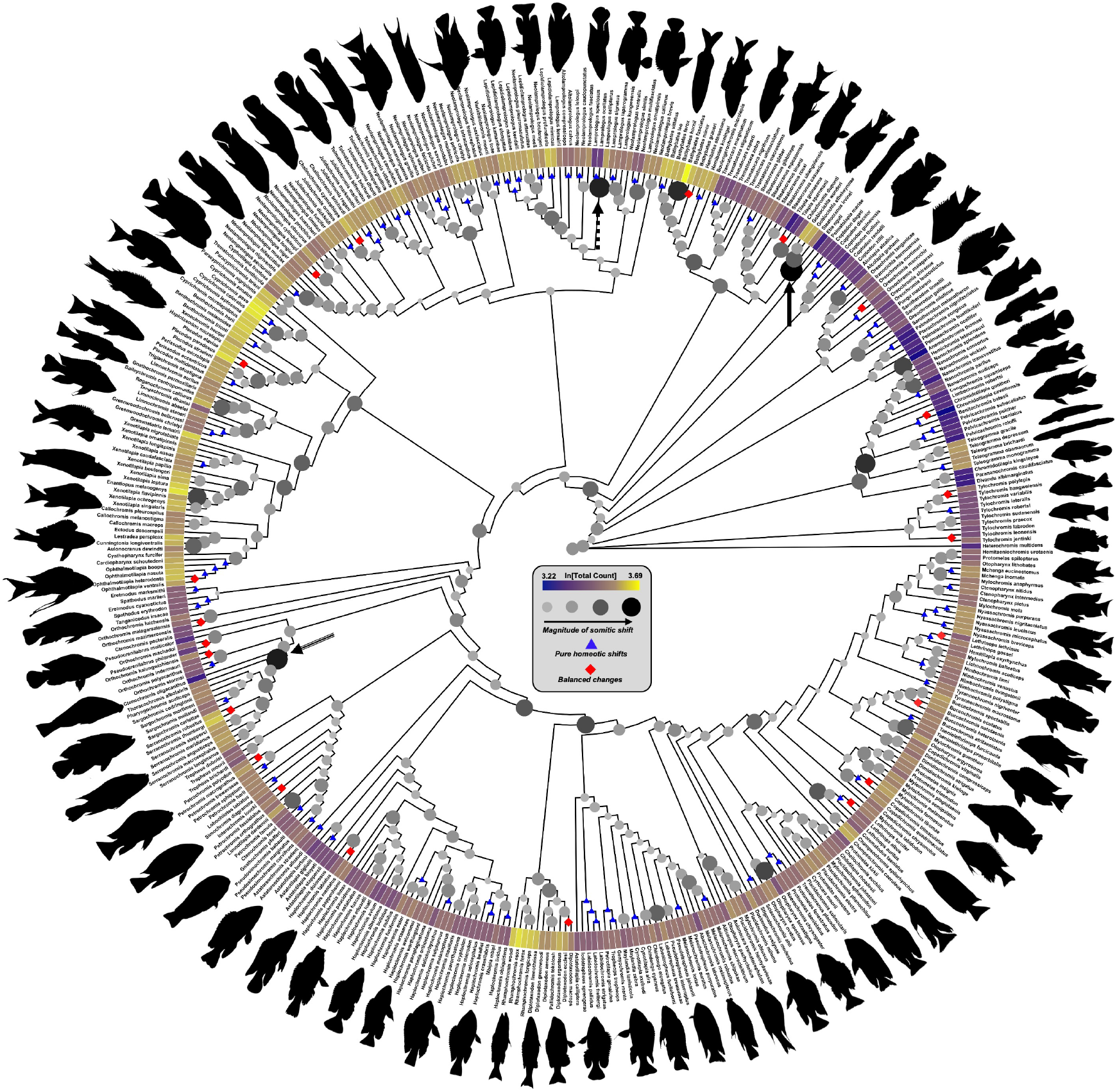
Magnitude of somitic changes in African cichlids. African cichlid cladogram. Node size and colour are scaled to the perpendicular distance (*d*) of nodes from the *y* = −*x* line (i.e. ‘pure homeotic shifts’, see Figure 2C), where larger, darker nodes represent the raw magnitude of estimated co-occurring somitic change (i.e. combined somitic effects). Blue triangles mark nodes with pure homeotic shifts and red diamonds where balanced somitic changes are predicted to have occurred. ln[Total Count] is plotted as a heatmap at the tips. Arrows match nodes exemplified in Figure 2C. Silhouettes of select cichlid species are displayed around the periphery of the phylogeny for visual reference. See Supplementary Materials for image credits.

### Intraspecific variation is low and lacks phylogenetic structure

Intraspecific variation in vertebral counts is tightly clustered around the mean for each species (Figure 4A). The median coefficient of variation (CV) is just 1.49%, indicating that for most species, the standard deviations are a small fraction of the mean total vertebral count. This suggests that vertebral counts are dispersed closely around the mean, with intraspecific variation in vertebral counts deviating minimally from the mean count. In real terms, the modal range of intraspecific counts is just *TC±* 1. Caudal variance (*σ* ^2^) is significantly higher than precaudal variance (Figure 4B, Wilcoxon Rank Sum, V = 5805.5, *p <* 0.001), suggesting that the loss or gain of caudal vertebrae is more common than that of precaudal vertebrae within species. Moreover, since the total count variance is the sum of the variances of the precaudal and caudal counts (see Methodology), caudal variance may contribute more to total vertebral count intraspecific variation than precaudal. However, we found no significant difference between the CVs for precaudal and caudal vertebral counts (Figure 4B, Wilcoxon Rank Sum, V = 16506, *p* = 0.196). Whilst caudal counts exhibit a larger absolute spread, likely a result of caudal counts generally being larger in African cichlids (see Figure 2B), the relative variation (compared to the mean vertebral count) is similar for both domains. Therefore, intraspecific variation in total vertebral count likely reflects both homeotic (shifts between regions) and somitic (additive or subtractive) effects. Estimates of phylogenetic signal revealed *K* values indistinguishable from zero (precaudal CV: *K* = 0.0089, *p* = 0.49; caudal CV: *K* = 0.0116, *p* = 0.40), and although *λ* values were significantly greater than zero (precaudal CV: *λ* = 0.166, *p* = 0.01; caudal CV: *λ* = 0.525, *p* = 0.0004), the flat likelihood profiles (see Supplementary Figure 2) suggest these results reflect estimation uncertainty rather than meaningful signal. Thus, both *K* and *λ* indicate little to no phylogenetic structure in the relative variation of vertebral counts, suggesting that intraspecific variation in precaudal and caudal counts is largely independent of phylogenetic relatedness and is unlikely to be constrained by shared ancestry.

**Figure 4.**
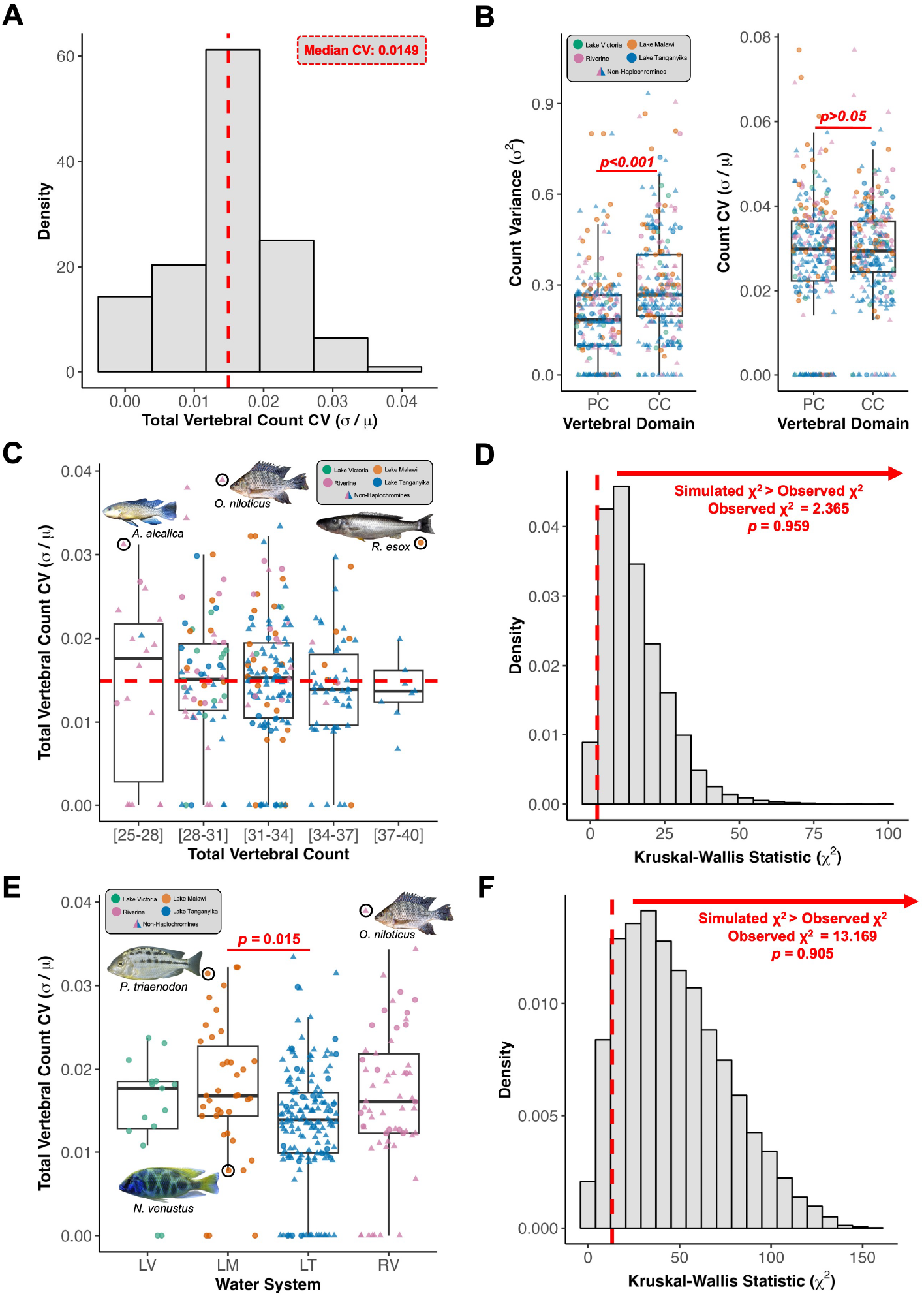
Intraspecific variation in vertebral counts is low. (A) Distribution of the coefficient of variation (CV; *σ/μ*) for total vertebral count across species. The median is indicated with a dashed red line. (B) Box plots of count variance (*σ* ^2^) for precaudal (PC) and caudal (CC) vertebrae (left), and corresponding CV values (right). Group medians are indicated by thick black lines. Differences in variance and CV were tested using a Wilcoxon rank-sum test; significance levels are indicated. Box plot of total count CV grouped by total count (C) and by water system (E), respectively. Overall median CV is indicated with a dashed red line. Permutation-based phylogenetic Kruskal-Wallis null distribution for total count CV by total count (D) and by water system (F), respectively. The observed *χ*^2^ value from the non-phylogenetically corrected test is indicated by a dashed red line. The *p*-value represents the proportion of permuted *χ*^2^ values greater than the observed value (see Methodology). Genera abbreviations: *A, Alcolapia*; *N, Nimbochromis*; *O, Oreochromis*; *P, Protomelas*; *R, Rhamphochromis*. Image credits provided in Supplementary Materials.

### The evolution of vertebral counts is not correlated with changes in intraspecific variation

We found no significant differences between the total count CV and the total vertebral count across any of the tested bins (Figure 4C, Kruskal-Wallis, *χ*^2^ = 2.366, *d.f.* = 4, p = 0.669). Results were consistent with a permutation-based Kruskal-Wallis test to correct for phylogenetic relationships (Figure 4D, n=10,000, p = 0.959), suggesting that accounting for phylogenetic relatedness did not substantially alter any of the group differences. Therefore, somitic count variability does not scale with total vertebral count and is unlikely to have undergone positive or indeed purifying (negative) selection. As a consequence, elongate lineages do not differ in somitic count variability from lineages with more conservative vertebral counts. We did find evidence of difference in the total count CV between water systems (Kruskal-Wallis, *χ*^2^ = 13.17, *d.f.* = 3, p = 0.004), but pairwise comparisons indicated a significant difference only between Lake Tanganyika and Lake Malawi (*p* = 0.015), see Figure 4E. However, a phylogenetic implementation of the Kruskal-Wallis test (see Methodology) found no significant differences in the median CV between any of the respective water systems (Figure 4F, p = 0.912). Consequently, intraspecific variation has remained stable across the phylogeny despite differences in evolutionary dynamics of interspecific total count evolution between water systems (Bucklow et al., 2025). Consistent with this, we found no evidence of phylogenetic signal in the total count CV. Both *λ* (0.196, p = 0.197) and *K* (0.030, p = 0.077) were indistinguishable from 0 (Supplementary Figure 2), suggesting that interspecific variation in intraspecific variability is not strongly structured by phylogenetic relatedness. Therefore, although vertebral counts have evolved extensively, somitic count variability has likely remained consistent across the phylogeny. This suggests that low intraspecific variation in vertebral number results from a conserved developmental constraint that has persisted throughout the diversification of African cichlids.

### Intraspecific variation does not scale predictably with body elongation

Despite the importance of vertebral addition in driving body elongation between species (Bucklow et al., 2025), we found no evidence that individuals with increased vertebral counts exhibit higher body aspect ratios across the full range of body shapes present in African cichlids (Figure 5A). This lack of association held both across species and within them. Within-species regressions of ln[Total Count] on ln[Length] – ln[Depth] (body aspect ratio), produced highly variable and generally weak slopes, with no consistent pattern in direction or magnitude (Figure 5B). The REML estimated mean slope was 0.448 *±* 0.401 (z = 1.116, *p* = 0.264; 95% CI: –0.338 to 1.234), indicating no significant association between vertebral count and elongation across species (Figure 5C). Despite this, heterogeneity was substantial (Q(175) = 338.35, *p <* 0.0001), and the estimated between-species variance was high (*σ* ^2^ = 0.811), suggesting that species differ markedly in the strength and direction of this relationship (including decreases in body aspect ratio being associated with vertebral count increases). Crucially, there is no consistent evidence that individuals with more vertebrae also tend to be more elongate. Therefore, intraspecific variation in vertebral number does not scale predictably with body aspect ratio, and vertebral count variation appears decoupled from large-scale morphological divergence in elongation across species.

**Figure 5.**
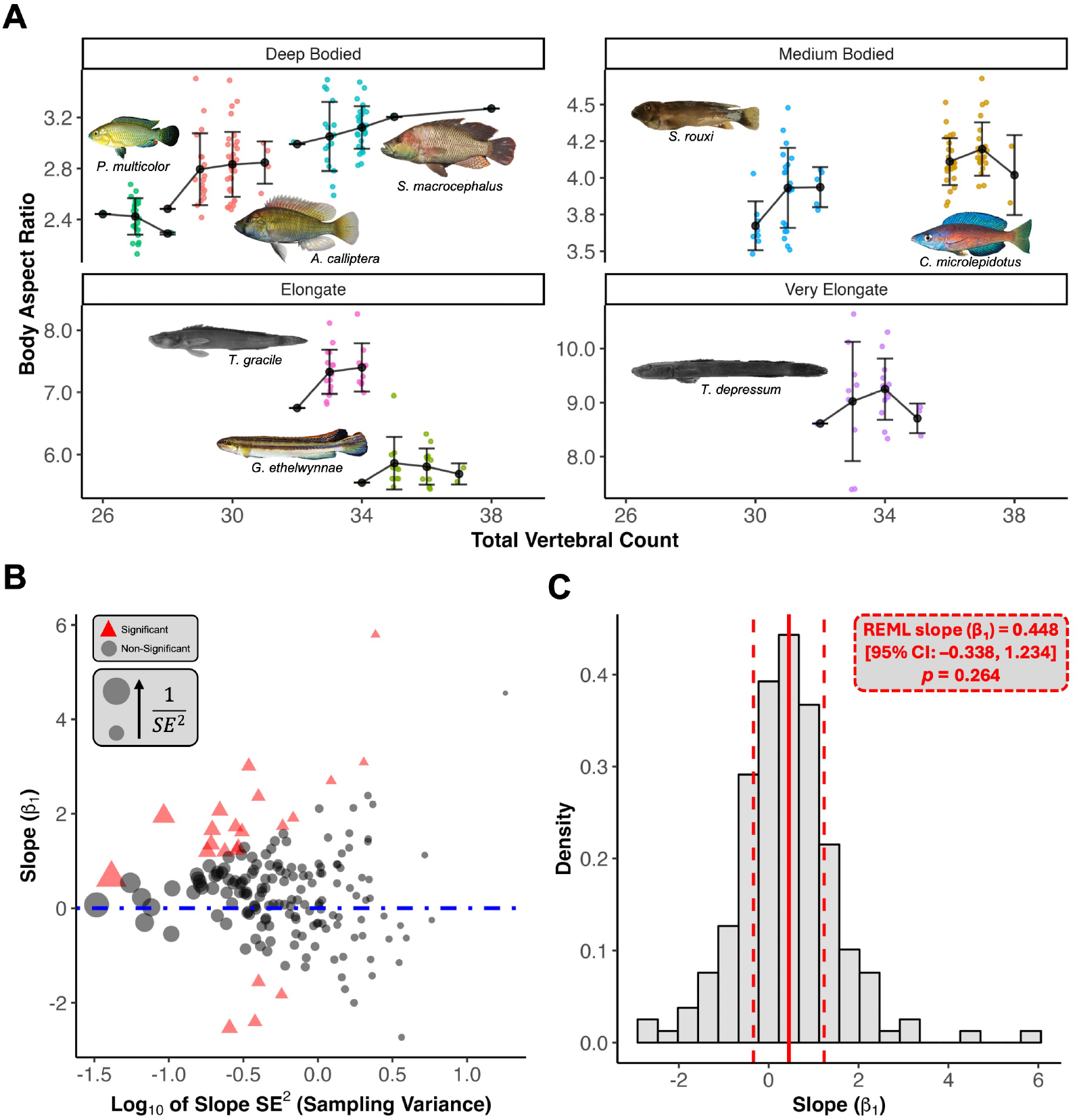
Intraspecific variation does not scale predictably with body elongation. (A) Body aspect ratios plotted against vertebral count for selected species with more than 25 individuals. Panels are faceted by elongation category. Black circles represent the mean body aspect ratio for each vertebral count, with vertical bars indicating the standard deviation. Species images are included for reference. (B) Scatterplot of intraspecific regression slopes (*β*_1_) of ln[Total Vertebral Count] on ln[Length]-ln[Depth] against their respective sampling variances. Point sizes are scaled according to the weights used in the meta-analysis (i.e., inverse of the squared standard error); larger points reflect more precise estimates and thus greater weight. Red triangles indicate slopes significantly different from zero (p*<*0.05). A dot-dashed blue line indicates a slope of zero (i.e., no relationship between ln[Total Count] and ln[Length]-ln[Depth]). (C) Histogram showing the distribution of intraspecific slopes across all species (n=177) with at least ten individuals. The solid red line shows the restricted maximum likelihood (REML) estimate of the mean slope; dashed red lines denote the 95% confidence interval. Note that the confidence interval overlaps zero. Genera abbreviations: *A, Astatotilapia*; *C, Cyprichromis*; *G, Gobiocichla*; *P, Pseudocrenilabrus*; *S, Serranochromis* (*macrocephalus*), *Steatocranus* (*rouxi*); *T, Teleogramma*. Image credits provided in Supplementary Materials.

## DISCUSSION

Modification of the regionalisation of the vertebral column is common in teleosts (Ward and Brainerd, 2007; Ward and Kley, 2012), including in African cichlids (Bucklow et al., 2025). Our results further support that teleostean clades have independently modified the relative proportions of precaudal and caudal vertebrae (Ward and Brainerd, 2007; Mehta et al., 2010). In addition, our analysis suggests that homeotic shifts can, and likely have, occurred during the diversification of African cichlids, helping to explain some of the variation in vertebral proportions observed across the subfamily. The presence of pure homeotic shifts suggests that AP patterning can be modified without altering somitogenesis. Homeotic transformations are well documented in mutational studies across vertebrates but it is less clear whether these mutations could be maintained across populations (and species). In tetrapods, constraints on vertebral column regionalisation are well documented. Famously, mammals are constrained to have exactly seven cervical vertebrae in developing embryos (Böhmer et al., 2018; Buchholtz and Stepien, 2009), albeit complicated by the early ossification of ‘cervical’ (thoracic) vertebrae posterior to the seventh cervical vertebrae in sloths (Asher et al., 2011). Cervical vertebral counts in archosaurs, in contrast, are highly variable (Marek et al., 2021). Furthermore, thoracic-to-lumbar transformations are common in mammals (Narita and Kuratani, 2005; Cerbus et al., 2024), indeed homeotic transformations within all domains posterior of the cervical domain can describe much of the vertebral formulae variation in mammals (Cerbus et al., 2024).

Pure homeotic shifts could be driven by changes in the regulation of the *hox* genes responsible for patterning the precaudal-caudal boundary. However, elucidating anterior-posterior patterning during teleostean development is particularly difficult due to the presence of seven/eight *hox* clusters (Crow et al., 2006; Hoegg et al., 2007), where overlapping expression of many paralogous genes, often with functional redundancy (Adachi et al., 2024), makes identifying the key *hox* genes difficult. *hox6* paralogs have been implicated in determining precaudal vertebral identity in *Danio rerio* (zebrafish) and the anterior-expression limits of *hoxC10a* and *hoxD12a* correlate with the somites that go on to contribute to vertebrae at the precaudal-caudal boundary (Morin-Kensicki et al., 2002; Hayward et al., 2015). There is little reason, however, to assume that the specific *hox* genes involved are conserved across teleosts. Subfunctionalisation of *hox* rhombomere patterning has occurred multiple times in teleosts (Scemama et al., 2006), including in cichlids (Le Pabic et al., 2007). Moreover, the identity of *hox* genes which specify fin formation (Sordino et al., 1995) are also not conserved in teleosts (Adachi et al., 2024). Epigenetic regulation of *hox* expression may also play a role in teleosts. Dysregulation of *hox* methylation due to the loss of maternally-expressed factors has been shown to lead to homeotic transformations in zebrafish (Xue et al., 2022). Even in tetrapods, the *Hox* code does not appear to be particularly conserved; paralogous groups 9–11 have been implicated in patterning transitions between different axial domains along the anterior-posterior axis (Cerbus et al., 2024). Therefore, the key genes involved in patterning the precaudal-caudal boundary could vary amongst different teleostean lineages. A systematic, comparative investigation of *hox* expression across multiple teleost clades will be essential to clarify how these regional boundaries are established and how they have evolved.

We have considered homeotic transformations in just two domains but previous studies have suggested that the teleostean vertebral column has more than two distinct regions (De Clercq et al., 2017; Jawad et al., 2018), however, unlike in terrestrial tetrapods these regions are difficult to visually distinguish due to the relatively homogenous vertebral shape present along the anterior-posterior axis (De Clercq et al., 2017), prohibiting large scale comparative studies such as those that can be more easily conducted in tetrapods (Carapuço et al., 2005; Asher et al., 2011; Cerbus et al., 2024). Constraints placed on the structure of the vertebral column due to a fully aquatic environment may be responsible for this homogenisation. Skates and cetaceans both have what appear to be relatively homogenised vertebral columns. However, landmarked-based geometric morphometric quantification of vertebral shape variation demonstrates the presence of multiple domains nested within the precaudal and caudal domains, with the number of regions being comparable to tetrapods (Criswell et al., 2021; Gillet et al., 2024). Systematic quantification of vertebral shape within teleosts along the AP axis is likely to identify the presence of multiple, nested domains within the precaudal and caudal domains. Once these nested domains have been defined, counting the number of vertebrae within each domain could provide novel insights into the evolution of vertebral column regionalisation, such as the non-homeotic positive correlation recently reported in vertebral counts between the distal cervical and sacral domains in birds (Cerbus et al., 2024).

The dominance of combined somitic effects suggests that much of the variation in the African cichlid vertebral column can be attributed to changes in somite number. Importantly, this also implies that most shifts in vertebral proportions arise as a by-product of somitic changes, rather than through direct modification of axial patterning mechanisms like the *Hox* genes. In other words, homeotic transformations could occur without requiring evolutionary changes in the *Hox* regulatory network itself. In teleosts, anterior *Hox* genes are expressed even prior to gastrulation (Stevens et al., 1996; Amali et al., 2013). While they are not essential for initiating gastrulation, these genes do help organise mesodermal derivatives during development (Nowicki and Burke, 2000; Iimura and Pourquié, 2007). Later-acting *Hox13* genes have also been implicated in regulating the neuromesodermal progenitor (NMP) niche (Ye and Kimelman, 2020), as well as controlling mesodermal cell movements out of the tailbud in tetrapods (Denans et al., 2015). However, if the rate of somite formation changes, producing more or fewer somites in the same amount of time, the overall *Hox* ‘timer’ may remain unchanged. In this scenario, the *hox* pattern shifts as a consequence of changing the number of somites. In Lake Malawi cichlids, an increased somitic rate, relative to *Astatotilapia calliptera* which forms fewer somites, has been shown to contribute towards the increased somite counts observed in *Rhamphochromis sp.* ‘Chilingali’ (Marconi et al., 2023). Thus, altering the rate or duration of somitogenesis may be sufficient to drive both changes in total vertebral count and regionalisation, without necessitating changes to the underlying *Hox* code. This could also explain why balanced somitic changes where both the somitic rate and *Hox* timer evolve in tandem to preserve proportions appear to be relatively rare: such changes would require coincident modifications affecting both somitogenesis and *Hox* regulation, which appears to be evolutionarily unlikely.

Given the increased rates observed in haplochromines, very little genetic variation appears to be required in order to drive these somitic changes (and indeed homeotic changes), reinforcing the utility of haplochromines as powerful models in evolutionary developmental biology (Santos et al., 2023). Lake Malawi cichlids, for example, are extremely closely related, more so than human populations (Malinsky et al., 2018). CTCCC-binding factor (CTCF) has been implicated in the regulation of *hox* timing. Genomic CTCF-binding sites act as anchor points for the formation of chromatin loops, which can alter the rate of *hox* transcription (Deschamps and Duboule, 2017; Rekaik et al., 2023). Modification of CTCF-binding sites has even been shown to trigger homeotic transformations (Narendra et al., 2016; Rekaik et al., 2023). Therefore, the evolutionary modification of CTCF-binding sites in the *hox* clusters could act as a mechanism to modulate the *Hox* ‘timer’. Therefore, relatively small genetic mutations between closely related animals could alter CTCF-binding sites in the *Hox* clusters. Larger intragenic distances (proxy for number of CTCF-binding sites) between *Hox* clusters do not appear to correlate with shifts in regionalisation across tetrapods, at least when corrected for phylogeny. However, significant clade specific differences were identified, such as a positive correlation between the number of thoracic vertebrae and HoxD1-HoxD3 intergenic distances in amphibians (Cerbus et al., 2024). Therefore, smaller scale effects may be possible. Many axial phenotypes have now been characterised in haplochromine cichlids (Oliver, 2024; Bucklow et al., 2025) and we now have robust whole genomes for many haplochromines (Meier et al., 2017; Malinsky et al., 2018; Masonick et al., 2022). An analysis of *Hox* intergenic distances and their correlation with multiple axial phenotypes in haplochromines could provide a powerful model system to investigate this further.

Strong developmental constraints are likely maintaining vertebral number within a narrow range, preventing intraspecific variation in vertebral counts to fluctuate within a species. This stability across the phylogeny implies that selection or genetic/developmental constraints prevent vertebral count from varying significantly within species, suggesting strong canalisation of intraspecific variation in the number of somitic pairs formed during somitogenesis. Intraspecific variation in vertebral counts has been identified across vertebrates (Yamahira et al., 2006; Slijepčević et al., 2015; Tibblin et al., 2016), including in humans (Hu et al., 2016; Yan et al., 2022). Both common garden experiments, as well as heritability estimates derived from quantitative genetic studies, have shown that intraspecific variation in vertebral counts has an additive and heritable genetic basis (Leary et al., 1985; Manier et al., 2007; Alho et al., 2011). This raises the question of why somitogenesis is canalised so strongly. At least in teleosts, it may partly be a consequence of needing to buffer against environmentally-induced phenotypic plasticity. Changes in salinity, light, and temperature have all been shown to lead to changes in vertebral counts (Fowler, 1970; Tibblin et al., 2016; Campbell et al., 2021). Therefore, canalisation of the intraspecific somitic count variability would minimise the production of maladaptive phenotypes, particularly in ecological contexts where axial morphology may be linked to locomotory performance, predator avoidance, or habitat use (Swain, 1992; Tibblin et al., 2016).

It was nonetheless surprising that we found no relationship between intraspecific variation in total vertebral counts and body aspect ratio, given the well-established role of vertebral addition in driving interspecific body elongation, across teleosts (Ward and Brainerd, 2007; Ward and Mehta, 2010; Mehta et al., 2010), including in African cichlids (Bucklow et al., 2025). If intraspecific variation is indeed important in generating intraspecific differences in phenotypes (Swain, 1992; Tibblin et al., 2016), it does not seem to affect elongation of the body on the level of individuals. In cichlids, body shape is known to be highly polygenic, with quantitative trait locus (QTL) analyses of hybrids revealing contributions from many loci (Husemann et al., 2017; DeLorenzo et al., 2023). Indeed, it appears that hybridisation is sufficient to lead to the emergence of transgressive body shapes (Husemann et al., 2017; Selz and Seehausen, 2019), including vertebral count in other teleostean lineages (Seleit et al., 2024). Many of these loci are likely involved in multiple developmental pathways throughout growth and development. In Central American cichlids (*Amphilophus* spp.), vertebral counts and lateral line scale numbers not only covary phenotypically but also co-segregate genomically, suggesting shared developmental control and tight linkage of the underlying genomic regions (Ehemann et al., 2024). In African cichlids, we have previously shown that evolutionary changes in posterior body elongation are tightly coupled to changes in cranial elongation. However, cranial elongation appears to occur without corresponding increases in vertebral number (Bucklow et al., 2025). Therefore, body elongation in cichlids could arise through the reuse or modification of developmental programs other than those controlling axial segmentation. QTL analysis focused specifically on identifying genomic and phenotypic covariation in interspecific hybrids differing in multiple axial phenotypes, including total numbers of vertebrae, proportions of precaudal and caudal vertebrae and body shape may help further elucidate the complex relationship between the axial skeleton and body shape evolution.

In summary, our findings highlight the dynamic interplay between homeotic transformations and somitic changes in shaping the vertebral column of African cichlids. While shifts in vertebral regionalisation can arise through modification of anterior-posterior patterning, our results suggest that much of the observed variation is likely driven by changes in somitogenesis, with homeotic effects emerging as a by-product rather than a direct target of selection. The developmental processes governing somite formation and regionalisation in cichlids are likely highly canalised, limiting intraspecific variation even as interspecific diversity evolves. Future work integrating comparative developmental genetics, quantitative morphology, and genomics, particularly leveraging the exceptional diversity and genomic resources of haplochromine cichlids, holds considerable promise for disentangling the relative roles of somitogenesis and homeotic regulation in vertebral evolution. Ultimately, understanding how these processes interact across clades will shed new light on the origins of vertebral regionalisation and its evolutionary flexibility in vertebrates.

## SUPPLEMENTARY FIGURES

**Supplementary Figure 1.**
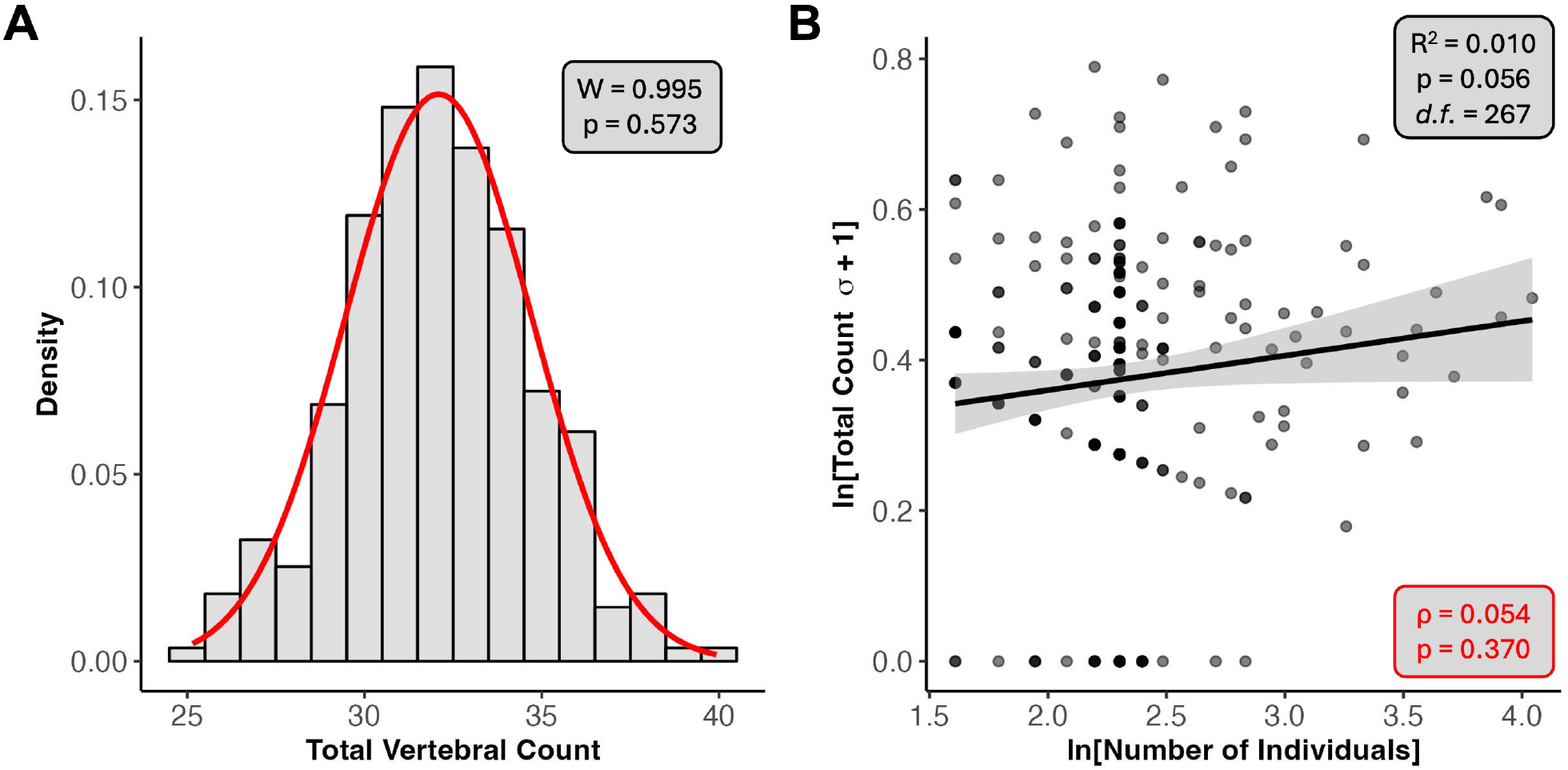
(A) Distribution of total vertebral counts for species represented by *≥* 5 individuals and included in the phylogeny (n = 277). The observed range (25–40 vertebrae) is consistent with previous reports for African cichlids (Oliver, 2024; Bucklow et al., 2025). A red curve shows an idealised normal distribution. Results of a Shapiro-Wilk test for normality are provided in the black box. (B) Relationship between ln-transformed total vertebral count standard deviation (*σ*) and ln-transformed sample size. To accommodate zero values, one was added to all standard deviations prior to transformation. The black line shows the ordinary least squares (OLS) regression fit, with shaded regions indicating standard error. Results of a Spearman rank correlation are shown in red.

**Supplementary Figure 2.**
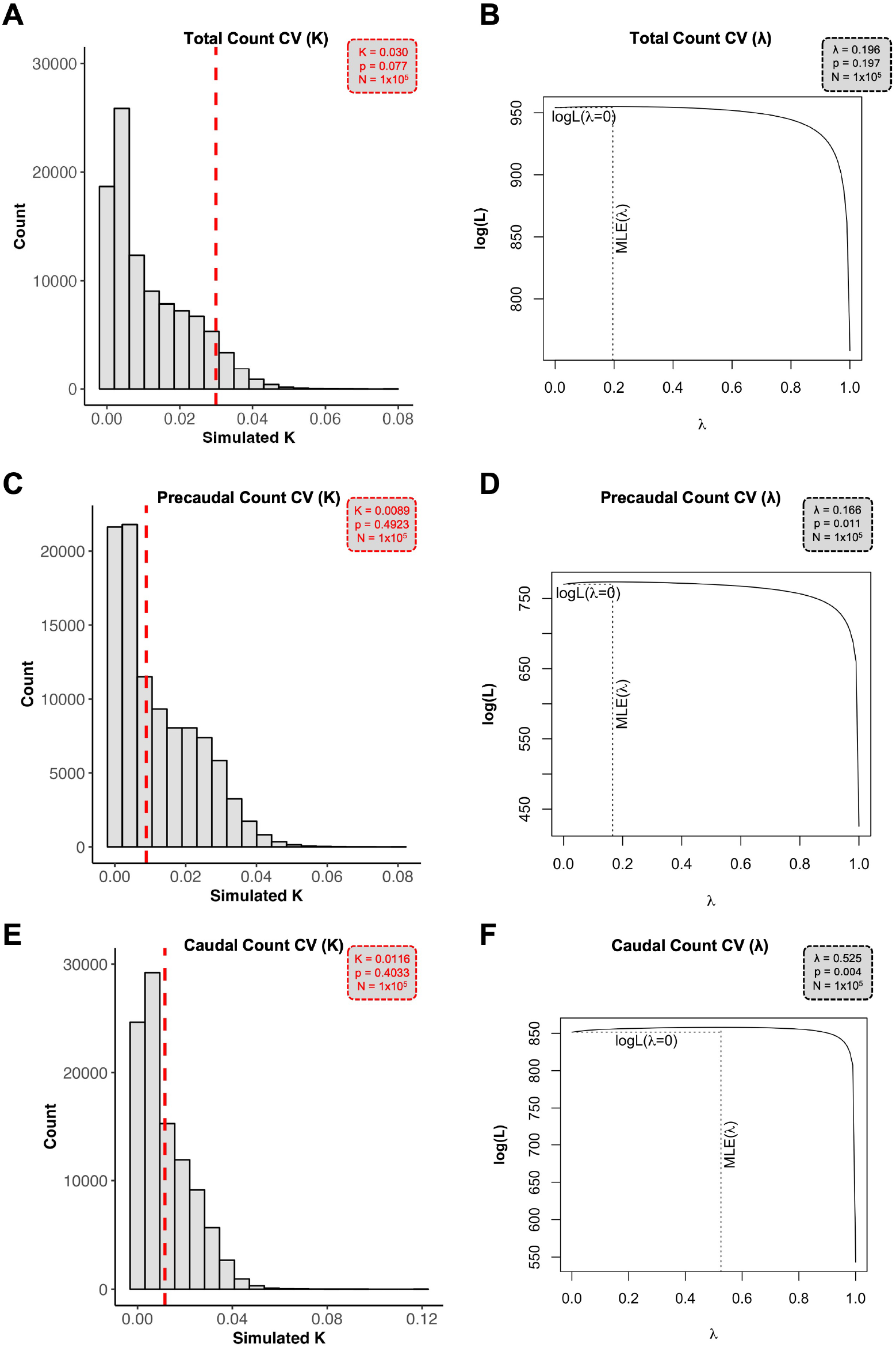
Null distributions of Blomberg’s *K* estimates (left) and likelihood profiles of Pagel’s *λ* estimates (right) for (A, B) total count CV, (C, D) precaudal count CV, and (E, F) caudal count CV. Observed *K* values are shown as red dashed lines. The maximum likelihood estimate (MLE) for *λ* is shown as a black dotted line. Results of hypothesis tests for *H*_0_ : *K, λ >* 0 are shown in red (*K*) and black (*λ*), respectively. The number of simulations is indicated in each panel. Note that although significant p-values were obtained for *λ >* 0 in precaudal and caudal CV, the likelihood profiles are very flat, suggesting these may represent false positives.

**Supplementary Figure 3.**
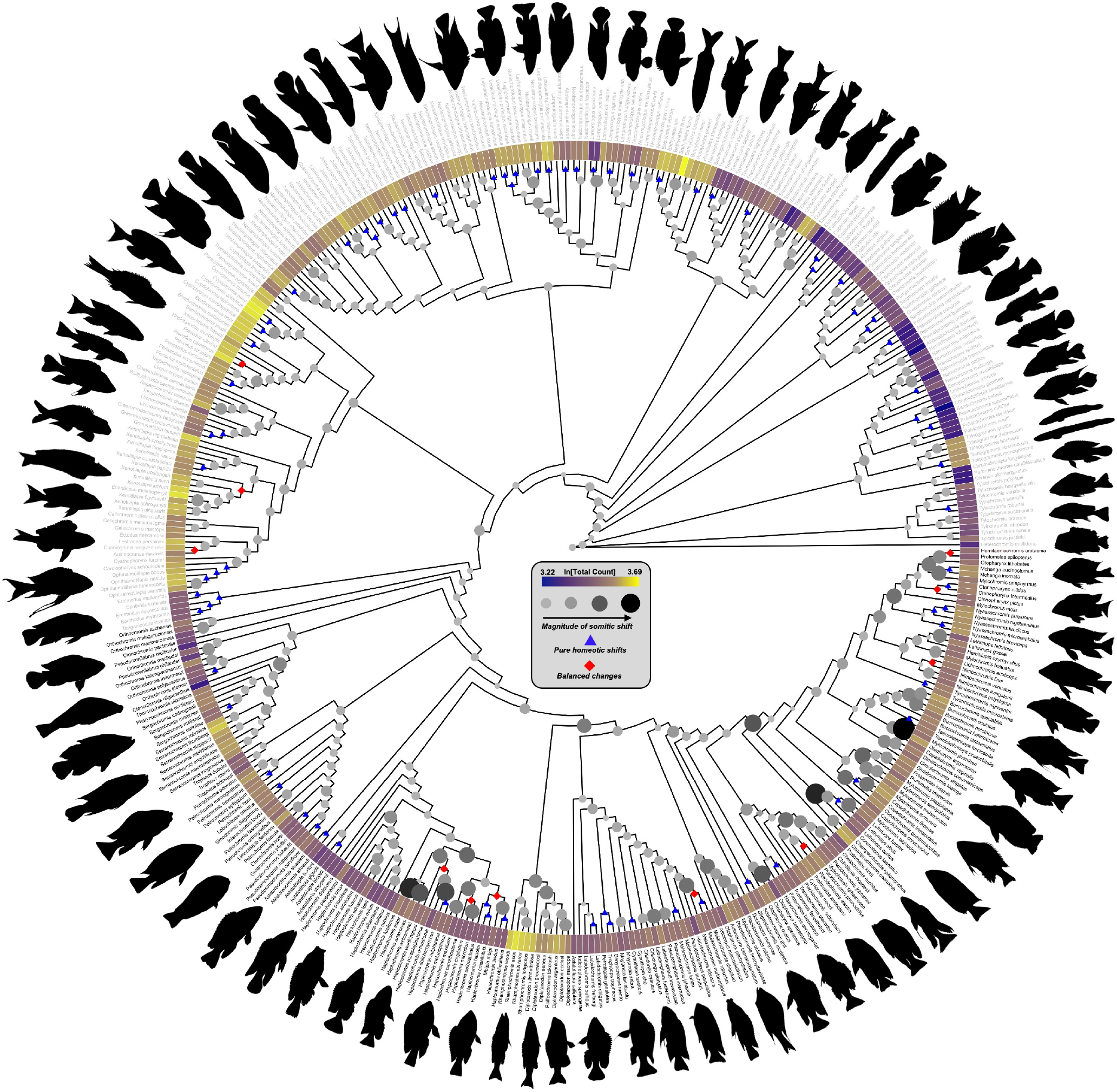
All branch lengths are fixed at 1 for visualisation purposes, as many internal branches are too short to display clearly alongside node labels. However, perpendicular distances were derived from phylogenetic independent contrasts (PICs) calculated using the original, time-calibrated branch lengths (see Methodology). Tip labels in black correspond to haplochromine species. Larger, darker internal nodes indicate elevated rates of co-occurring somitic change (i.e., combined precaudal and caudal effects). Nodes marked with blue triangles indicate inferred pure homeotic shifts, while red diamonds mark nodes where balanced somitic changes are predicted. ln[Total Count] (log-transformed total vertebral count) is plotted as a heatmap at the tips of the tree. Silhouettes of select cichlid species are displayed around the periphery of the phylogeny for visual reference. See Supplementary Materials for image credits.

## AUTHOR CONTRIBUTIONS

C.V.B, B.V. and R.B. conceived the study. E.D.R helped with phylogenetic generalised least squares analysis of precaudal and caudal counts. C.V.B wrote the manuscript. B.V. and R.B. edited the manuscript. All authors reviewed the manuscript.

## ACKNOWLEDGMENTS

This research was funded by a Biotechnology and Biological Sciences Research Council (BBSRC) studentship (Grant Number: 2445747). We thank Laura Soul for her original work describing the methodology to identify homeotic and somitic effects on phylogenetic scales. We thank James Hammond for helpful discussion regarding the evolvability of somitogenesis. Thank you to the fishes whose lives were sacrificed for this work.

## Notes

### Competing Interest Statement

The authors have declared no competing interest.

