## Supporting Code and Data for "Somitic Change Drives Changes in Vertebral Regionalisation in African Cichlids Despite Strong Canalisation of Somite Number": Somitogenesis and Regionalisation.rmd

````
---
title: "Somitogenesis and Regionalisation"
author: "Callum Bucklow"
date: "`r Sys.Date()`"
output: html_document
---

```{r Setup, include=FALSE}
knitr::opts_chunk$set(message = FALSE) ##Remove messages for all chunks
knitr::opts_chunk$set(warning = FALSE) ##Remove warnings for all chunks
```

```{r Libraries}
# Load the necessary libraries
library(tidyverse) #data wrangling
library(tidyr) #data wrangling
library(dplyr) #data wrangling
library(ggplot2) #for plotting when I don't want to use Base R (i.e. most of the time)
library(gridExtra) #for ggplot, to plot multiple plots together
library(ape) #for phylogeny manipulation and all the rest
library(phytools) #for various things, including phylogenetic PCA, PGLM etc.
library(caper) #for pgls and will report pseudo-r2 for these models
library(metafor) #for multiple
library(viridis)
library(ggtree) #hate using this but useful for plotting cladograms 
```

```{r Import the Data}
all_species <- readRDS('Data/all_species.rds') #for tree
node_map <- read.csv('Data/node_map_somitogenesis_regionalisation.csv')
vertebral_counts <- read.csv('Data/vertebral_counts_master.csv')
```

```{r Plot Tree with Node Labels}
pdf("Figures/Somitogenesis and Regionalisation/tree_for_map.pdf", width = 8.4, height = 11)

plot(all_species$tree,
     type="fan",
     cex=0.15,
     label.offset = 0.5)
nodelabels(cex=0.15, frame = "n", col = "red")

dev.off()
```

```{r Data Import and Wrangling}

#import all_species data
all_species <- readRDS('Data/all_species.rds')

#store in dataframe for analysis
all_species_data <- as.data.frame(cbind(all_species$matrix_data, "species_name" = all_species$stored_data$species_name))

#all_species_data$relative_posterior_elongation <- all_species_data$ln_ant_length_ln_width - all_species_data$ln_post_length_ln_width

#reformat data:
columns_not_numeric <- c("species_name")

#convert all columns except those in `columns_not_numeric` to numeric
all_species_data <- all_species_data %>%
  mutate(across(-all_of(columns_not_numeric), as.numeric))

#recombine with stored qualitative data
all_species_data <- left_join(all_species$stored_data, all_species_data, by='species_name')
```

```{r Construct VCV Matrix for Downstream Analysis}
#construct the vcv matrix:
aspect_total <- caper::comparative.data(all_species$tree, all_species_data, species_name, vcv=TRUE, vcv.dim=3)
```

```{r Fit PGLS model for ln[Precaudal] ~ ln[Caudal]}

#ols.precaudal.caudal <- lm(data=all_species_data, precaudal_count ~ caudal_count)
#summary(ols.precaudal.caudal)

#pgls.precaudal.caudal.lambda <- 
  #caper::pgls(precaudal_count ~ caudal_count, aspect_total, lambda = 'ML')
#summary(pgls.precaudal.caudal.lambda)
```

```{r Plot Precaudal versus Caudal Counts}

#colours taken from Wong colour blind palette
water_system_colours <- c("Lake Victoria" = "#43AA8B", 
                          "Lake Malawi" = "#D55E00", 
                          "Lake Tanganyika" = "#0072B2", 
                          "Riverine" = "#CC79A7")

c1 <- ggplot(all_species_data, aes(x=caudal_count, y=precaudal_count,  colour=water_system, shape=haplochromine))+ #, label=species_name))+
  geom_point(size = 1.50, alpha=0.75)+
  #geom_text(vjust=-0.5, hjust=0.5, size=1)+ #add species labels
  geom_abline(intercept = 2.25248, 
              slope = 0.13692, 
              size = 0.75, colour = 'black', linetype = 'solid')+ #ols is black
  geom_abline(intercept = 2.376684, 
              slope = 0.082359, 
              size = 0.75, colour = 'red', linetype = 'dashed')+#pgls is red
  scale_colour_manual(values = water_system_colours)+
  scale_shape_manual(values = c('Haplochromine' = 19, 'Other' = 17))+
  xlab("ln[Caudal Count]")+
  ylab("ln[Precaudal Count]")+
  theme_classic(base_size = 12)+
  theme(legend.position='none',
                 axis.text = element_text(size=11),
                 axis.title = element_text(size=11, face='bold'))
c1
ggsave("Figures/Somitogenesis and Regionalisation/precaudal_caudal_pgls.jpeg", c1, dpi = 600, width = 10, height = 10, units = "cm")
```

```{r Fit PGLS model for ln[Precaudal] - ln[Caudal] ~ ln[Total Count]}
ols.total.pc.c <- lm(data=all_species_data, ln_pc_ln_c ~ total_count)
summary(ols.total.pc.c)

pgls.total.pc.c.lambda <- 
  caper::pgls(ln_pc_ln_c ~ total_count, aspect_total, lambda = 'ML')
  summary(pgls.total.pc.c.lambda) #-871.8973

pgls.total.quad.pc.c.lambda  <- pgls(ln_pc_ln_c ~ poly(total_count, 2), aspect_total, lambda='ML')
summary(pgls.total.quad.pc.c.lambda) #-881.0456
```

```{r Plot Total Versus Precaudal:Caudal Ratio}
#generate random x (total count) values
prediction_data <- data.frame(total_count = seq(min(all_species_data$total_count), 
                                                max(all_species_data$total_count), 
                                                length.out = 100))
#use the PGLS model to predict ln[pc]-ln[c] (y) values from the quadratic pgls model:
prediction_data$predicted_ln_pc_ln_c <- predict(
pgls.total.quad.pc.c.lambda, newdata = prediction_data)

#plot the scatterplot of total count and ln[pc]-ln[c]
c2 <- ggplot(all_species_data, aes(x=total_count, y=ln_pc_ln_c, colour=water_system, shape=haplochromine))+
  geom_point(size = 1.50, alpha=0.75)+
  scale_colour_manual(values = water_system_colours)+
  scale_shape_manual(values = c('Haplochromine' = 19, 'Other' = 17))+
  geom_abline(intercept = 0.84143, 
              slope = -0.30584, 
              size = 0.75, colour = 'black', linetype = 'solid')+
  geom_abline(intercept = -1.010811, 
              slope = 0.268766, 
              size = 0.75, colour = 'red', linetype = 'dashed')+
  geom_line(data = prediction_data, 
          aes(x = total_count, y = predicted_ln_pc_ln_c), 
          inherit.aes = FALSE,  # <<--- forces it to ignore the global aes
          color = "red", size = 0.75, linetype = 'dotdash')+
   #geom_text(aes(label = species_name), 
            #size = 0.50, 
            #vjust = -0.5, hjust = 0.5, 
            #color = "black")+
  xlab("ln[Total Count]")+ 
  ylab("ln[Precaudal]-ln[Caudal]")+
  theme_classic()+
  theme(legend.position='none',
        axis.text = element_text(size=11),
        axis.title = element_text(size=11, face='bold'))+
  expand_limits(y = c(min(all_species_data$ln_pc_ln_c), max(all_species_data$ln_pc_ln_c) + 0.20))
c2

ggsave("Figures/Somitogenesis and Regionalisation/total_count_pc_c_ratio_pgls.jpeg", c2, dpi = 600, width = 10, height = 10, units = "cm")
```

```{r Find Mode for Count Data}

#convert count columns to numeric for some reason???
as_numeric <- c("total_count", "precaudal_count", "caudal_count")
vertebral_counts[as_numeric] <- lapply(vertebral_counts[as_numeric], as.numeric)

#R doesn't have a built in mode function!? Define one here:
get_mode <- function(x) {
  ux <- unique(x)
  ux[which.max(tabulate(match(x, ux)))]
}

#group and find the modal count for total, precaudal and caudal vertebrae
modal_species_data <- 
  vertebral_counts %>%
  group_by(species_name, tribe, haplochromine, water_system, depth, piscivore) %>%
  summarise(
    total_count = get_mode(total_count),
    precaudal_count = get_mode(precaudal_count),
    caudal_count = get_mode(caudal_count),
    .groups = "drop"  #avoids grouped warning
  )
```

```{r Subset data to tree}

#get tree tips
tree_tips <- all_species$tree$tip.label

# Filter modal_species_data to include only species present in the tree
modal_species_data_filtered <- modal_species_data %>%
  filter(species_name %in% tree_tips) %>%
  arrange(match(species_name, all_species$tree$tip.label))

#convert to matrix so pic function defined above can be re-used
rownames(modal_species_data_filtered) <- modal_species_data_filtered$species_name
modal_species_data_filtered <- as.matrix(modal_species_data_filtered)

#define the count columns to keep
count_columns <- c("total_count", "precaudal_count", "caudal_count")

#filter to retain only count columns
modal_species_data_filtered <- modal_species_data_filtered[, count_columns, drop = FALSE]

#set branch lengths to 1.00 (so that we can consider the relative magnitude of morphological differences across the phylogeny rather than scaling them by evolutionary time)
all_species$tree$edge.length <- all_species$tree$edge.length / all_species$tree$edge.length
```

```{r Define Function to Compute PICs}
#define function to compute pics for all traits in the all species matrix data
compute_pic <- function(matrix_data, tree) {
  #initialize a list to store PIC results
  pic_results <- list()
  
  #get column names
  col_names <- colnames(matrix_data)
  
  #compute PIC for each trait
  for (i in seq_along(col_names)) {
    pic_results[[col_names[i]]] <- ape::pic(matrix_data[, i], tree)
  }
  
  #convert results to a dataframe
  pic_df <- as.data.frame(do.call(cbind, pic_results))
  
  #add node labels
  pic_df$node <- rownames(pic_df)
  
  return(pic_df)
}
```

```{r Calculate PICs for Modal Counts (Branch Lengths 1)}
#calculate the PICs for each trait
all_species_pics_modal <- compute_pic(modal_species_data_filtered, all_species$tree)

#merge nodemap data with pics dataframe
all_species_pics_modal <- merge(all_species_pics_modal, node_map, by='node')

#define tribe as a factor 
all_species_pics_modal$tribe <- as.factor(all_species_pics_modal$tribe)
```

```{r Scatterplot of Precaudal and Caudal PICs (Homeotic Shifts)}
#plot pics of precaudal and caudal count (homeotic changes)
a1 <- ggplot(all_species_pics_modal, aes(x = caudal_count, y = precaudal_count))+
  geom_point(aes(
    color = case_when(
      abs(precaudal_count + caudal_count) < 1e-7 ~ "on_neg_line",  # y = -x
      abs(precaudal_count - caudal_count) < 1e-7 ~ "on_pos_line",  # y = x
      TRUE ~ "off_line"
    ),
    shape = case_when(
      abs(precaudal_count + caudal_count) < 1e-7 ~ "on_neg_line",
      abs(precaudal_count - caudal_count) < 1e-7 ~ "on_pos_line",
      TRUE ~ "off_line"
    ),
    size = case_when(
      abs(precaudal_count + caudal_count) < 1e-7 ~ "on_neg_line",
      abs(precaudal_count - caudal_count) < 1e-7 ~ "on_pos_line",
      TRUE ~ "off_line"
    ),
    alpha = case_when(
      abs(precaudal_count + caudal_count) < 1e-7 ~ 0.75,  
      abs(precaudal_count - caudal_count) < 1e-7 ~ 0.75,  
      TRUE ~ 0.75
    )
  ))+
  xlab("PICs of Caudal Count (Raw)") +
  ylab("PICs of Precaudal Count (Raw)") +
  scale_x_continuous(limits = c(-4, 4), breaks = seq(-4, 4, by = 2))+
  scale_y_continuous(limits = c(-4, 4), breaks = seq(-4, 4, by = 2))+
  geom_vline(xintercept = 0, linetype = 'dotdash', size = 0.50)+
  geom_hline(yintercept = 0, linetype = 'dotdash', size = 0.50)+
  geom_abline(slope = -1, size = 0.50, linetype = 'dotted', colour = 'blue', alpha = 1.00)+  # y = -x
  geom_abline(slope = 1, size = 0.50, linetype = 'dotted', colour = 'red', alpha = 1.00)+   # y = x
  scale_color_manual(values = c("on_neg_line" = "blue", "on_pos_line" = "red", "off_line" = "black"))+
  scale_shape_manual(values = c("on_neg_line" = 17, "on_pos_line" = 18, "off_line" = 19))+
  scale_size_manual(values = c("on_neg_line" = 2.5, "on_pos_line" = 2.5, "off_line" = 1))+
    #geom_text(
    #label = all_species_pics_modal$node,
    #nudge_x = 0, nudge_y = -0.10, check_overlap = TRUE
  #)+  # plot node labels to points
  theme_classic() +
  theme(
    legend.position = 'none',
    axis.text = element_text(size = 11),
    axis.title = element_text(size = 11, face = 'bold')
  )
a1

ggsave("Figures/Somitogenesis and Regionalisation/precaudal_caudal_pics_homeoetic_effects.jpeg", a1, dpi = 600, width = 10, height = 10, units = "cm")
```

```{r Distributions of Perpendicular Distances (Co-occuring Somitic Changes)}
#calculate the perpendicular distance of each point from the diagonal line (assuming slope is -1)
distances_homeotic_effects <- abs(all_species_pics_modal$caudal_count + all_species_pics_modal$precaudal_count) / sqrt(2)

#bind to dataframe
all_species_pics_modal$perpendicular_distance <- distances_homeotic_effects

#filter for pure homeotic shifts
pure_homeotic <- 
  all_species_pics_modal %>%
  filter(perpendicular_distance == 0)

#plot distribution of perpendicular distances (i.e. magnitude of co-occuring somitic effect (i.e. change in number of vertebrae/somites))
ggplot(all_species_pics_modal, aes(x=perpendicular_distance))+
  geom_histogram(bins = 20, colour='black')+
  xlab("Perpendicular Distance (Magnitude of Co-occuring Somitic Effect)")+
  theme_classic()

#histogram plotting the perpendicular distances -- log10(distance + 1) to transform but keep 0 as 0 
ggplot(all_species_pics_modal, aes(x=log10(perpendicular_distance + 1)))+
  geom_histogram(bins = 20, colour='black')+
  xlab("Log10[Perpendicular Distance + 1] (Magnitude of Co-occuring Somitic Effect)")+
  theme_classic()
```

```{r Homeotic Shifts versus Combined Effects}

#assign 'Zero', 'Non-Zero' and 'One' based on perpendicular_distance (Zero is y=-x, 1.00 is y=x, non-zero is neither)
all_species_pics_modal <- all_species_pics_modal %>%
  mutate(group = case_when(
    perpendicular_distance == 0 ~ "Zero",
    abs(perpendicular_distance - 1.00) < .Machine$double.eps^0.5 ~ "One",
    TRUE ~ "Non-Zero"
  ))

# cnvert 'group' column to a factor with specified levels and labels
all_species_pics_modal_2 <- all_species_pics_modal
all_species_pics_modal_2$group <- factor(all_species_pics_modal$group, 
                                         levels = c('One', 'Zero', 'Non-Zero'),
                                         labels = c('Balanced Changes', 'Pure Homeotic', 'Combined Effects'))  #define new labels

# Create bar plot
d1 <- ggplot(all_species_pics_modal_2, aes(x = group, fill = group)) +
  geom_bar(color = "black")+
  xlab("Predicted Developmental Change")+
  ylab("Count")+
  scale_fill_manual(values = c("Balanced Changes" = 'red', "Pure Homeotic" = 'blue', "Combined Effects" = 'grey'))+
   theme_classic() +
  theme(
    legend.position = 'none',
    axis.text = element_text(size = 11),
    axis.title = element_text(size = 11, face = 'bold'),
    axis.text.x = element_text(size=9)
  )
d1

ggsave("Figures/Somitogenesis and Regionalisation/bar_chart_number_changes.jpeg", d1, dpi = 600, width = 10, height = 10, units = "cm")
```

```{r Plot Tree with Node Circles Scaled to Perpendicular Distance (ggTree)}
#merge the continuous variable data into the tree node data
all_species_pics_modal$node <- as.integer(all_species_pics_modal$node)
tree_data <- all_species$tree %>% 
  fortify() %>%  #convert tree into a dataframe
  left_join(all_species_pics_modal, by = "node")

#randomly select two colours along plasma scale
#set.seed(1931)
#scale_cols <- c(plasma(10))

#ladderise tree
all_species_tree <- ladderize(all_species$tree)

#convert trait data to a dataframe with species as rownames
trait_df <- data.frame(another_trait = all_species_data$total_count)
rownames(trait_df) <- all_species_data$species_name  # Ensure row names match tree tip labels

#plotting the tree and scale node point size and colour to perpendicular distance
a1 <- ggtree(all_species_tree, branch.length = 'none', layout = 'circular', size=0.33) +
  geom_tiplab(size = 1.00, fontface = "bold", offset=1.75) +
  geom_nodepoint(aes(size = ifelse(tree_data$group == "One", tree_data$perpendicular_distance / 12, #scale balanced effects to 1.00, so that size reflects just 'combined effects/co-occuring somitic changes'
                                   tree_data$perpendicular_distance), 
                     colour = tree_data$perpendicular_distance, 
                     shape = tree_data$group)) +
  scale_shape_manual(values = c("Zero" = 17, "One" = 18, "Non-Zero" = 20)) + # Shapes
  scale_size_continuous(range = c(1.00, 8.00)) + # Adjust overall scaling of node sizes
  scale_color_gradient(low='grey', high='black') + # Adjust colour scale
  theme(legend.position = "none")
a1

#add heatmap for total vertebral count
#randomly select two colours along plasma scale
set.seed(1931)
scale_cols <- c(plasma(100))

a1 <- gheatmap(a1, trait_df, offset = -1, width = 0.05, 
               colnames = FALSE)+
  scale_fill_gradient(low = scale_cols[1], high = scale_cols[100])  # Adjust color scale

a1 + theme(legend.position = "none")

ggsave("Figures/Somitogenesis and Regionalisation/tree_co_occuring_somitic_homeoetic_effects.svg", dpi = 600, width = 21, height = 29.7, units = "cm")
```

#Phylogenetic Independent Contrasts (Original, Time Calibrated Scale)

```{r Calculate PICs for Modal Counts (Branch Lengths Time)}

#reimport species_data
all_species <- readRDS('data/all_species.rds')

#prune tree to remove lV species which predict very large co-occurring somitic change (because of very little genetic variation/short branch lengths)
species_to_remove <- c("Haplochromis labiatus", "Haplochromis mentatus")

#filter from the modal data
modal_species_data_filtered <- modal_species_data_filtered[!rownames(modal_species_data_filtered) %in% species_to_remove, , drop = FALSE]

#prune them from the tree
all_species_tree <- drop.tip(all_species$tree, species_to_remove)

#calculate the PICs for each trait
all_species_pics_modal_time <- compute_pic(modal_species_data_filtered, all_species_tree)

#add haplochromine label
all_species_pics_modal_time <- all_species_pics_modal_time %>%
  mutate(node = as.numeric(node)) %>%  # Convert node to numeric
  mutate(haplochromine = ifelse(between(node, 439, 641), "Haplochromine", "Non-Haplochromine"))

#add lake malawi and lake victoria label:
all_species_pics_modal_time <- all_species_pics_modal_time %>%
  mutate(water_system = case_when(
    between(node, 447, 563) ~ "Lake Malawi",     # Assign "Lake Malawi" for nodes between 447 and 563
    between(node, 566, 598) ~ "Lake Victoria",   # Assign "Lake Victoria" for nodes between 566 and 598
    TRUE ~ "Non-Haplochromine"                    # Assign "Non-Haplochromine" for all other nodes
  ))
```

```{r Scatterplot of Precaudal and Caudal PICs (Homeotic Shifts - PICs, Time)}
#define haplochromine as a factor
all_species_pics_modal_time$haplochromine <- as.factor(all_species_pics_modal_time$haplochromine)

#plot pics of precaudal and caudal count (homeotic changes)
b1 <- ggplot(all_species_pics_modal_time, aes(x = caudal_count, y = precaudal_count))+
  geom_point(aes(
    color = case_when(
      abs(precaudal_count + caudal_count) < 1e-7 ~ "on_neg_line",  # y = -x
      abs(precaudal_count - caudal_count) < 1e-7 ~ "on_pos_line",  # y = x
      TRUE ~ "off_line"
    ),
    shape = case_when(
      abs(precaudal_count + caudal_count) < 1e-7 ~ "on_neg_line",
      abs(precaudal_count - caudal_count) < 1e-7 ~ "on_pos_line",
      TRUE ~ "off_line"
    ),
    size = case_when(
      abs(precaudal_count + caudal_count) < 1e-7 ~ "on_neg_line",
      abs(precaudal_count - caudal_count) < 1e-7 ~ "on_pos_line",
      TRUE ~ "off_line"
    ),
    alpha = haplochromine  # Map alpha to haplochromine
  )) +
  xlab("PICs of Caudal Count") +
  ylab("PICs of Precaudal Count") +
  scale_x_continuous(limits = c(-4, 4), breaks = seq(-4, 4, by = 2))+
  scale_y_continuous(limits = c(-4, 4), breaks = seq(-4, 4, by = 2))+
  geom_vline(xintercept = 0, linetype = 'dotdash', size = 0.50)+
  geom_hline(yintercept = 0, linetype = 'dotdash', size = 0.50)+
  geom_abline(slope = -1, size = 0.50, linetype = 'dotted', colour = 'blue', alpha = 1.00)+  #y = -x
  geom_abline(slope = 1, size = 0.50, linetype = 'dotted', colour = 'red', alpha = 1.00)+   #y = x
  scale_color_manual(values = c("on_neg_line" = "blue", "on_pos_line" = "red", "off_line" = "black"))+
  scale_shape_manual(values = c("on_neg_line" = 17, "on_pos_line" = 18, "off_line" = 19))+
  scale_size_manual(values = c("on_neg_line" = 2.5, "on_pos_line" = 2.5, "off_line" = 1.5))+
  scale_alpha_manual(values = c("Non-Haplochromine" = 0.15, "Haplochromine" = 0.80))+
 #geom_text(#label = all_species_pics_modal$node,
 #nudge_x = 0, nudge_y = -0.10, check_overlap = TRUE)+  # plot node labels to points
  theme_classic()+
  theme(
    legend.position = 'none',
    axis.text = element_text(size = 11),
    axis.title = element_text(size = 11, face = 'bold'))

b1

ggsave("Figures/Somitogenesis and Regionalisation/precaudal_caudal_pics_homeoetic_effects_time_scale.jpeg", b1, dpi = 600, width = 10, height = 10, units = "cm")
```

```{r Distributions of Perpendicular Distances (Co-occuring Somitic Changes)}
#calculate the perpendicular distance of each point from the diagonal line (assuming slope is -1)
distances_homeotic_effects <- abs(all_species_pics_modal_time$caudal_count + all_species_pics_modal_time$precaudal_count) / sqrt(2)

#bind to dataframe
all_species_pics_modal_time$perpendicular_distance <- distances_homeotic_effects

#filter for pure homeotic shifts
pure_homeotic <- 
  all_species_pics_modal_time %>%
  filter(perpendicular_distance == 0)

#plot distribution of perpendicular distances (i.e. magnitude of co-occuring somitic effect (i.e. change in number of vertebrae/somites))
ggplot(all_species_pics_modal_time, aes(x=perpendicular_distance))+
  geom_histogram(bins = 20, colour='black')+
  xlab("Perpendicular Distance (Magnitude of Co-occuring Somitic Effect)")+
  theme_classic()

#histogram plotting the perpendicular distances -- log10(distance + 1) to transform but keep 0 as 0 
ggplot(all_species_pics_modal_time, aes(x=log10(perpendicular_distance + 1)))+
  geom_histogram(bins = 20, colour='black')+
  xlab("Log10[Perpendicular Distance + 1] (Magnitude of Co-occuring Somitic Effect)")+
  theme_classic()
```

```{r Homeotic Shifts versus Combined Effects}

#assign 'Zero', 'Non-Zero' and 'One' based on perpendicular_distance (Zero is y=-x, 1.00 is y=x, non-zero is neither)
all_species_pics_modal_time <- all_species_pics_modal_time %>%
  mutate(group = case_when(
    perpendicular_distance == 0 ~ "Zero",
    perpendicular_distance >= 0.95 & perpendicular_distance <= 1.05 ~ "One",
    TRUE ~ "Non-Zero"
  ))

# cnvert 'group' column to a factor with specified levels and labels
all_species_pics_modal_time_2 <- all_species_pics_modal_time
all_species_pics_modal_time_2$group <- factor(all_species_pics_modal_time$group, 
                                         levels = c('One', 'Zero', 'Non-Zero'),
                                         labels = c('Balanced Changes', 'Pure Homeotic', 'Combined Effects'))  #define new labels

# Create bar plot
d1 <- ggplot(all_species_pics_modal_time_2, aes(x = group, fill = group)) +
  geom_bar(color = "black")+
  xlab("Predicted Developmental Change")+
  ylab("Count")+
  scale_fill_manual(values = c("Balanced Changes" = 'red', "Pure Homeotic" = 'blue', "Combined Effects" = 'grey'))+
   theme_classic() +
  theme(
    legend.position = 'none',
    axis.text = element_text(size = 11),
    axis.title = element_text(size = 11, face = 'bold'),
    axis.text.x = element_text(size=9)
  )
d1

ggsave("Figures/Somitogenesis and Regionalisation/bar_chart_number_changes_time.jpeg", d1, dpi = 600, width = 10, height = 10, units = "cm")
```

```{r Plot Tree with Node Circles Scaled to Perpendicular Distance (ggTree)}
#merge the continuous variable data into the tree node data
all_species_pics_modal_time$node <- as.integer(all_species_pics_modal_time$node)

tree_data <- all_species_tree %>% 
  fortify() %>%  #convert tree into a dataframe
  left_join(all_species_pics_modal_time, by = "node")
#randomly select two colours along plasma scale
#set.seed(1931)
#scale_cols <- c(viridis(10))

#plotting the tree and scale node point size and colour to perpendicular distance
a1 <- ggtree(all_species_tree, branch.length = 'none', layout = 'circular', size=0.33) +
  geom_tiplab(size = 1.00, fontface = "bold") +
  geom_nodepoint(aes(size = ifelse(tree_data$group == "One", tree_data$perpendicular_distance / 12, #scale balanced effects to 1.00, so that size reflects just 'combined effects/co-occuring somitic changes'
                                   tree_data$perpendicular_distance), 
                     colour = tree_data$perpendicular_distance, 
                     shape = tree_data$group)) +
  scale_shape_manual(values = c("Zero" = 17, "One" = 18, "Non-Zero" = 20)) + # Shapes
  scale_size_continuous(range = c(1.00, 8.00)) + # Adjust overall scaling of node sizes
  scale_color_gradient(low='grey', high='black') + # Adjust colour scale
  theme(legend.position = "none")
a1

ggsave("Figures/Somitogenesis and Regionalisation/tree_co_occuring_somitic_homeoetic_effects_with_time.pdf", dpi = 600, width = 21, height = 29.7, units = "cm")
```

```{r Plot Tree with Node Circles Scaled to Perpendicular Distance (ggTree)}
#merge the continuous variable data into the tree node data
all_species_pics_modal_time$node <- as.integer(all_species_pics_modal_time$node)
tree_data <- all_species_tree %>% 
  fortify() %>%  #convert tree into a dataframe
  left_join(all_species_pics_modal_time, by = "node")

#randomly select two colours along plasma scale
#set.seed(1931)
#scale_cols <- c(plasma(10))

#ladderise tree
all_species_tree <- ladderize(all_species_tree)


#convert trait data to a dataframe with species as rownames
trait_df <- data.frame(another_trait = all_species_data$total_count)
rownames(trait_df) <- all_species_data$species_name  # Ensure row names match tree tip labels

#plotting the tree and scale node point size and colour to perpendicular distance
a1 <- ggtree(all_species_tree, branch.length = 'none', layout = 'circular', size=0.33) +
  geom_tiplab(size = 1.00, offset=1.75, colour='grey') +
  geom_nodepoint(aes(size = ifelse(tree_data$group == "One", tree_data$perpendicular_distance / 12, #scale balanced effects to 1.00, so that size reflects just 'combined effects/co-occuring somitic changes'
                                   tree_data$perpendicular_distance), 
                     colour = tree_data$perpendicular_distance, 
                     shape = tree_data$group)) +
  scale_shape_manual(values = c("Zero" = 17, "One" = 18, "Non-Zero" = 20)) + # Shapes
  scale_size_continuous(range = c(1.00, 8.00)) + # Adjust overall scaling of node sizes
  scale_color_gradient(low='grey', high='black') + # Adjust colour scale
  theme(legend.position = "none")
a1

#add heatmap for total vertebral count
#randomly select two colours along plasma scale
set.seed(1931)
scale_cols <- c(plasma(100))

a1 <- gheatmap(a1, trait_df, offset = -1, width = 0.05, 
               colnames = FALSE)+
  scale_fill_gradient(low = scale_cols[1], high = scale_cols[100])  # Adjust color scale

a1 + theme(legend.position = "none")

ggsave("Figures/Somitogenesis and Regionalisation/tree_co_occuring_somitic_homeoetic_effects_cladogram.svg", dpi = 600, width = 21, height = 29.7, units = "cm")
```

```{r Plot Tree with Node Circles Scaled to Perpendicular Distance (ggTree - with branch lengths)}
#merge the continuous variable data into the tree node data
all_species_pics_modal_time$node <- as.integer(all_species_pics_modal_time$node)
tree_data <- all_species_tree %>% 
  fortify() %>%  #convert tree into a dataframe
  left_join(all_species_pics_modal_time, by = "node")

#randomly select two colours along plasma scale
#set.seed(1931)
#scale_cols <- c(plasma(10))

#ladderise tree
all_species_tree <- ladderize(all_species_tree)


#convert trait data to a dataframe with species as rownames
trait_df <- data.frame(another_trait = all_species_data$total_count)
rownames(trait_df) <- all_species_data$species_name  # Ensure row names match tree tip labels

#plotting the tree and scale node point size and colour to perpendicular distance
a1 <- ggtree(all_species_tree, branch.length = TRUE, layout = 'circular', size=0.33) +
  geom_tiplab(size = 1.00, offset=1.75, colour='grey') +
  geom_nodepoint(aes(size = ifelse(tree_data$group == "One", tree_data$perpendicular_distance / 12, #scale balanced effects to 1.00, so that size reflects just 'combined effects/co-occuring somitic changes'
                                   tree_data$perpendicular_distance), 
                     colour = tree_data$perpendicular_distance, 
                     shape = tree_data$group)) +
  scale_shape_manual(values = c("Zero" = 17, "One" = 18, "Non-Zero" = 20)) + # Shapes
  scale_size_continuous(range = c(1.00, 8.00)) + # Adjust overall scaling of node sizes
  scale_color_gradient(low='grey', high='black') + # Adjust colour scale
  theme(legend.position = "none")
a1

#add heatmap for total vertebral count
#randomly select two colours along plasma scale
set.seed(1931)
scale_cols <- c(plasma(100))

a1 <- gheatmap(a1, trait_df, offset = 1, width = 0.05, 
               colnames = FALSE)+
  scale_fill_gradient(low = scale_cols[1], high = scale_cols[100])  # Adjust color scale

a1 + theme(legend.position = "none")

ggsave("Figures/Somitogenesis and Regionalisation/tree_co_occuring_somitic_homeoetic_effects_time_branches_scaled.svg", dpi = 600, width = 21, height = 29.7, units = "cm")
```

```{r Combined Bar Charts of Predicted Developmental Change}
#classify groups for all_species_pics_modal (Raw)
all_species_pics_modal <- all_species_pics_modal %>%
  mutate(group = case_when(
    perpendicular_distance == 0 ~ "Zero",
    near(perpendicular_distance, 1) ~ "One",  #handle potential floating-point precision issues
    TRUE ~ "Non-Zero"
  ))

#classify groups for all_species_pics_modal_time (Evolutionary Rates)
all_species_pics_modal_time <- all_species_pics_modal_time %>%
  mutate(group = case_when(
    perpendicular_distance == 0 ~ "Zero",
    perpendicular_distance >= 0.95 & perpendicular_distance <= 1.05 ~ "One",
    TRUE ~ "Non-Zero"
  ))

#ensure factor levels are set before merging
group_levels <- c("One", "Zero", "Non-Zero")
group_labels <- c("Balanced Changes", "Pure Homeotic", "Combined Effects")

all_species_pics_modal$group <- factor(all_species_pics_modal$group, levels = group_levels, labels = group_labels)
all_species_pics_modal_time$group <- factor(all_species_pics_modal_time$group, levels = group_levels, labels = group_labels)

#add identifier column for merging
all_species_pics_modal$source <- "Raw"
all_species_pics_modal_time$source <- "Evolutionary Rates"

#combine both datasets
all_species_pics_combined <- bind_rows(all_species_pics_modal, all_species_pics_modal_time)

#ensure ggplot recognizes all levels even if some are missing in one dataset
all_species_pics_combined$group <- factor(all_species_pics_combined$group, levels = group_labels)

#plot combined predicted developmental changes (i.e. for the raw and time PICs)
x1 <- ggplot(all_species_pics_combined, aes(x = group, fill = source))+
  geom_bar(position = "dodge", colour = 'black')+
  labs(x = "Category", y = "Count", fill = "Dataset")+ 
  scale_fill_manual(values = c("Raw" = "#0072B2", "Evolutionary Rates" = "#E69F00"))+
  xlab("Developmental Change")+
  ylab("Count (Nodes)")+
  coord_flip()+
  theme_classic()+
  theme(
    legend.position = 'none',
    axis.text = element_text(size = 10),
    axis.title = element_text(size = 10, face = 'bold'),
    axis.text.y = element_blank(),  # Removes Y axis labels
    axis.ticks.y = element_blank()  # Removes Y tick marks
  ) +
  scale_x_discrete(limits = rev(levels(all_species_pics_combined$group)))
x1

ggsave("Figures/Somitogenesis and Regionalisation/combined_bar_chart_predicted_dev_change.jpeg", dpi = 600, width = 20, height = 5, units = "cm")
```

# Intraspecific Variation

```{r Data Wrangling}
#group according to species (and other variables) and calculate the standard deviation of the RAW precaudal, caudal and total counts
vertebral_counts_intraspecific_variation <- vertebral_counts %>% 
  group_by(
    species_name, genus, tribe, haplochromine, water_system,
    lm_ecology, mouthbrooder, depth, piscivore
  ) %>% 
  summarise(
    precaudal_count_sd = sd(precaudal_count, na.rm = TRUE),
    precaudal_count = mean(precaudal_count, na.rm = TRUE),
    caudal_count_sd = sd(caudal_count, na.rm = TRUE),
    caudal_count = mean(caudal_count, na.rm = TRUE),
    total_count_sd = sd(total_count, na.rm = TRUE),
    total_count = mean(total_count, na.rm = TRUE),
    total_count_cv = total_count_sd / total_count, #coefficient of variation, may not be necessary mean range of vertebrae is narrow between species (25-40)
    n_count = n()
  )

vertebral_counts_intraspecific_variation_filtered <- 
  vertebral_counts_intraspecific_variation %>%
  filter(n_count >= 5.00)

#get ranges
help <- vertebral_counts %>% 
  group_by(
    species_name, genus, tribe, haplochromine, water_system,
    lm_ecology, mouthbrooder, depth, piscivore
  ) %>% 
  summarise(
    total_range = max(total_count) - min(total_count)
  )
```

```{r Prune McGee 2020 Tree to Species with Greater Than or Equal to 5 Specimens}

#import the tree
mcgee_2020 <- read.tree('Phylogenies/phylogenies_mcgee_2020/phylogeny_mcgee_2020.tre')

#format tip labels to match the formatting of the species_names in data-frames
mcgee_2020$tip.label <- gsub("_", " ", mcgee_2020$tip.label)

#ladderise the tree
mcgee_2020 <- ladderize(mcgee_2020)

#prune the mcgee 2020 tree to data
mcgee_intraspecific_pruned_tree <- drop.tip(mcgee_2020, 
                                            which(!mcgee_2020$tip.label %in% vertebral_counts_intraspecific_variation_filtered$species_name))

#subset data frame to include only species present in the pruned tree
vertebral_counts_intraspecific_pruned <- vertebral_counts_intraspecific_variation_filtered[
 vertebral_counts_intraspecific_variation_filtered$species_name %in% mcgee_intraspecific_pruned_tree$tip.label, 
]

#reorder data frame based on the order of tip labels in the pruned tree
vertebral_counts_intraspecific_pruned <- vertebral_counts_intraspecific_pruned[
  match(mcgee_intraspecific_pruned_tree$tip.label, vertebral_counts_intraspecific_pruned$species_name), 
]
```

```{r Intraspecific Variation in Precaudal and Caudal Counts}

#group according to species (and other variables) and calculate variance of precaudal and caudal counts
precaudal_caudal_variance <- vertebral_counts %>% 
  group_by(
    species_name, genus, tribe, haplochromine, water_system,
    lm_ecology, mouthbrooder, depth, piscivore
  ) %>% 
  filter(n() >= 5) %>%  #filter groups with at least 5 observations
  summarize(
    precaudal_variance = var(precaudal_count, na.rm = TRUE),
    caudal_variance = var(caudal_count, na.rm = TRUE),
    total_variance = var(total_count, na.rm = TRUE),
    cv_precaudal = sd(precaudal_count, na.rm = TRUE) / mean(precaudal_count, na.rm = TRUE),
    cv_caudal = sd(caudal_count, na.rm = TRUE) / mean(caudal_count, na.rm = TRUE),
    n_count = n(),
    .groups = "drop"
  )

#subset data frame to include only species present in the pruned tree
precaudal_caudal_variance_pruned <-precaudal_caudal_variance[
 precaudal_caudal_variance$species_name %in% mcgee_intraspecific_pruned_tree$tip.label, 
]

#reorder data frame based on the order of tip labels in the pruned tree
precaudal_caudal_variance_pruned  <- precaudal_caudal_variance_pruned[
  match(mcgee_intraspecific_pruned_tree$tip.label, precaudal_caudal_variance_pruned$species_name), 
]
```

```{r Intraspecific Variation in Precaudal Counts and Caudal Counts (Variance)}
#test for sig difference in medians between precaudal and caudal cv
wilcox_precaual_caudal_var <- wilcox.test(precaudal_caudal_variance_pruned$precaudal_variance, precaudal_caudal_variance_pruned$caudal_variance, paired = TRUE, alternative = 'less')
wilcox_precaual_caudal_var

#calculate ratio of caudal:precaudal variance
mean(precaudal_caudal_variance_pruned$caudal_variance) / mean(precaudal_caudal_variance_pruned$precaudal_variance)

#convert to long format for plotting
species_pc_c_variation_long <- precaudal_caudal_variance_pruned %>%
  pivot_longer(
    cols = c(precaudal_variance, caudal_variance),
    names_to = "vertebral_region",
    values_to = "variance"
  )

#calculate the proportion of variance from precaudal and caudal to total variance (since var(TC) = var(PC) + var(C) + 2Co(PC,C), but PC~C is around 0, Therefore, reduces to var(TC) = var(PC) + var(C) + 0)
variance_data <- precaudal_caudal_variance_pruned %>%
  mutate(
    proportion_PC = precaudal_variance / (precaudal_variance + caudal_variance),
    proportion_C = caudal_variance / (precaudal_variance + caudal_variance)
  )

#Check how much precaudal variance contributes
summary(variance_data$proportion_PC)  
summary(variance_data$proportion_C) 

#recode the levels of the domain
species_pc_c_variation_long$vertebral_region <- dplyr::recode(species_pc_c_variation_long$vertebral_region,
                                                  "precaudal_variance" = "PC",
                                                  "caudal_variance" = "CC")

#reordering the levels of water_system
species_pc_c_variation_long$vertebral_region <- factor(species_pc_c_variation_long$vertebral_region,
                                                  levels = c("PC", "CC"))

#colours taken from Wong colour blind palette
water_system_colours <- c("Lake Victoria" = "#43AA8B", 
                          "Lake Malawi" = "#D55E00", 
                          "Lake Tanganyika" = "#0072B2", 
                          "Riverine" = "#CC79A7")

#plot the precaudal and caudal variances
set.seed(1931) #for jitter
o1 <- ggplot(species_pc_c_variation_long, aes(x=vertebral_region, y=variance))+
  geom_boxplot(outlier.shape = NA, linewidth=0.5)+
  geom_jitter(aes(shape=haplochromine, colour=water_system), 
              size=1, width=0.35, alpha=0.50)+
  scale_colour_manual(values = water_system_colours)+
  scale_shape_manual(values = c('Haplochromine' = 19, 'Other' = 17))+
  ylab(expression(bold(paste("Count Variance (", sigma^2, ")"))))+
  xlab("Vertebral Domain")+
  theme_classic()+
          theme(legend.position='none',
                 axis.text = element_text(size=11),
                 axis.title = element_text(size=11, face='bold'))
o1

ggsave("Figures/Somitogenesis and Regionalisation/precaudal_var_caudal_var_boxplot.jpeg", o1, dpi = 600, width = 5, height = 10, units = "cm")
```

```{r Intraspecific Variation in Precaudal Counts and Caudal Counts (Coefficient of Variation)}

#test for sig difference in medians between precaudal and caudal cv
wilcox_precaual_caudal_cv <- wilcox.test(precaudal_caudal_variance_pruned$cv_precaudal, precaudal_caudal_variance_pruned$cv_caudal, paired = TRUE)
wilcox_precaual_caudal_cv

#convert to long format for plotting
species_pc_c_variation_long <- precaudal_caudal_variance_pruned %>%
  pivot_longer(
    cols = c(cv_precaudal, cv_caudal),
    names_to = "vertebral_region",
    values_to = "coeff_var"
  )

#recode the levels of the domain
species_pc_c_variation_long$vertebral_region <- dplyr::recode(species_pc_c_variation_long$vertebral_region,
                                                  "cv_precaudal" = "PC",
                                                  "cv_caudal" = "CC")


#reordering the levels of water_system
species_pc_c_variation_long$vertebral_region <- factor(species_pc_c_variation_long$vertebral_region,
                                                  levels = c("PC", "CC"))

#colours taken from Wong colour blind palette
water_system_colours <- c("Lake Victoria" = "#43AA8B", 
                          "Lake Malawi" = "#D55E00", 
                          "Lake Tanganyika" = "#0072B2", 
                          "Riverine" = "#CC79A7")

#plot the precaudal and caudal cvs
set.seed(1931) #for jitter
o2 <- ggplot(species_pc_c_variation_long, aes(x=vertebral_region, y=coeff_var))+
  geom_boxplot(outlier.shape = NA, linewidth=0.5)+
   geom_jitter(aes(shape=haplochromine, colour=water_system), 
              size=1, width=0.35, alpha=0.50)+
  scale_colour_manual(values = water_system_colours)+
  scale_shape_manual(values = c('Haplochromine' = 19, 'Other' = 17))+
  ylab(expression(bold(paste("Count CV (", sigma, " / ", mu, ")"))))+
  xlab("Vertebral Domain")+
  theme_classic()+
          theme(legend.position='none',
                 axis.text = element_text(size=11),
                 axis.title = element_text(size=11, face='bold'))
o2

ggsave("Figures/Somitogenesis and Regionalisation/precaudal_cv_caudal_cv_boxplot.jpeg", o2, dpi = 600, width = 5, height = 10, units = "cm")
```

```{r Phylogenetic Signal in Precaudal and Caudal CV}

#set seed for simulations 
set.seed(1931)

#precaudal

#convert CV values to a named vector
precaudal_cv_values <- precaudal_caudal_variance_pruned$cv_precaudal
names(precaudal_cv_values) <- precaudal_caudal_variance_pruned$species_name

#test for phylogenetic signal
#Blomberg's (2003) K:
K_precaudal_cv <- phylosig(mcgee_intraspecific_pruned_tree, precaudal_cv_values, nsim=100000, method = "K", test = TRUE)
K_precaudal_cv

#Pagel's (1999) Lambda:
lambda_precaudal_cv <- phylosig(mcgee_intraspecific_pruned_tree, precaudal_cv_values, nsim=100000, method = "lambda", test = TRUE)
lambda_precaudal_cv

#plot distributions
plot(K_precaudal_cv)
plot(lambda_precaudal_cv)
#phylogenetic signal is indistinguishable from 0. for K but not lambda. 

#caudal

#convert CV values to a named vector
caudal_cv_values <- precaudal_caudal_variance_pruned$cv_caudal
names(caudal_cv_values) <- precaudal_caudal_variance_pruned$species_name

#test for phylogenetic signal
#Blomberg's (2003) K:
K_caudal_cv <- phylosig(mcgee_intraspecific_pruned_tree, caudal_cv_values, nsim=100000, method = "K", test = TRUE)
K_caudal_cv

#Pagel's (1999) Lambda:
lambda_caudal_cv <- phylosig(mcgee_intraspecific_pruned_tree, caudal_cv_values, nsim=100000, method = "lambda", test = TRUE)
lambda_caudal_cv

#plot distributions
plot(K_caudal_cv)
plot(lambda_caudal_cv)
#phylogenetic signal is indistinguishable from 0. for K but not lambda. 
```

```{r Plot Null Distributions/Likelihood Profiles (Precaudal Count CV)}

#Null Distribution for K
sim_k_precaudal_cv <- as.data.frame(K_precaudal_cv$sim.K)
observed_k_precaudal_cv <- K_precaudal_cv$K

#plot the Null distribution (K)
h1 <- ggplot(sim_k_precaudal_cv, aes(x=K_precaudal_cv$sim.K))+
  geom_histogram(aes(y = ..count..), bins = 20, fill = "lightgray", color = "black", alpha = 0.7)+
  geom_vline(xintercept = observed_k_precaudal_cv,  color = "red", size = 1, linetype='dashed')+
  ylim(0, 22000)+
  xlab("Simulated K")+
  ylab("Count")+
  theme_classic()+
  theme(legend.position='none',
                 axis.text = element_text(size=11),
                 axis.title = element_text(size=11, face='bold'))
h1

ggsave("Figures/Somitogenesis and Regionalisation/precaudal_cv_k_null.jpeg", h1, dpi = 600, width = 10, height = 10, units = "cm")


#Likelihood profile (Lambda)
jpeg("Figures/Somitogenesis and Regionalisation/precaudal_cv_lambda_likelihood_profile.jpg", 
     width = 12, height = 12, units = "cm", res = 600, quality = 95)

par(font.lab = 2)  # Make axis labels bold
plot(lambda_precaudal_cv)

dev.off()
```

```{r Plot Null Distributions/Likelihood Profiles (Caudal Count CV)}

#Null Distribution for K
sim_k_caudal_cv <- as.data.frame(K_caudal_cv$sim.K)
observed_k_caudal_cv <- K_caudal_cv$K

#plot the Null distribution (K)
h1 <- ggplot(sim_k_caudal_cv, aes(x=K_caudal_cv$sim.K))+
  geom_histogram(aes(y = ..count..), bins = 20, fill = "lightgray", color = "black", alpha = 0.7)+
  geom_vline(xintercept = observed_k_caudal_cv,  color = "red", size = 1, linetype='dashed')+
  ylim(0, 30000)+
  xlab("Simulated K")+
  ylab("Count")+
  theme_classic()+
  theme(legend.position='none',
                 axis.text = element_text(size=11),
                 axis.title = element_text(size=11, face='bold'))
h1

ggsave("Figures/Somitogenesis and Regionalisation/caudal_cv_k_null.jpeg", h1, dpi = 600, width = 10, height = 10, units = "cm")


#Likelihood profile (Lambda)
jpeg("Figures/Somitogenesis and Regionalisation/caudal_cv_lambda_likelihood_profile.jpg", 
     width = 12, height = 12, units = "cm", res = 600, quality = 95)

par(font.lab = 2)  # Make axis labels bold
plot(lambda_caudal_cv)

dev.off()
```

```{r Plot Distribution of Total Counts}
#plot distribution of total counts (checking for normality)
y2 <- ggplot(vertebral_counts_intraspecific_pruned, aes(x = total_count))+
  geom_histogram(aes(y = ..density..), 
                 binwidth = 1, fill = "lightgray", color = "black", alpha = 0.7)+
  stat_function(fun = dnorm, args = list(mean = mean(vertebral_counts_intraspecific_pruned$total_count),
                                         sd = sd(vertebral_counts_intraspecific_pruned$total_count)),
                color = "red", size = 1)+ #plot distribution curve over the histogram
  xlab("Total Vertebral Count")+
  ylab("Density")+
  theme_classic()+
  theme(legend.position='none',
                 axis.text = element_text(size=11),
                 axis.title = element_text(size=11, face='bold'))
#seems pretty normally distributed to me
y2

ggsave("Figures/Somitogenesis and Regionalisation/total_count_distribution_at_least_5.jpeg", y2, dpi = 600, width = 10, height = 10, units = "cm")

#shapiro test to double check normality:
shapiro.test(vertebral_counts_intraspecific_pruned$total_count)
#it's normal! W = 0.99534, p-value = 0.5728
```

```{r Compare Residuals for Total Count SD ~ N}
par(mfrow = c(2, 2))
plot(lm(total_count_sd ~ n_count, data = vertebral_counts_intraspecific_pruned))  #non-transformed
plot(lm(log(total_count_sd + 1) ~ log(n_count), data = vertebral_counts_intraspecific_pruned)) #transformed
par(mfrow = c(1, 1)) #reset layout

##note that residuals for log-transformed fit better on Q-Q plot (specifically species with high ncounts), significant relationship in raw was being driven by these species (of which there are relatively few). Unlikely that intraspecific variation artifically increasing with increased counts. 
```

```{r Scatterplot and LM (Total Count SD ~ n_count)}
#sense check to make sure that intraspecific variation doesn't correlate with n_count (i.e. if we count more indivdiuals does the variation increase -- which could be error)

#fit dirty lm to check (dirty because n_count is discrete, but log transform anyway)
total_count_sd_n_count_log <- lm(log(total_count_sd + 1) ~ log(n_count), 
                                 data = vertebral_counts_intraspecific_pruned)
summary(total_count_sd_n_count_log)

#spearman rank coefficient just to double check
cor.test(vertebral_counts_intraspecific_pruned$total_count_sd, 
         vertebral_counts_intraspecific_pruned$n_count, 
         method = "spearman")

#scatter plot to visually inspect the relationship
y1 <- ggplot(vertebral_counts_intraspecific_pruned, aes(x = log(n_count), y = log(total_count_sd + 1.00)))+
  geom_point(size = 1.50, alpha=0.50)+
  geom_smooth(method = "lm", se = TRUE, colour='black')+
  xlab("ln[Number of Individuals]") +
  ylab(expression(bold("ln[Total Count " ~ sigma ~ "+ 1]")))+
  theme_classic()+
  theme(legend.position='none',
                 axis.text = element_text(size=11),
                 axis.title = element_text(size=11, face='bold'))
y1
  
##weak positive correlation (and insignificant) between the total count SD and n_count -- means that addition of sampling errors not going to be a major problem

ggsave("Figures/Somitogenesis and Regionalisation/checking_n_count_variation.jpeg", dpi = 600, width = 10, height = 10, units = "cm")
```

```{r Plot Total Count CV versus Total Count}
# Create bins for the total_count variable
vertebral_counts_intraspecific_pruned <- vertebral_counts_intraspecific_pruned %>%
  mutate(total_count_bin = cut(total_count, breaks = seq(25, 40, by = 3), 
                               include.lowest = TRUE, right = FALSE))

#convert the factor levels to the desired format for prettier plotting
levels(vertebral_counts_intraspecific_pruned$total_count_bin) <- gsub(",","-", levels(vertebral_counts_intraspecific_pruned$total_count_bin))
levels(vertebral_counts_intraspecific_pruned$total_count_bin) <- gsub("\\[", "[", levels(vertebral_counts_intraspecific_pruned$total_count_bin))
levels(vertebral_counts_intraspecific_pruned$total_count_bin) <- gsub("\\)", "]", levels(vertebral_counts_intraspecific_pruned$total_count_bin))

#colours taken from Wong colour blind palette
water_system_colours <- c("Lake Victoria" = "#43AA8B", 
                          "Lake Malawi" = "#D55E00", 
                          "Lake Tanganyika" = "#0072B2", 
                          "Riverine" = "#CC79A7")

#plot boxplots of total_count_cv for each total_count_bin
c1 <- ggplot(vertebral_counts_intraspecific_pruned, aes(x = total_count_bin, y = total_count_cv))+
   geom_boxplot(outlier.shape = NA, linewidth=0.5)+
   geom_jitter(aes(shape=haplochromine, colour=water_system), 
              size=1.25, width=0.35, alpha=0.75)+
  scale_colour_manual(values = water_system_colours)+
  scale_shape_manual(values = c('Haplochromine' = 19, 'Other' = 17))+
  geom_hline(aes(yintercept = median(total_count_cv)), 
             color = "red", linetype = "dashed", size = 1)+ #add median line
  xlab('Total Vertebral Count')+
  ylab(expression(bold(paste("Total Vertebral Count CV (", sigma, " / ", mu, ")"))))+
  theme_classic()+
          theme(legend.position='none',
                 axis.text = element_text(size=11),
                 axis.title = element_text(size=11, face='bold'))
c1

ggsave("Figures/Somitogenesis and Regionalisation/total_count_cv_by_total_count_bin.jpeg", c1, dpi = 600, width = 10, height = 10, units = "cm")
```

```{r Define function Permutation Kruskal-Wallis Accounting for phylogeny}
#define function to perform Kruskal-Wallis test between groups, accounting for phylogeny
phylo_kruskal <- function(trait, group, phy, nperm = 10000) { #define number of permutations (10k)
  
  observed_kw <- kruskal.test(trait ~ group)$statistic  #original test statistic (i.e. no phylogenetic correction, as above)
  
  perm_kw <- numeric(nperm)  #store permuted results
  
  for (i in 1:nperm) {
    perm_trait <- fastBM(phy, sig2 = var(trait), bounds = c(0,1))  #simulate total count CV under BM (not ideal, but hey ho). Bounds set to 0,1 to ensure that negative values are not estimated (not possible)
    perm_kw[i] <- kruskal.test(perm_trait ~ group)$statistic
  }
  
  #compute p-value
  p_value <- mean(perm_kw >= observed_kw) #how many times is chi-squared of permutation larger than observed?
  
  #return both observed statistic and permutation results
  return(list(observed_statistic = observed_kw, #returned chi-squared for observed 
              p_value = p_value, #return the p-value
              null_distribution = perm_kw)) #return all calculated chi-squared values for each permutation
}
```

```{r Kruskal-Wallis Test (Total Count CV ~ Total Count) - Phylogenetic Permutation}

set.seed(1931)
#run test
result <- phylo_kruskal(vertebral_counts_intraspecific_pruned$total_count_cv, 
                        vertebral_counts_intraspecific_pruned$total_count_bin, 
                        mcgee_intraspecific_pruned_tree)

#output result
result

#define variables for plotting etc.
null_distribution <- result$null_distribution
observed_statistic <- result$observed_statistic #Kruskal-Wallis chi-squared 2.3657 
p_value <- result$p_value
```

```{r Plot Distribution of Chi-Squared Values From Simulations}

#define chi_sq values from the null distribution as a dataframe
null_distribution <- data.frame(chi_square = null_distribution)

#plot the null distribution of then chi-squared values
b2 <- ggplot(null_distribution, aes(x=chi_square))+
  geom_histogram(aes(y = ..density..), bins = 20, fill = "lightgray", color = "black", alpha = 0.7)+
  ylab("Density")+
  xlab(expression(bold("Kruskal-Wallis Statistic (") * bold(chi^2) * bold(")"))) +
  geom_vline(xintercept = 2.3657, color = "red", linetype='dashed', size = 1.2)+
  theme_classic()+
  theme(legend.position='none',
                 axis.text = element_text(size=11),
                 axis.title = element_text(size=11, face='bold'))
b2

ggsave("Figures/Somitogenesis and Regionalisation/simulated_chi_sq_distribution_total_count_cv_total_count.jpeg", b2, dpi = 600, width = 10, height = 10, units = "cm")
```

```{r Fitting Individual Species BAR ~ TC Linear Models}

#fit linear models for each species with at least 10 individuals, note that for some of them the vertebral counts are the same between individuals and will therefore be undefined.
within_species_linear_models <- vertebral_counts %>%
  group_by(species_name) %>%
  filter(n() >= 10) %>%
  group_split() %>%
  map_df(~ {
    mod <- tryCatch(
      lm(ln_length_ln_width ~ log(total_count), data = .x),
      error = function(e) NULL
    )
    
    if (is.null(mod)) {
      return(tibble(
        species_name = unique(.x$species_name),
        slope = NA_real_,
        slope_se = NA_real_,
        p_value = NA_real_,
        r_squared = NA_real_,
        n = nrow(.x)
      ))
    }
    
    coefs <- summary(mod)$coefficients
    if (nrow(coefs) < 2) {
      return(tibble(
        species_name = unique(.x$species_name),
        slope = NA_real_,
        slope_se = NA_real_,     
        p_value = NA_real_,
        r_squared = NA_real_,
        n = nrow(.x)
      ))
    }

    tibble( 
      species_name = unique(.x$species_name),
      slope = coefs[2, 1],
      slope_se = coefs[2, 2],   
      p_value = coefs[2, 4],
      r_squared = summary(mod)$r.squared,
      n = nrow(.x)
    )
  })

#remove NA values (i.e. species where no intraspecific variation is present)
within_species_linear_models <- na.omit(within_species_linear_models)

#prune the mcgee 2020 tree to data
within_species_linear_models_pruned_tree <- drop.tip(mcgee_2020, 
                                            which(!mcgee_2020$tip.label %in% within_species_linear_models$species_name))

#subset data frame to include only species present in the pruned tree
#subset according to the phylogeny
within_species_linear_models_subsetted <- within_species_linear_models[
 within_species_linear_models$species_name %in% within_species_linear_models_pruned_tree$tip.label, 
]
#convert to dataframe
within_species_linear_models_subsetted <- as.data.frame(within_species_linear_models_subsetted)
#set rownames as species_names for ease later
rownames(within_species_linear_models_subsetted) <- within_species_linear_models_subsetted$species_name

#reorder data frame based on the order of tip labels in the pruned tree
within_species_linear_models_subsetted <- within_species_linear_models_subsetted[
  match(within_species_linear_models_pruned_tree$tip.label, within_species_linear_models_subsetted$species_name), 
]
```

```{r Do Small Sample Sizes Increase Slope Noise?}
ggplot(within_species_linear_models_subsetted, aes(x=n, y=slope))+
  geom_point()+
  geom_smooth(method = "lm")+
  theme_classic()
```

```{r Construct VCV Matrix for Downstream Analysis}
#construct the vcv matrix:
slopes_species <- caper::comparative.data(
  phy = within_species_linear_models_pruned_tree,
  data = within_species_linear_models_subsetted,
  names.col = species_name,
  vcv = TRUE,
  vcv.dim = 3
)
```

```{r Phylogenetic Meta-Analysis on Slopes}
#convert species_name to factor with levels matching tree tip order
within_species_linear_models_subsetted$species_name <- factor(within_species_linear_models_subsetted$species_name, levels = within_species_linear_models_pruned_tree$tip.label)

#calculate phylogenetic correlation matrix
phy_cor <- ape::vcv(phy, corr = TRUE)
phy_cor <- phy_cor[within_species_linear_models_subsetted$species_name, 
                   within_species_linear_models_subsetted$species_name]

#run phylogenetic meta-analysis weighted by slope SE, accounting for phylogeny
res <- rma.mv(
  yi = within_species_linear_models_subsetted$slope, #use slopes from each species’ linear model as effect sizes
  V = within_species_linear_models_subsetted$slope_se^2, #weight each slope by the SAMPLING variance of the slope estimate (se^2)
  random = ~ 1 | species_name, #account for species-level variation, allow each species to have its own deviation from overall mean slope
 #this models species-specific effects as random deviations sampled from a shared distribution
  R = list(species_name = phy_cor), #incorporate phylogenetic relationships by including the phylogenetic covariance matrix
  method = "REML", #estimate parameters using restricted maximum likelihood (REML)
  data = within_species_linear_models_subsetted
)
#summarise the model
summary(res)
```

```{r Plot Slope Sampling Variances}

#add significance column based on p-value
within_species_linear_models_subsetted$significant <- ifelse(
  within_species_linear_models_subsetted$p_value < 0.05, "Significant", "Not Significant"
)

#plot sampling variances and scale points according to weights used in model fit (i.e. 1/se^2)
help <- ggplot(within_species_linear_models_subsetted, aes(x = log10(slope_se^2), y = slope, colour = significant, shape=significant))+
  geom_point(aes(size = 1/slope_se^2), alpha=0.50)+ #weighted variance (1/sampling variance) - i.e. the weights used
  xlab(expression(bold("Log"[10]*" of Slope SE"^2*" (Sampling Variance)")))+
  ylab(expression(bold("Slope (" * beta[1] * ")")))+
  geom_hline(yintercept = 0, linetype='dotdash', colour='blue', linewidth=1)+
  scale_colour_manual(values = c("Significant" = "red", "Not Significant" = "black"))+
  theme_classic()+
  theme(legend.position='none',
                 axis.text = element_text(size=11),
                 axis.title = element_text(size=11, face='bold'))
help

ggsave("Figures/Somitogenesis and Regionalisation/slopes_versus_slope_sampling_variance.jpeg", help, dpi = 600, width = 10, height = 10, units = "cm")

```

```{r Plot Distribution of Slopes}

#extract fixed effect (i.e. intercept of model) and standard error from REML model
fixed_effect <- coef(res)[1] #note only fit intercept-only model so only one coefficient estimated
se <- res$se #standard error for the slope

#calculate 95% confidence interval manually
lower_ci <- fixed_effect - 1.96 * se
upper_ci <- fixed_effect + 1.96 * se

#plot histogram with both REML and raw mean + CI lines
help_2 <- ggplot(within_species_linear_models_subsetted, aes(x = slope))+
  geom_histogram(aes(y = ..density..), bins = 20, fill = "lightgray", color = "black", alpha = 0.7)+
  ylab("Density")+
  xlab(expression(bold("Slope (" * beta[1] * ")")))+
  #REML mean and CI lines (red)
  geom_vline(xintercept = fixed_effect, colour = "red", linetype = "solid", size = 1)+
  geom_vline(xintercept = lower_ci, colour = "red", linetype = "dashed", size = 0.8)+
  geom_vline(xintercept = upper_ci, colour = "red", linetype = "dashed", size = 0.8)+
  theme_classic()+
  theme(legend.position='none',
                 axis.text = element_text(size=11),
                 axis.title = element_text(size=11, face='bold'))
help_2

ggsave("Figures/Somitogenesis and Regionalisation/slopes_distribution_metaanalysis_ci.jpeg", help_2, dpi = 600, width = 10, height = 10, units = "cm")

```
```{r Filter to Species with 25 (or more) Individuals}
#filter for species which have 25 or more individual specimens
individual_counts_25_above <-
  vertebral_counts_intraspecific_variation_filtered %>%
  filter(n_count >= 25) #25 gives a nice range of species (n=17), some very elongate or deep bodied

#identify species
species_lots_individuals <- individual_counts_25_above$species_name

#filter vertebral counts data (i.e. individuals) to species present defined above
vertebral_counts_individual_counts_25_above <- 
  vertebral_counts %>%
  filter(species_name %in% species_lots_individuals)

#check that it has worked
all(sort(unique(vertebral_counts_individual_counts_25_above$species_name)) == sort(species_lots_individuals))

#filter NA (i.e. missing aspect ratio)
vertebral_counts_individual_counts_25_above <- 
  vertebral_counts_individual_counts_25_above %>%
  drop_na()

#calculate mean and standard error
vertebral_counts_individual_counts_25_above_summary <- 
  vertebral_counts_individual_counts_25_above %>%
  group_by(species_name, total_count) %>%
  summarise(
    mean_aspect_ratio = mean(aspect_ratio, na.rm = TRUE),
    sd_aspect_ratio = ifelse(n() > 1, sd(aspect_ratio, na.rm = TRUE), 0),  # Set SD to 0 if only 1 entry
    n = sum(!is.na(aspect_ratio)),  # Ensure n counts only valid values
    se_aspect_ratio = ifelse(n > 1, sd_aspect_ratio / sqrt(n), 0)  # Set SE to 0 if SD is 0
  )
vertebral_counts_individual_counts_25_above_summary
```

```{r Check with Astatotilapia calliptera first}

#filter for species == A. callitpera
astatotilapia_calliptera <- vertebral_counts_individual_counts_25_above_summary %>%
  filter(species_name == 'Astatotilapia calliptera')

#for jitter:
astatotilapia_calliptera_all <- 
  vertebral_counts_individual_counts_25_above %>%
  filter(species_name == 'Astatotilapia calliptera')

ggplot(astatotilapia_calliptera, aes(x = total_count, y = mean_aspect_ratio, group = species_name))+
  geom_jitter(data=astatotilapia_calliptera_all, aes(x=total_count, y=aspect_ratio), width=0.00, alpha=0.25)+
  geom_point(colour='red', size=3)+
  geom_line(aes(group = species_name), size=0.50)+
  geom_errorbar(aes(ymin = mean_aspect_ratio - sd_aspect_ratio, 
                    ymax = mean_aspect_ratio + sd_aspect_ratio), 
                width = 0.25, alpha = 1, colour = "black")+
  xlab("Total Vertebral Count")+
  #ylab(expression(bold(paste("Body Aspect Ratio (", mu, ")"))))+
  ylab("Body Aspect Ratio")+
  theme_classic()+
          theme(legend.position='none',
                 axis.text = element_text(size=11),
                 axis.title = element_text(size=11, face='bold'))
```

```{r Plot All Species >25 individuals (TC versus AR)}

vertebral_counts_individual_counts_25_above_summary <- vertebral_counts_individual_counts_25_above_summary %>%
  mutate(aspect_ratio_range = cut(mean_aspect_ratio, 
                                  breaks = c(2, 4, 6, 8, 10), labels = c("Deep Bodied", "Medium Bodied", "Elongate", "Elongate")))

#random colour generator
set.seed(1971)
random_colours <- sample(colours(), 17)

#plot the mean aspect ratio for vertebral count for each species with error bar
ggplot(vertebral_counts_individual_counts_25_above_summary, 
       aes(x = total_count, y = mean_aspect_ratio, color = species_name, group = species_name)) +
  geom_point(size = 1)+
  geom_line(size = 0.50)+
  geom_errorbar(aes(ymin = mean_aspect_ratio - sd_aspect_ratio, 
                    ymax = mean_aspect_ratio + sd_aspect_ratio), 
                width = 0.25, alpha = 1) +
  facet_wrap(~ aspect_ratio_range, scales = "free_y") +  #Facet by aspect range
  scale_colour_manual(values=c(random_colours))+
  xlab("Total Vertebral Count") +
  ylab("Body Aspect Ratio") +
  theme_classic()+
  theme(legend.position = 'right')

```

```{r Plot All Species >25 individuals (TC versus AR)}
#group body aspect ratio for facet wrapping
vertebral_counts_individual_counts_25_above_summary <- vertebral_counts_individual_counts_25_above_summary %>%
  mutate(aspect_ratio_range = cut(mean_aspect_ratio, 
                                  breaks = c(2, 3.6, 5.5, 8, 10), labels = c("Deep Bodied", "Medium Bodied", "Elongate", "Very Elongate")))

#select species to plot so its less messy
species_to_retain <- c("Teleogramma depressum", "Teleogramma gracile", 
                       "Cyprichromis microlepidotus", "Steatocranus rouxi", 
                       "Serranochromis macrocephalus", "Astatotilapia calliptera", 
                       "Gobiocichla wonderi", 
                       "Pseudocrenilabrus multicolor") #"Steatocranus gibbiceps",

#filter dataframe for the species to plot
vertebral_counts_individual_counts_25_above_summary_filtered <- 
  vertebral_counts_individual_counts_25_above_summary %>%
  filter(species_name %in% species_to_retain)

#plot the mean aspect ratio for vertebral count for each species with error bar and legend
a1 <- ggplot(vertebral_counts_individual_counts_25_above_summary_filtered, 
       aes(x = total_count, y = mean_aspect_ratio, color = species_name, group = species_name)) +
  geom_point(size = 1.5)+
  geom_line(size = 0.50)+
  geom_errorbar(aes(ymin = mean_aspect_ratio - se_aspect_ratio, 
                    ymax = mean_aspect_ratio + se_aspect_ratio), 
                width = 0.50, alpha = 0.75)+
  facet_wrap(~ aspect_ratio_range, scales = "free_y")+  #Facet by aspect range
  xlab("Total Vertebral Count")+
  ylab("Body Aspect Ratio")+
  scale_x_continuous(breaks = seq(26, 39, by = 4)) +  # Keeps tick labels at 26-39
  coord_cartesian(xlim = c(26, 39)) +  # Prevents clipping while keeping axis range
  theme_classic()+
  theme(axis.text = element_text(size=11),
                 axis.title = element_text(size=11, face='bold'))
a1

ggsave("Figures/Somitogenesis and Regionalisation/total_count_by_aspect_ratio_full.pdf", a1, dpi = 600, width = 10, height = 10, units = "cm")

```

```{r Plot All Species >25 individuals (TC versus AR) - Clean}
#group body aspect ratio for facet wrapping
vertebral_counts_individual_counts_25_above_summary <- vertebral_counts_individual_counts_25_above_summary %>%
  mutate(aspect_ratio_range = cut(mean_aspect_ratio, 
                                  breaks = c(2, 3.6, 5.5, 8, 10), labels = c("Deep Bodied", "Medium Bodied", "Elongate", "Very Elongate")))

#select species to plot so its less messy
species_to_retain <- c("Teleogramma depressum", "Teleogramma gracile", 
                       "Cyprichromis microlepidotus", "Steatocranus rouxi", 
                       "Serranochromis macrocephalus", "Astatotilapia calliptera", 
                       "Gobiocichla wonderi", 
                       "Pseudocrenilabrus multicolor") #"Steatocranus gibbiceps",

#filter dataframe for the species to plot
vertebral_counts_individual_counts_25_above_summary_filtered <- 
  vertebral_counts_individual_counts_25_above_summary %>%
  filter(species_name %in% species_to_retain)

#for jitter:
total_counts_all <- 
  vertebral_counts_individual_counts_25_above %>%
  filter(species_name %in% species_to_retain)

vertebral_counts_individual_counts_25_above_summary
#assign aspect_ratio_range to total_counts_all
total_counts_all <- total_counts_all %>%
  left_join(dplyr::select(vertebral_counts_individual_counts_25_above_summary, species_name, aspect_ratio_range),
            by = "species_name")
total_counts_all <- total_counts_all %>% distinct()

#plot the mean aspect ratio for vertebral count for each species with error bar (clean)
a2 <- ggplot(vertebral_counts_individual_counts_25_above_summary_filtered, 
       aes(x = total_count, y = mean_aspect_ratio, colour = species_name, group = species_name))+
  geom_jitter(data = total_counts_all, 
              aes(x = total_count, y = aspect_ratio, colour = species_name),  
              width = 0.15, height = 0.02, alpha = 0.75, size=0.75, inherit.aes = FALSE)+
  geom_point(size = 1.5, alpha=0.75, colour='black') +
  geom_line(size = 0.50, alpha=0.75, colour='black') +
  geom_errorbar(aes(ymin = mean_aspect_ratio - sd_aspect_ratio, 
                    ymax = mean_aspect_ratio + sd_aspect_ratio), 
                width = 0.50, alpha = 0.75, colour='black')+
  facet_wrap(~ aspect_ratio_range, scales = "free_y") +  # Facet by aspect range
  xlab("Total Vertebral Count") +
  ylab("Body Aspect Ratio") +
  scale_x_continuous(breaks = seq(26, 39, by = 4)) +  # Keeps tick labels at 26-39
  coord_cartesian(xlim = c(26, 39)) +  # Prevents clipping while keeping axis range
  scale_y_continuous(labels = scales::number_format(accuracy = 0.1))+
  theme_classic() +
  theme(legend.position = 'none',
        axis.text = element_text(size = 11),
        axis.title = element_text(size = 11, face = 'bold'))
a2

ggsave("Figures/Somitogenesis and Regionalisation/total_count_by_aspect_ratio_clean.jpeg", a2, dpi = 600, width = 20, height = 10, units = "cm")
```

```{r Plot Distribution of Total Count CVs}

#plot distribution of total counts (checking for normality)
y2 <- ggplot(vertebral_counts_intraspecific_pruned, aes(x = total_count_cv))+
  geom_histogram(aes(y = ..density..), 
                 bins = 6, fill = "lightgray", color = "black", alpha = 0.7)+
  geom_vline(aes(xintercept = median(total_count_cv)), 
             color = "red", linetype = "dashed", size = 1)+ #add median line
  xlab(expression(bold(paste("Total Vertebral Count CV (", sigma, " / ", mu, ")"))))+
  ylab("Density")+
  theme_classic()+
  theme(legend.position='none',
                 axis.text = element_text(size=11),
                 axis.title = element_text(size=11, face='bold'))
y2

ggsave("Figures/Somitogenesis and Regionalisation/total_count_cv_distribution.jpeg", y2, dpi = 600, width = 10, height = 10, units = "cm")
```

```{r Has Variation Remained Same as Cichlids Diversified?}

##cannot directly ancestrally reconstruct intraspecific variation, have to rely on weaker inferences.
##do basal extant species have same variation as more recently diverged species?

# tree: mcgee_intraspecific_pruned_tree
# species data:  vertebral_counts_intraspecific_pruned

#first calculate lambda and Blombergs Kappa for the Total Count CV:

#convert CV values to a named vector
cv_values <- vertebral_counts_intraspecific_pruned$total_count_cv
names(cv_values) <- vertebral_counts_intraspecific_pruned$species_name

#test for phylogenetic signal
#Blomberg's (2003) K:
K_cv <- phylosig(mcgee_intraspecific_pruned_tree, cv_values, nsim=100000, method = "K", test = TRUE)
K_cv

#Pagel's (1999) Lambda:
lambda_cv <- phylosig(mcgee_intraspecific_pruned_tree, cv_values, nsim=100000, method = "lambda", test = TRUE)
lambda_cv

#plot distributions
plot(K_cv)
plot(lambda_cv)

#phylogenetic signal is indistinguishable from 0. for both K and lambda. 
```

```{r Plot Null Distributions/Likelihood Profiles (Total Count CV)}

#Null Distribution for K
sim_k_total_cv <- as.data.frame(K_cv$sim.K)
observed_k_total_cv <- K_cv$K

observed_k_total_cv

#plot the Null distribution (K)
c1 <- ggplot(sim_k_total_cv, aes(x=K_cv$sim.K))+
  geom_histogram(aes(y = ..count..), bins = 20, fill = "lightgray", color = "black", alpha = 0.7)+
  geom_vline(xintercept = observed_k_total_cv,  color = "red", size = 1, linetype='dashed')+
  ylim(0, 30000)+
  xlab("Simulated K")+
  ylab("Count")+
  theme_classic()+
  theme(legend.position='none',
                 axis.text = element_text(size=11),
                 axis.title = element_text(size=11, face='bold'))
c1 

ggsave("Figures/Somitogenesis and Regionalisation/total_count_cv_k_null.jpeg", c1, dpi = 600, width = 10, height = 10, units = "cm")

#Likelihood profile (Lambda)
jpeg("Figures/Somitogenesis and Regionalisation/total_count_cvlambda_likelihood_profile.jpg", 
     width = 12, height = 12, units = "cm", res = 300, quality = 95)

par(font.lab = 2)  # Make axis labels bold
plot(lambda_cv)

dev.off()
```

```{r Kruskal-Wallis Test (Total Count CV ~ Water System)}
#ordinary Kruskal-Wallis test (test for sig difference in medians):
kruskal.test(total_count_cv ~ water_system, data = vertebral_counts_intraspecific_pruned)

#post-hoc test:
test <- pairwise.wilcox.test(vertebral_counts_intraspecific_pruned$total_count_cv, 
                     vertebral_counts_intraspecific_pruned$water_system, 
                     p.adjust.method = "bonferroni")
test$p.value
```

```{r Kruskal-Wallis Test (Total Count CV ~ Water System) - Phylogenetic Permutation}

set.seed(1931)
#run test
result <- phylo_kruskal(vertebral_counts_intraspecific_pruned$total_count_cv, 
                        vertebral_counts_intraspecific_pruned$water_system, 
                        mcgee_intraspecific_pruned_tree)

#output result
result

#define variables for plotting etc.
null_distribution <- result$null_distribution
observed_statistic <- result$observed_statistic #Kruskal-Wallis chi-squared 13.16999 
p_value <- result$p_value
```

```{r Plot Distribution of Chi-Squared Values From Simulations}

#define chi_sq values from the null distribution as a dataframe
null_distribution <- data.frame(chi_square = null_distribution)

#plot the null distribution of then chi-squared values
b2 <- ggplot(null_distribution, aes(x=chi_square))+
  geom_histogram(aes(y = ..density..), bins = 20, fill = "lightgray", color = "black", alpha = 0.7)+
  ylab("Density")+
  xlab(expression(bold("Kruskal-Wallis Statistic (") * bold(chi^2) * bold(")"))) +
  geom_vline(xintercept = 13.16999, color = "red", linetype='dashed', size = 1.2)+
  theme_classic()+
  theme(legend.position='none',
                 axis.text = element_text(size=11),
                 axis.title = element_text(size=11, face='bold'))
b2

ggsave("Figures/Somitogenesis and Regionalisation/simulated_chi_sq_distribution_total_count_water_system.jpeg", b2, dpi = 600, width = 10, height = 10, units = "cm")
```

```{r Plot CV According to Water System}
#for jitter
set.seed(1931)

#reorder factors for plot into recent --> oldest
#recode the levels of water_system factor
vertebral_counts_intraspecific_pruned$water_system <- dplyr::recode(vertebral_counts_intraspecific_pruned$water_system,
                                                  "Lake Victoria" = "LV",
                                                  "Lake Malawi" = "LM",
                                                  "Lake Tanganyika" = "LT",
                                                  "Riverine" = "RV")

#reordering the levels of water_system
vertebral_counts_intraspecific_pruned$water_system <- factor(vertebral_counts_intraspecific_pruned$water_system, 
                                                  levels = c("LV", "LM", "LT", "RV"))

#colours taken from Wong colour blind palette
water_system_colours <- c("LV" = "#43AA8B", 
                          "LM" = "#D55E00", 
                          "LT" = "#0072B2", 
                          "RV" = "#CC79A7")
  
#plot the CV for each species according to water system occupancy
b1 <- ggplot(vertebral_counts_intraspecific_pruned, aes(x=water_system, y=total_count_cv)) +
  geom_boxplot(outlier.shape = NA, linewidth=0.5) +
  geom_jitter(aes(shape=haplochromine, colour=water_system), 
              size=1.25, width=0.35, alpha=0.75) +
  scale_colour_manual(values = water_system_colours) +
  scale_shape_manual(values = c('Haplochromine' = 19, 'Other' = 17)) +
  xlab('Water System') +
  ylab(expression(bold(paste("Total Vertebral Count CV (", sigma, " / ", mu, ")")))) +
  theme_classic() +
  theme(legend.position='none',
        axis.text = element_text(size=11),
        axis.title = element_text(size=11, face='bold'))

b1

ggsave("Figures/Somitogenesis and Regionalisation/total_count_cv_by_water_system.jpeg", b1, dpi = 600, width = 10, height = 10, units = "cm")
```

```{r Plotting Total Count SD as Contmap on All Species Phylogeny}

#note we are plotting only species retained on tree which have 2 or more specimens/observations

#prepare the matrix:
total_count_cv <- as.vector(vertebral_counts_intraspecific_pruned$total_count_cv)
names(total_count_cv) <- vertebral_counts_intraspecific_pruned$species_name


#Compute 'contmap' object for Total Count CV:
all_species_total_count_cv_contmap <- 
  contMap(mcgee_intraspecific_pruned_tree,
          type="fan",
          total_count_cv,
          plot=FALSE) #can set colour limits using limits=c(min, max) default is the min and max value

#Set custom colour gradient:
set.seed(1931)
colour_gradient <- viridis(n=5) #randomly select 6 evenly spaced colours from viridis colour

all_species_total_count_cv_contmap <- 
  setMap(all_species_total_count_cv_contmap, colour_gradient)

##### Plot total_count SD onto the phylogeny:

pdf("Figures/Somitogenesis and Regionalisation/all_species_total_count_cv_tree.pdf", width=8.4, height=11)

#Plot 'contmap' object:
plot(all_species_total_count_cv_contmap,
     legend=FALSE,
     type = 'fan',
     lwd=2.00,
     fsize = 0.33,
     ftype='b',
     offset = 3.00)
add.color.bar(15, all_species_total_count_cv_contmap$cols,title="Total Count CV", 
    lims=all_species_total_count_cv_contmap$lims,digits=2,prompt=FALSE,x=0,
    y=4.00,lwd=10,fsize=0.55,subtitle="15 Myrs") ##length of legend also scaled to time

dev.off()
```
````
