## Supplementary figures and images for "Somitic Change Drives Changes in Vertebral Regionalisation in African Cichlids Despite Strong Canalisation of Somite Number"

### all_species_total_count_cv_tree.pdf

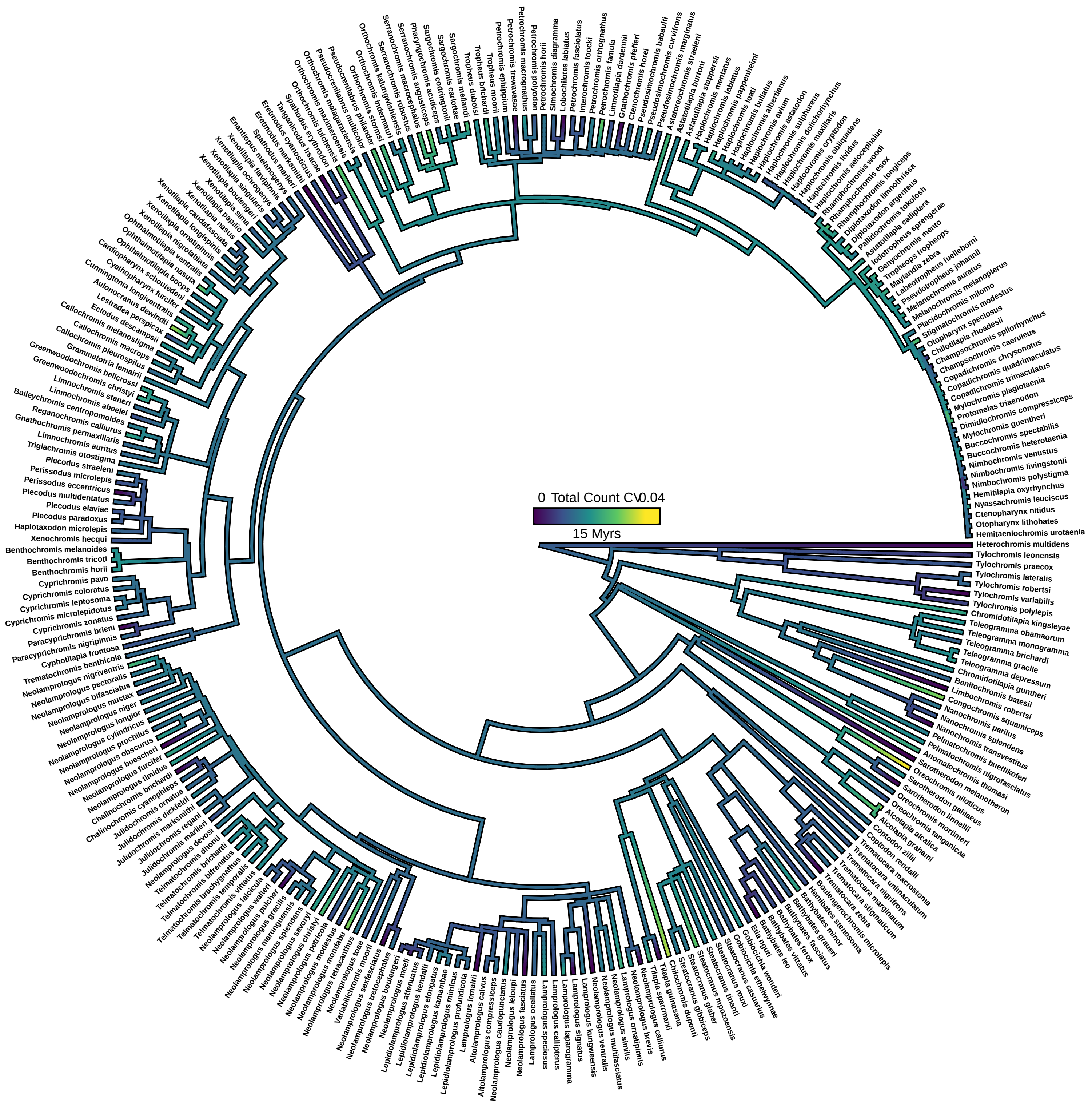

### bar_chart_number_changes.jpeg

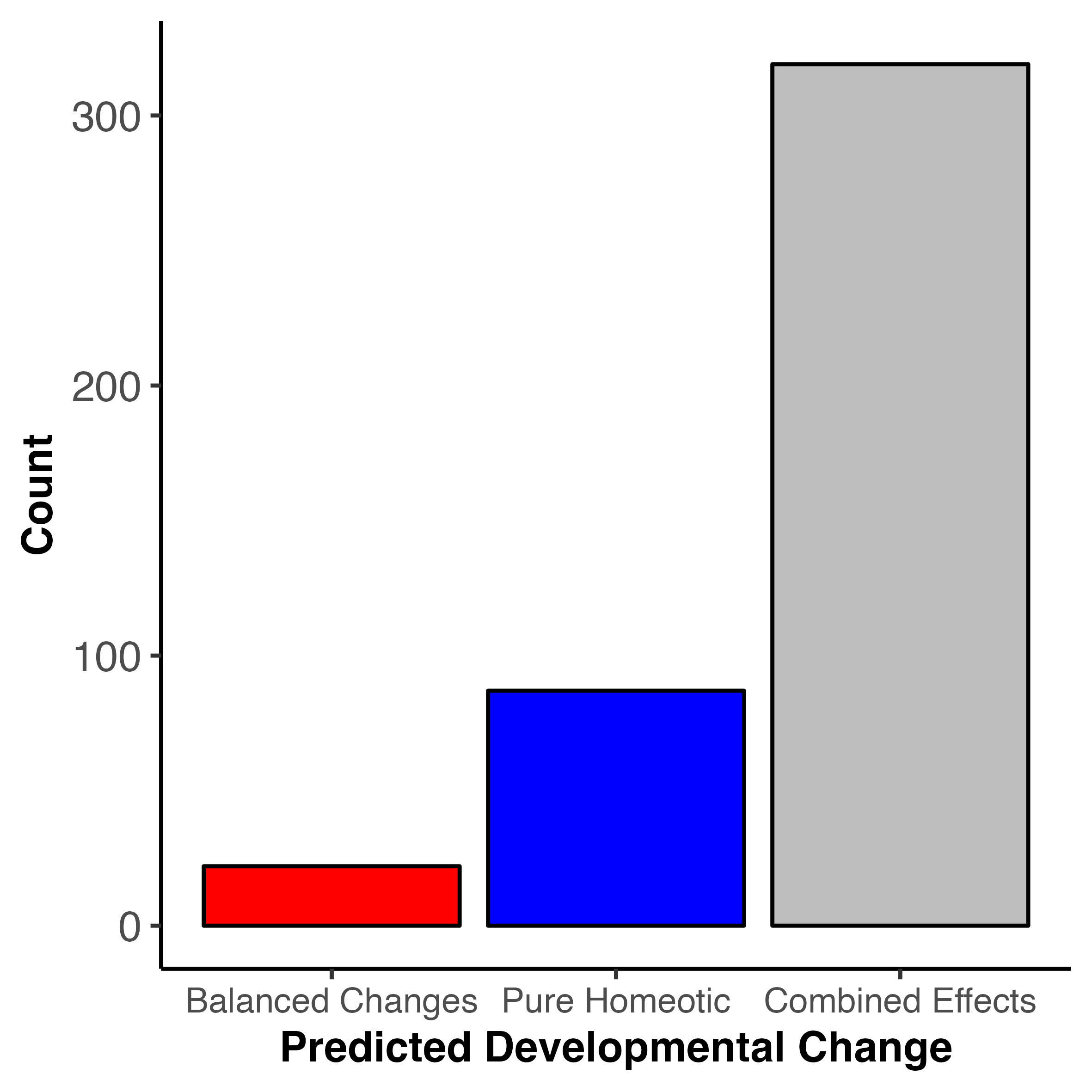

### bar_chart_number_changes_time.jpeg

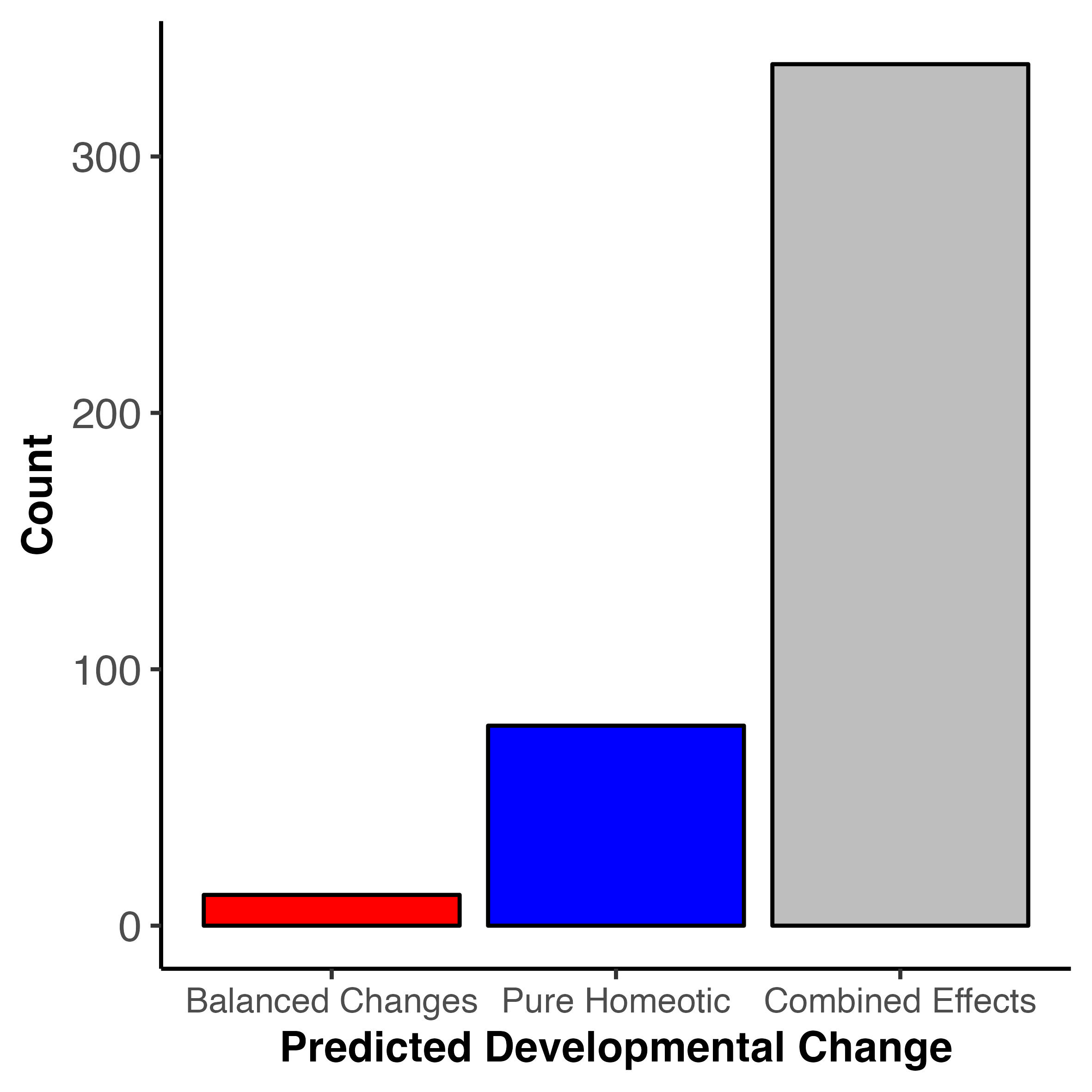

### caudal_cv_k_null.jpeg

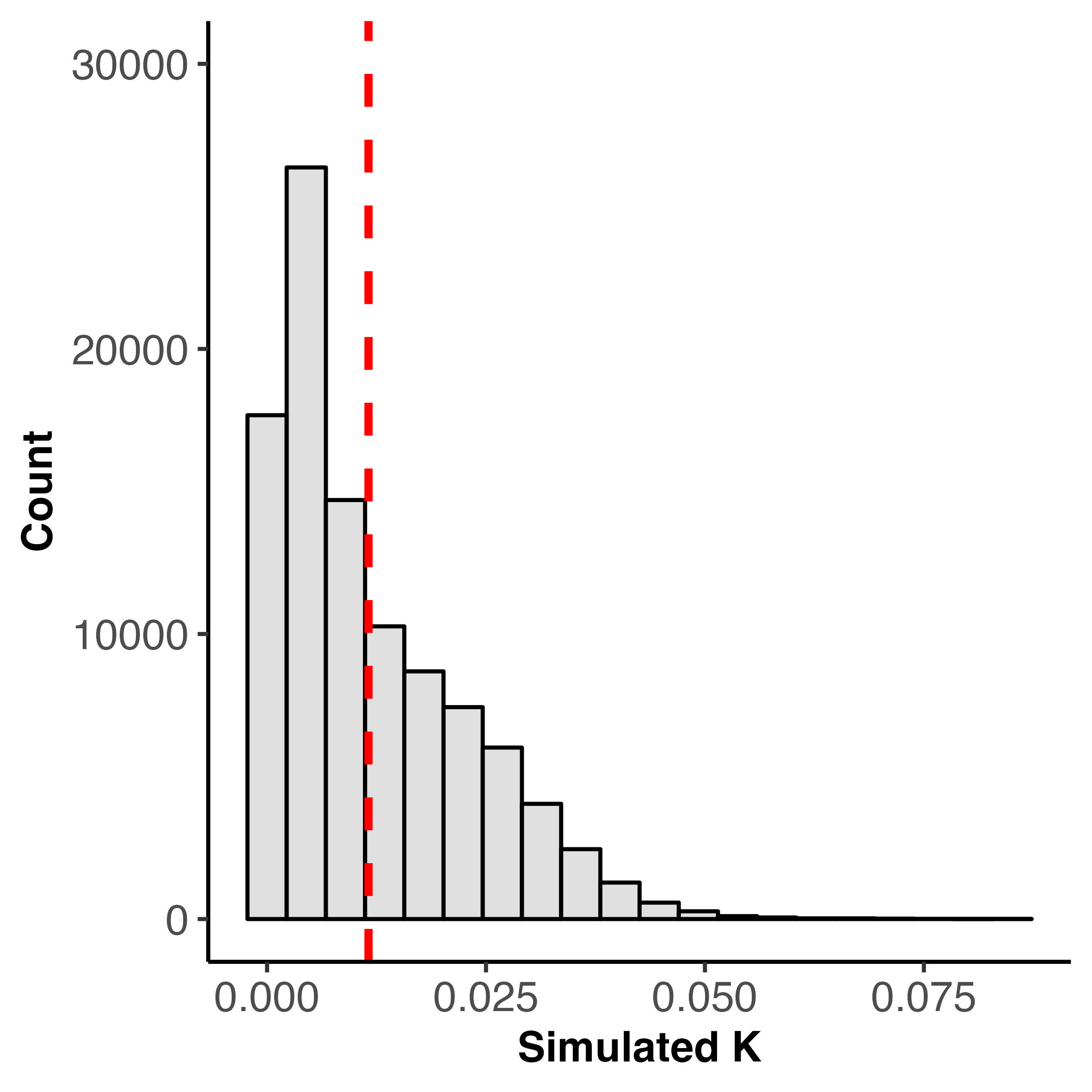

### caudal_cv_lambda_likelihood_profile.jpg

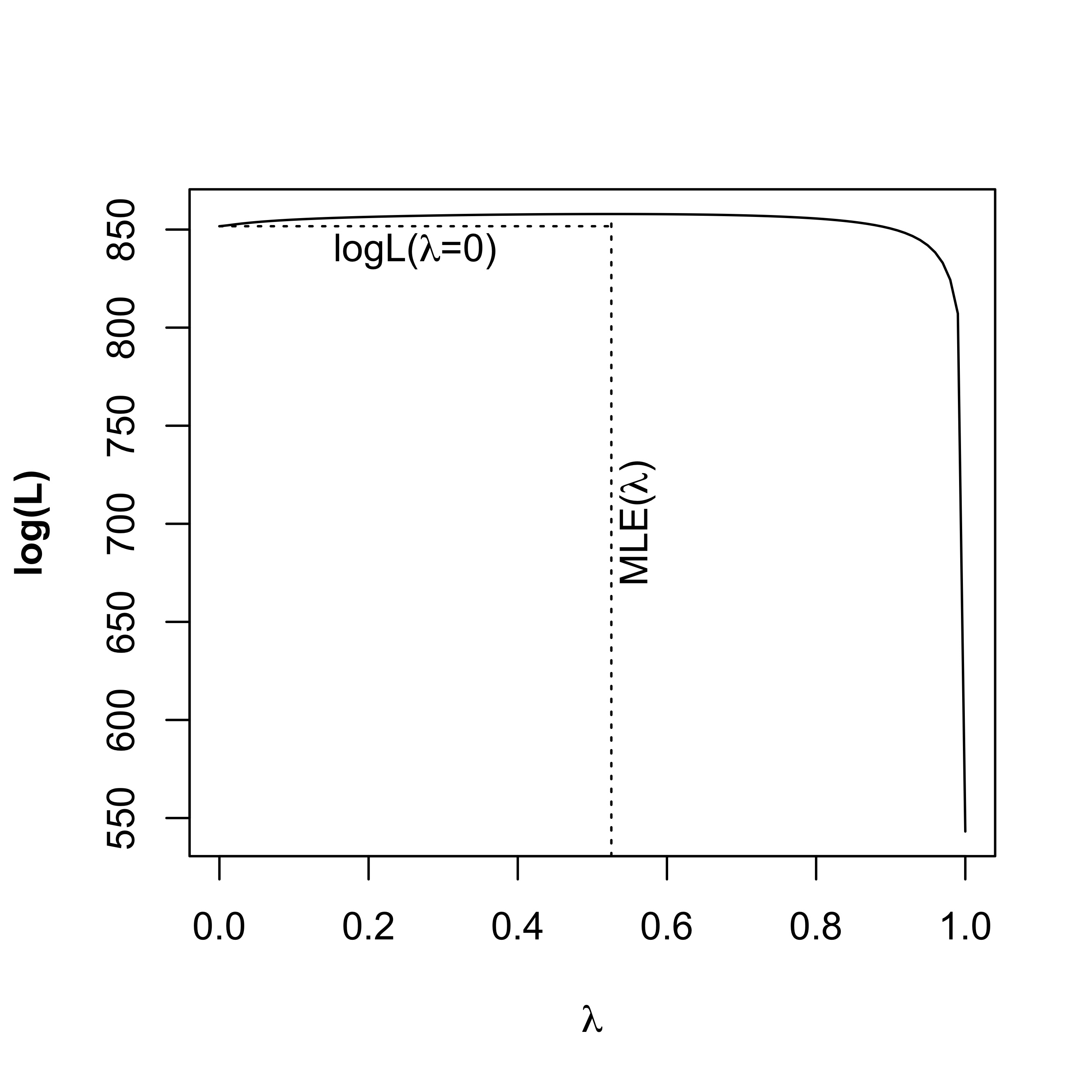

### checking_n_count_variation.jpeg

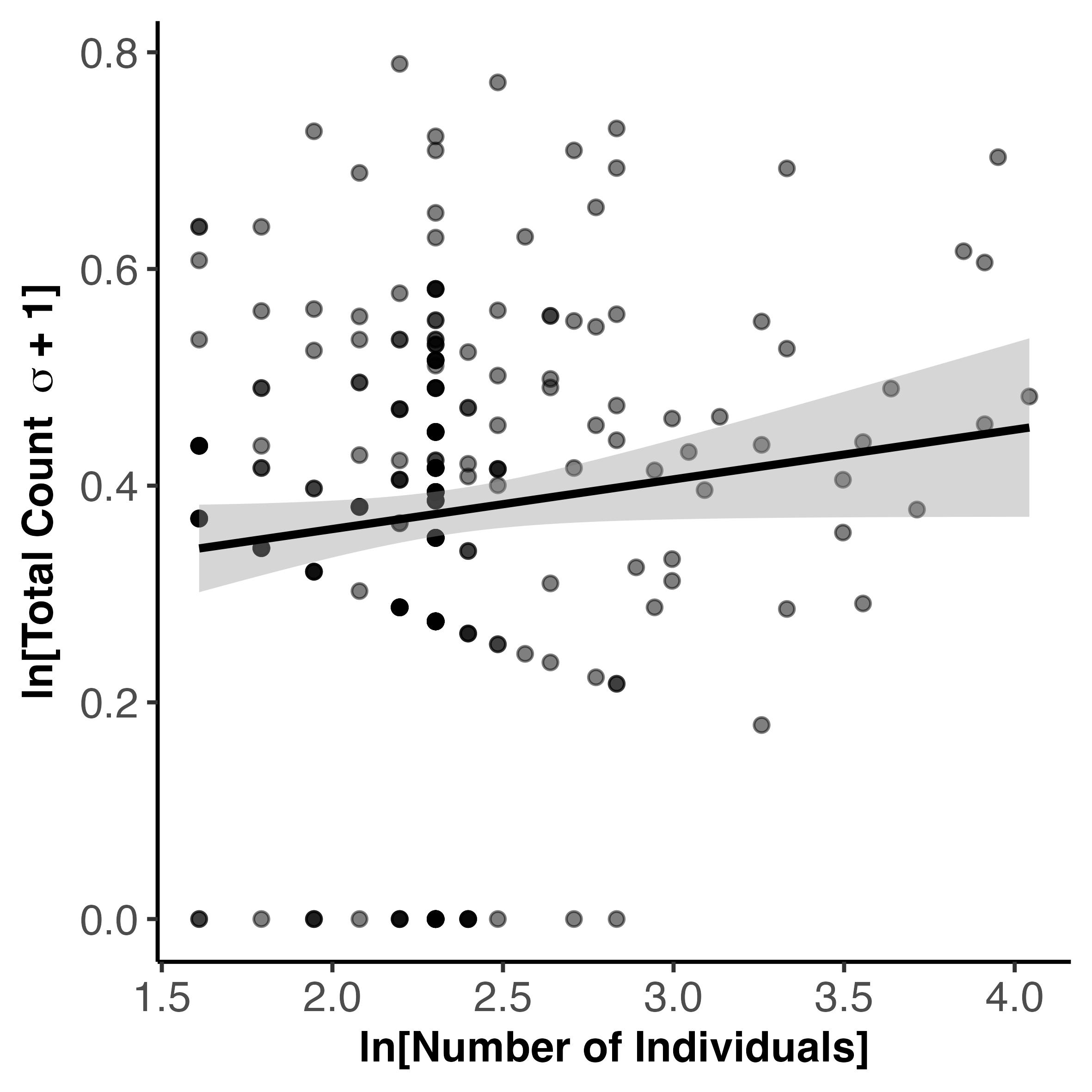

### co_occurring_somitic_changes.pdf

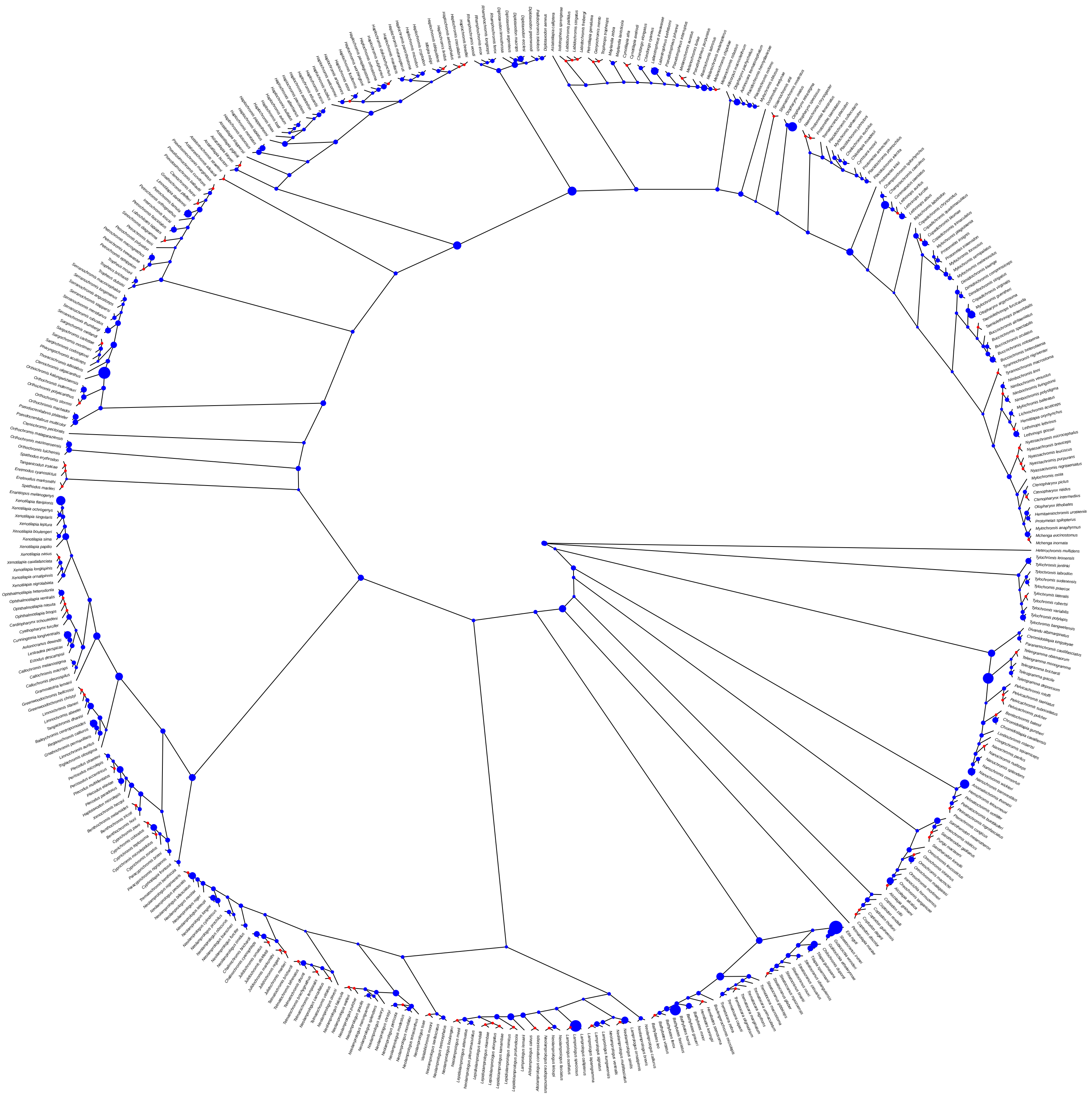

### combined_bar_chart_predicted_dev_change.jpeg

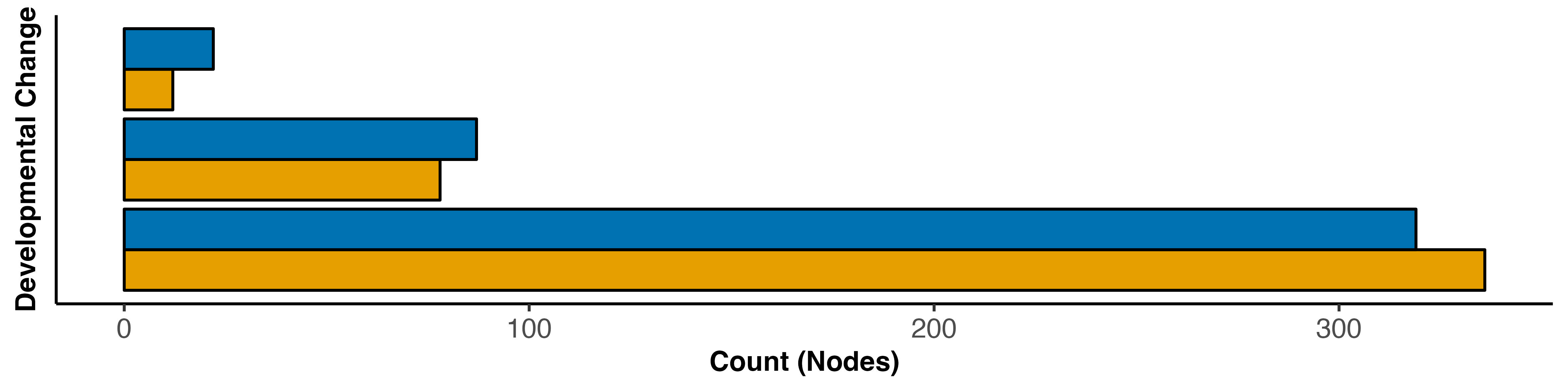

### precaudal_caudal_pgls.jpeg

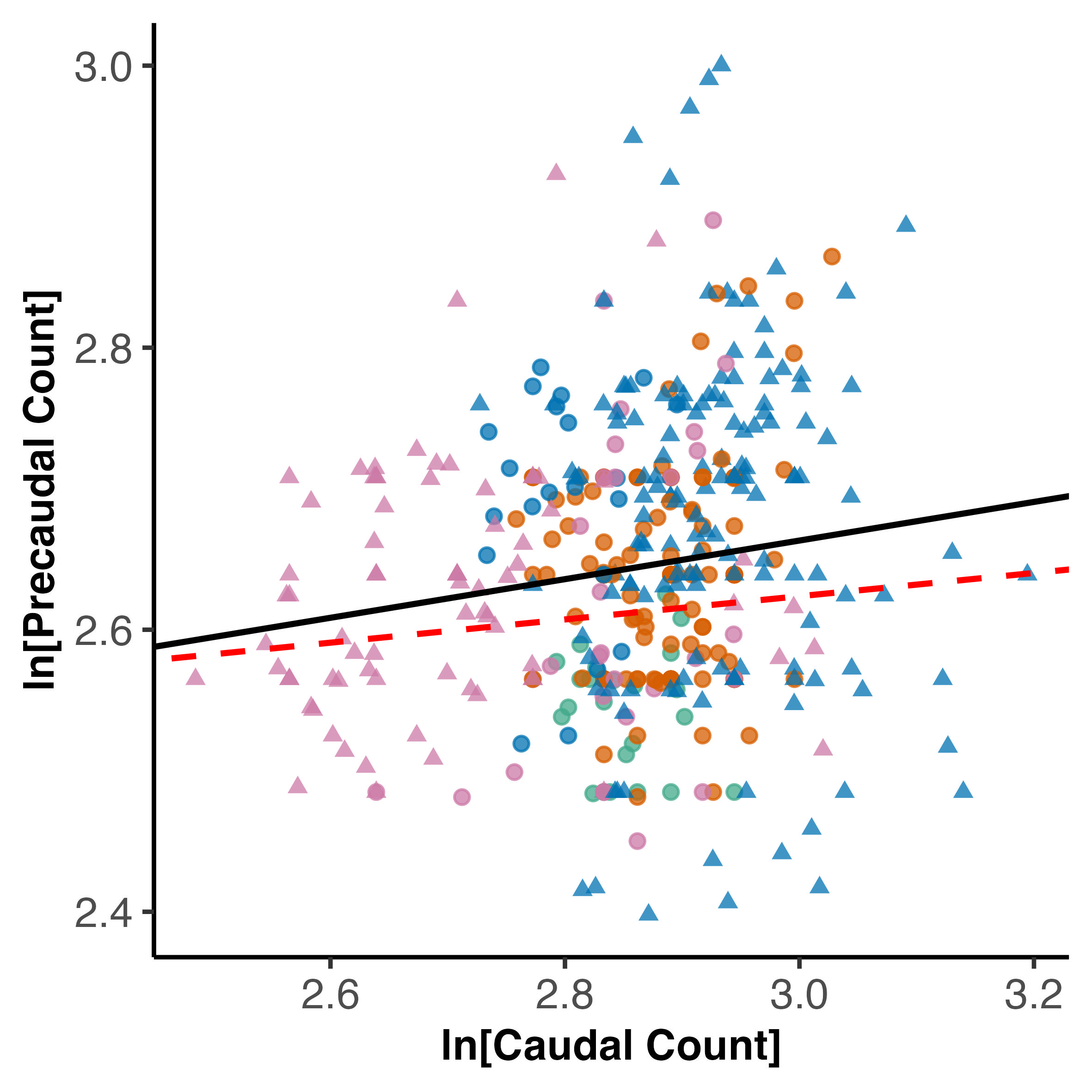

### precaudal_caudal_pgls.pdf

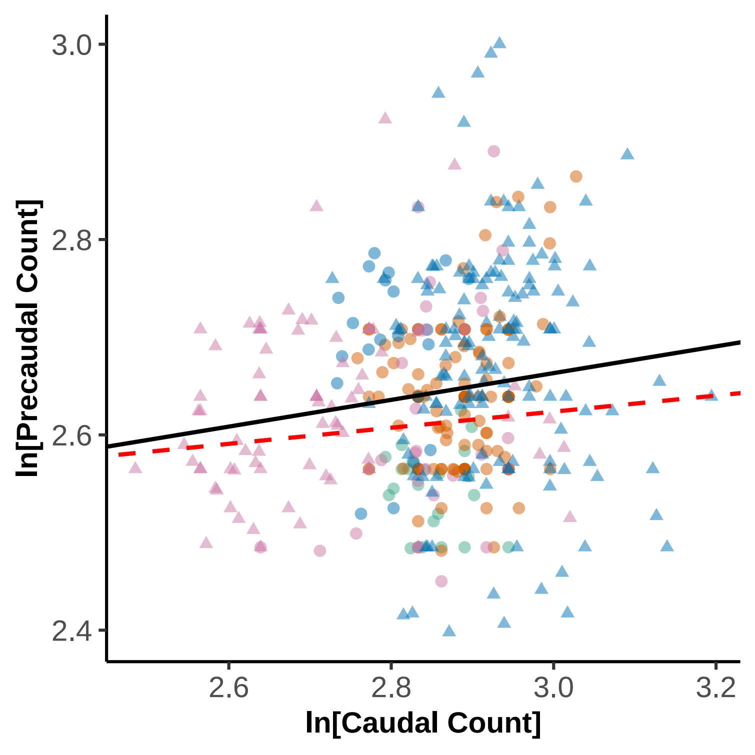

### precaudal_caudal_pics_homeoetic_effects.jpeg

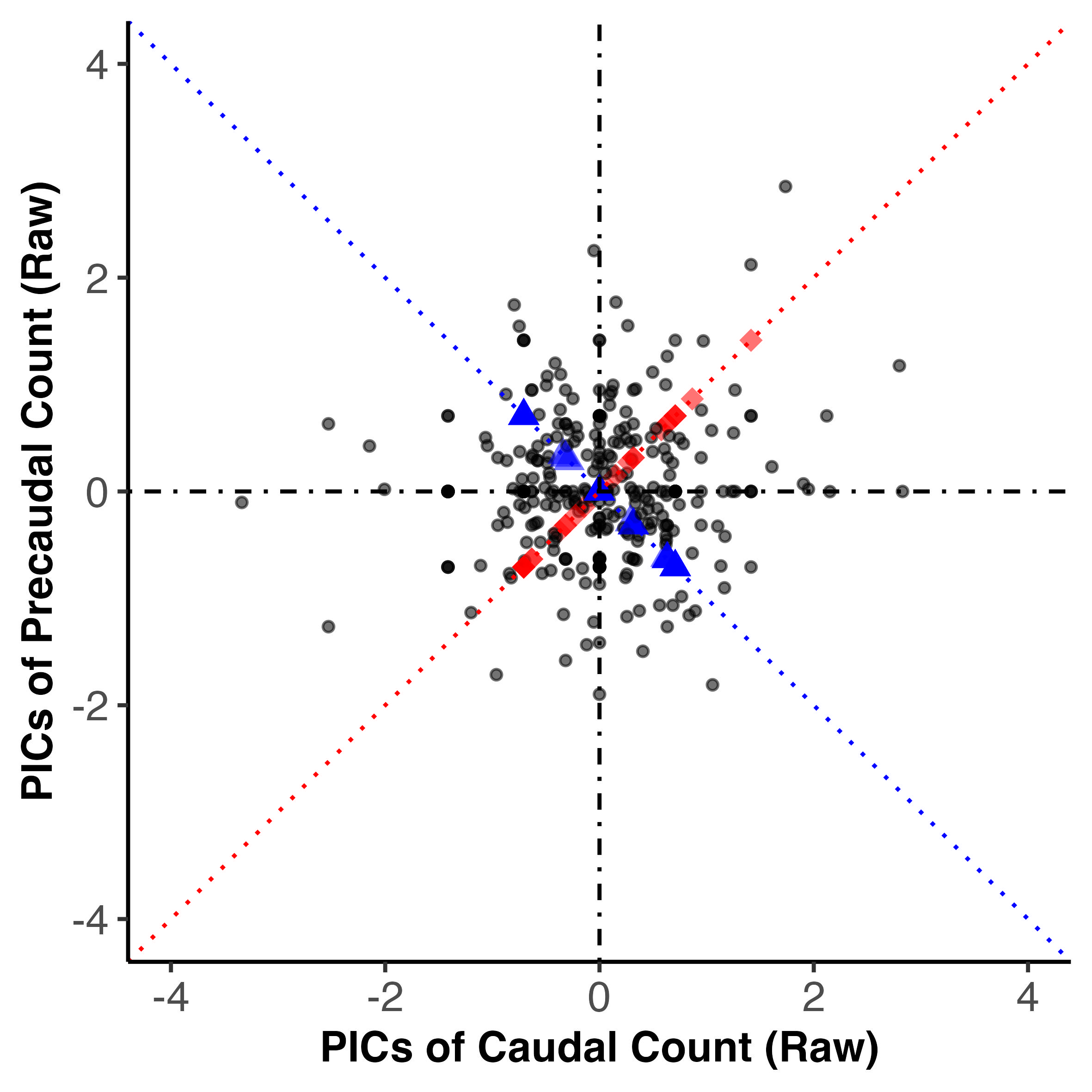

### precaudal_caudal_pics_homeoetic_effects.pdf

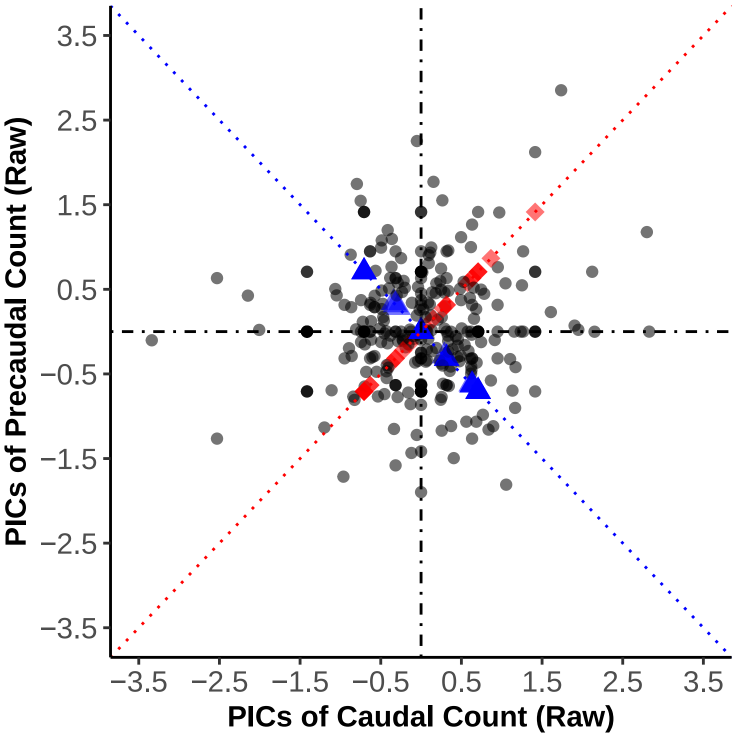

### precaudal_caudal_pics_homeoetic_effects_time_scale.jpeg

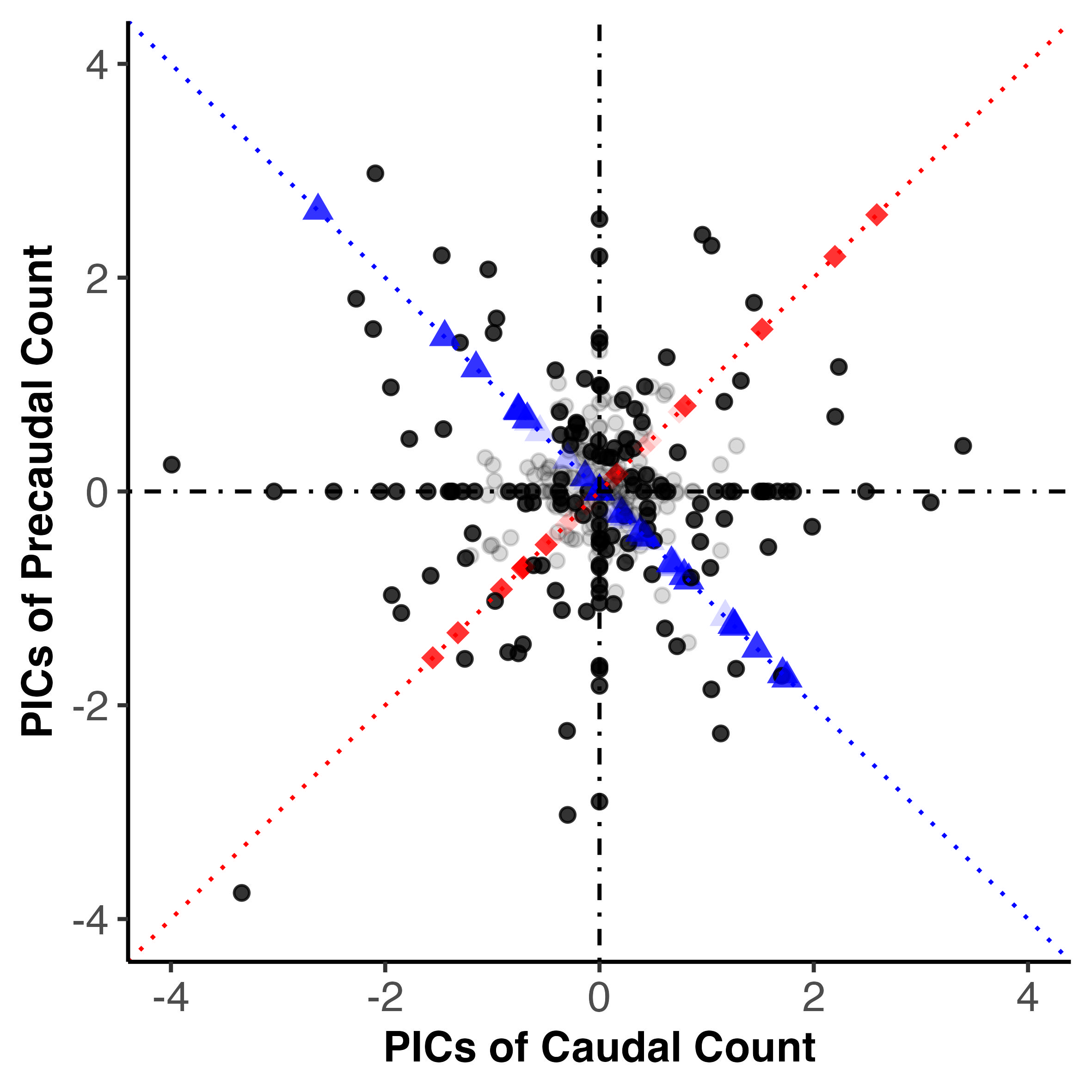

### precaudal_caudal_pics_homeoetic_effects_time_scale.pdf

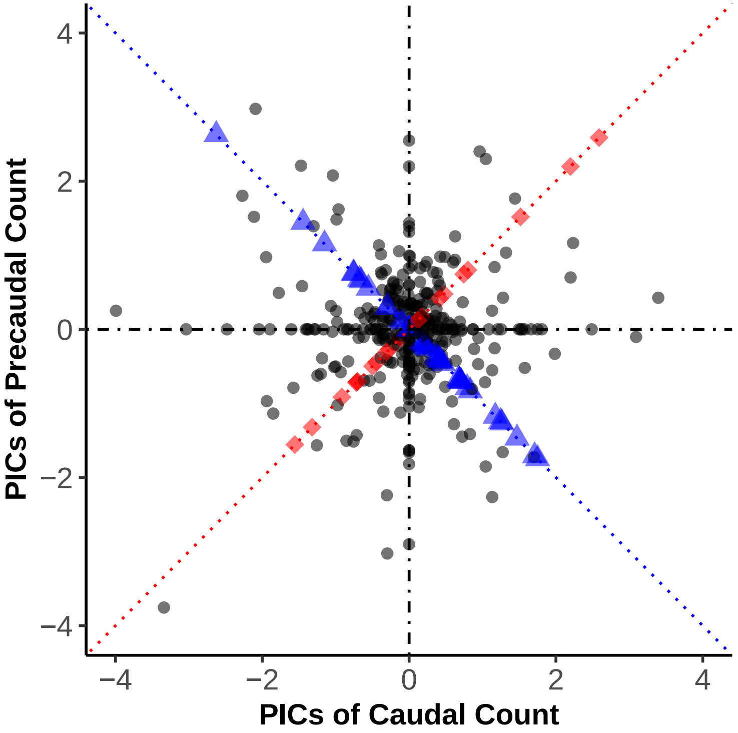

### precaudal_cv_caudal_cv_boxplot.jpeg

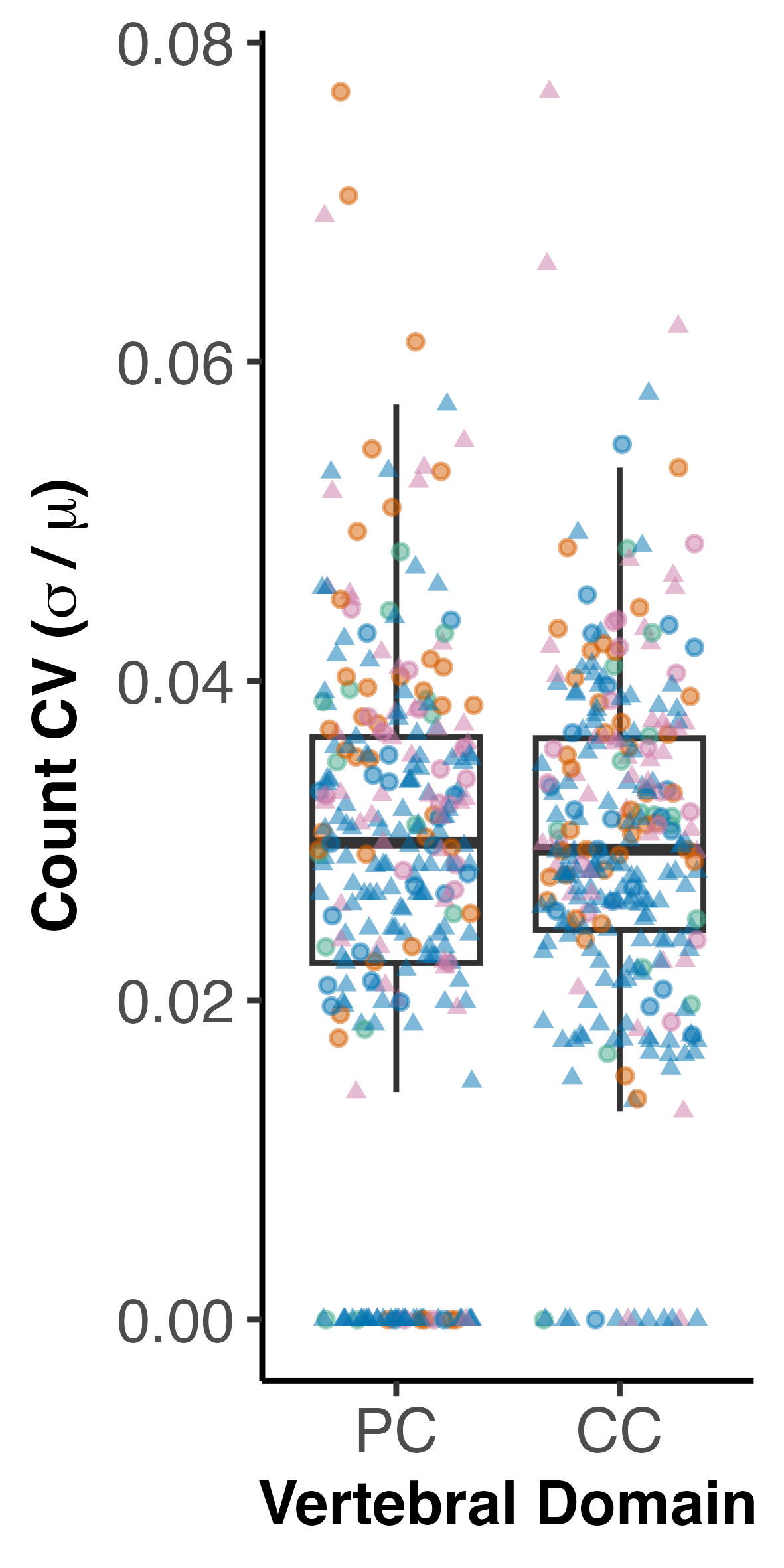

### precaudal_cv_k_null.jpeg

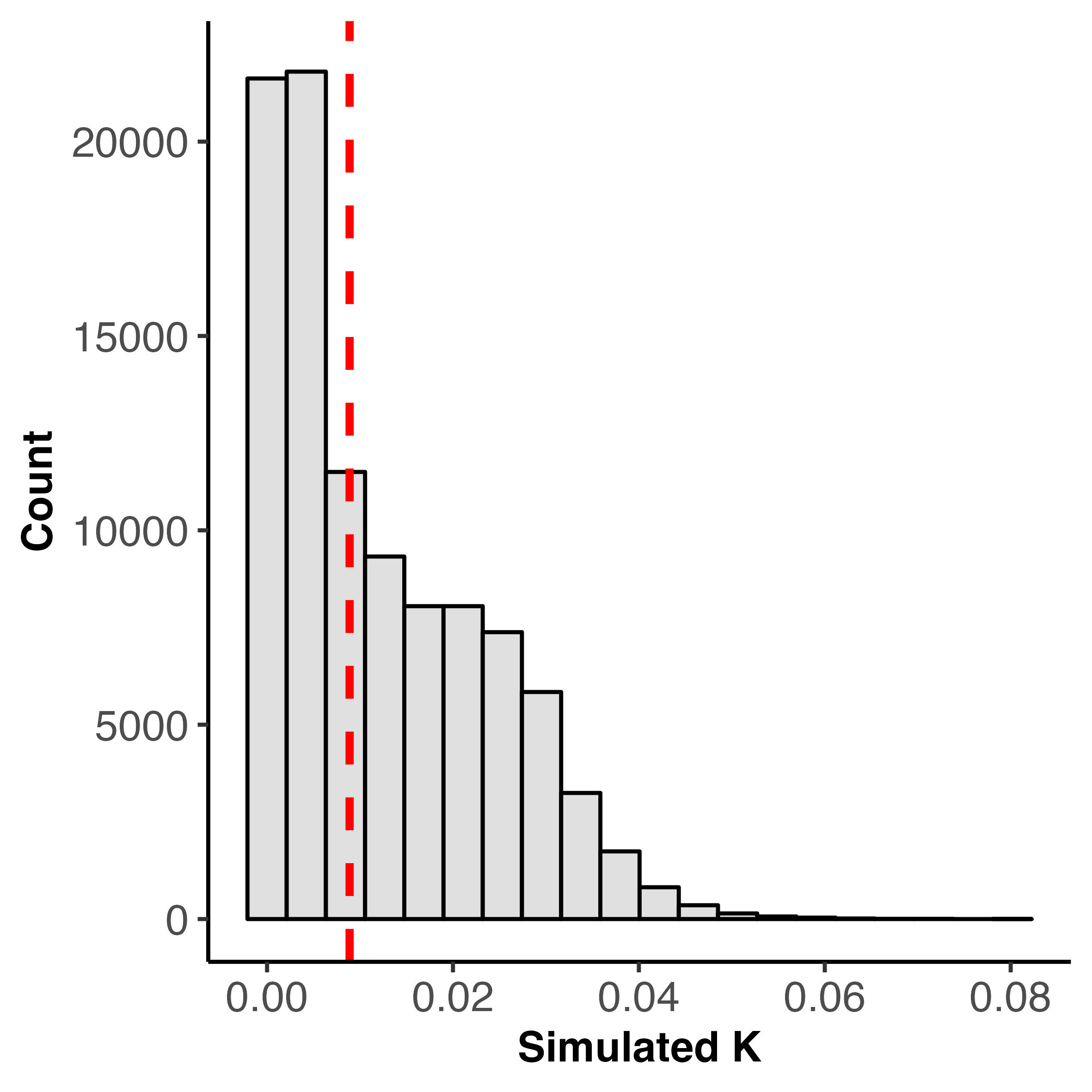

### precaudal_cv_lambda_likelihood_profile.jpg

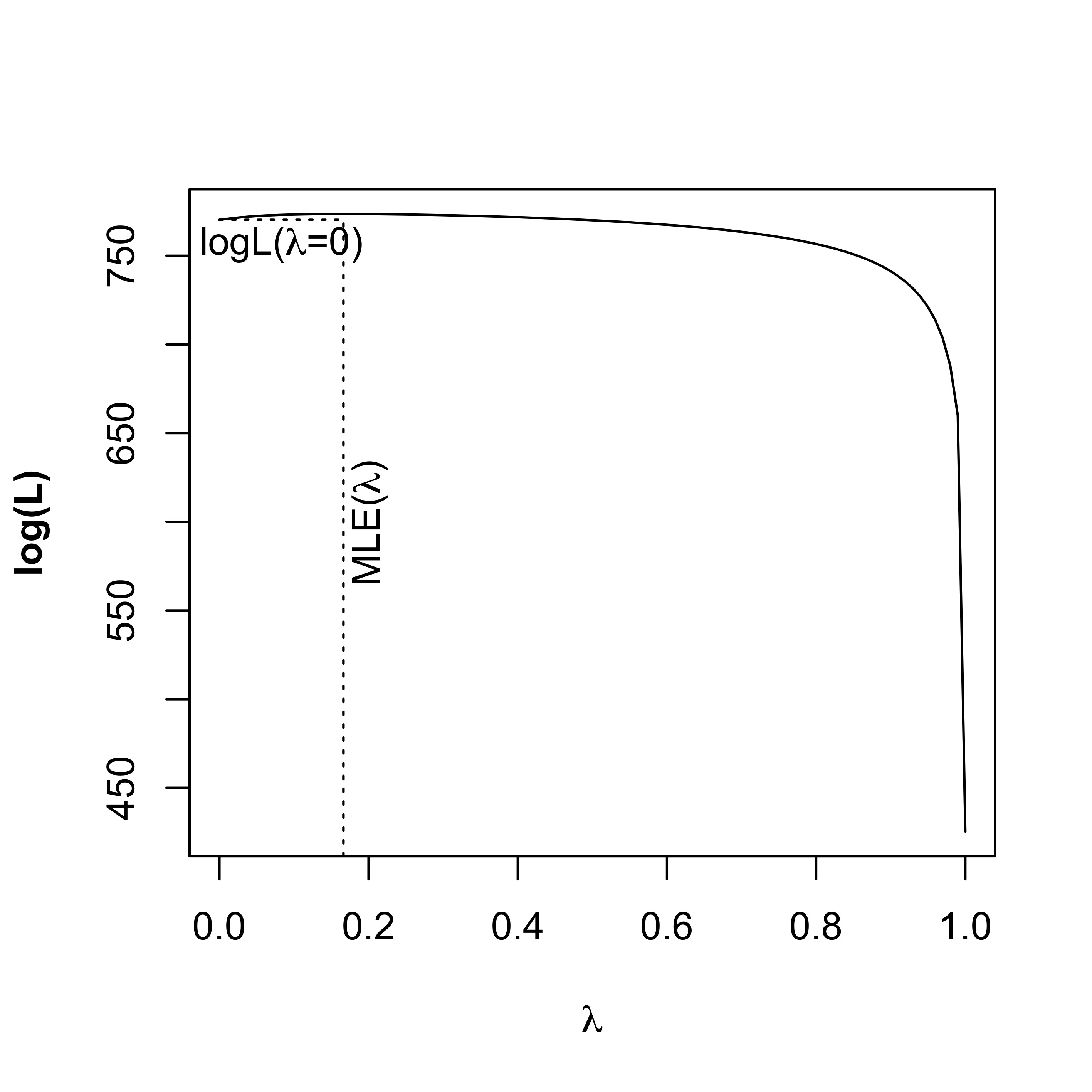

### precaudal_var_caudal_var_boxplot.jpeg

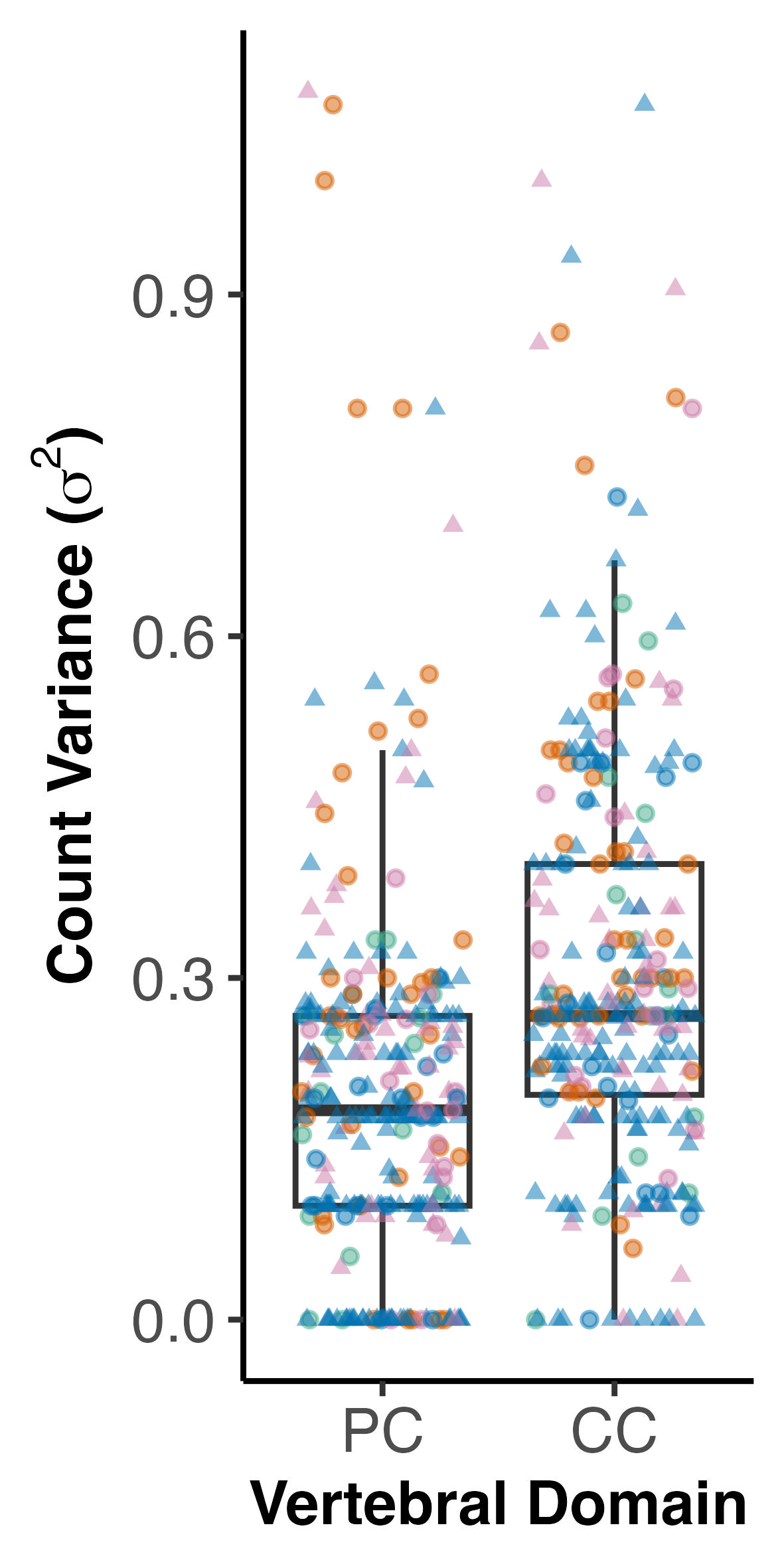

### simulated_chi_sq_distribution.jpeg

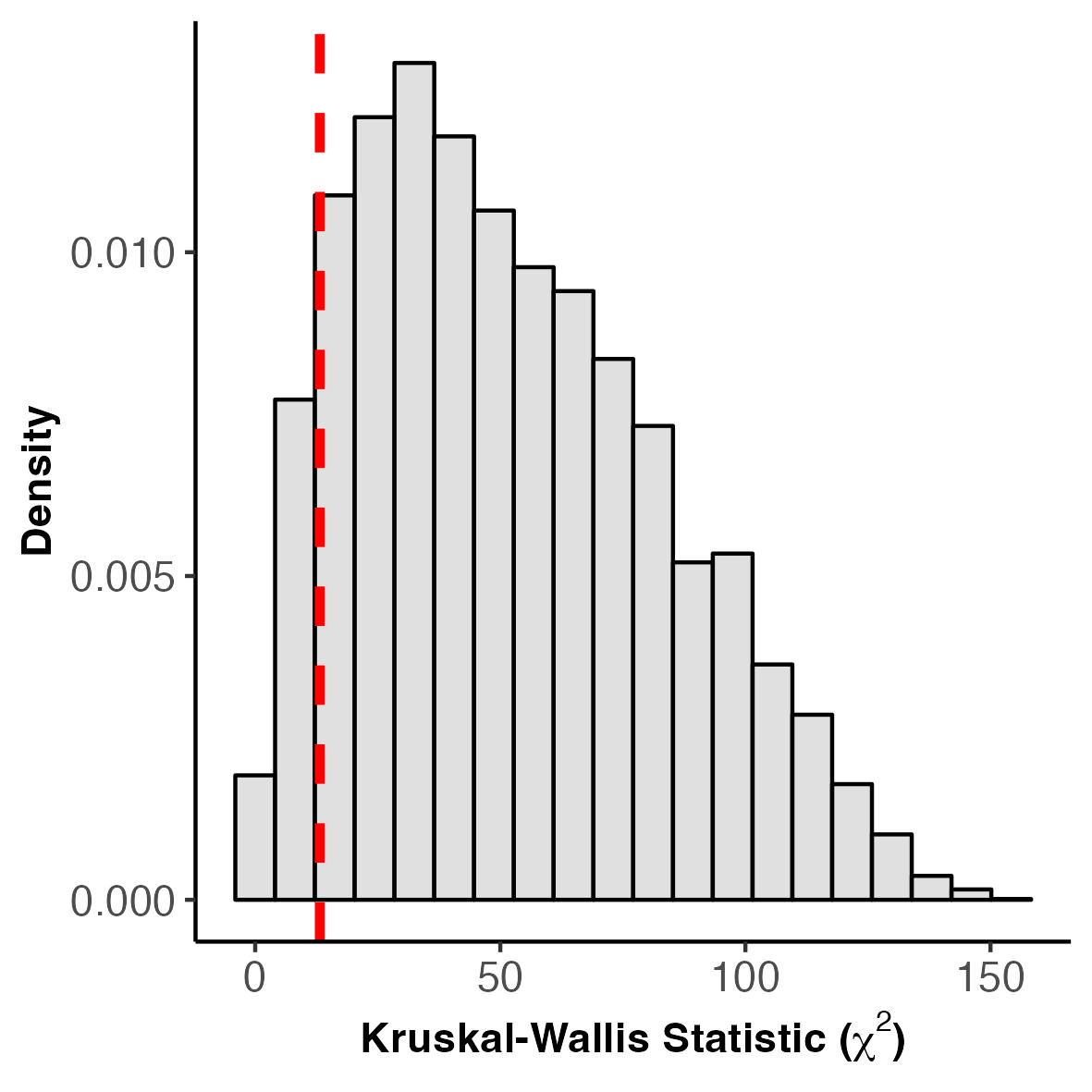

### simulated_chi_sq_distribution_total_count_cv_total_count.jpeg

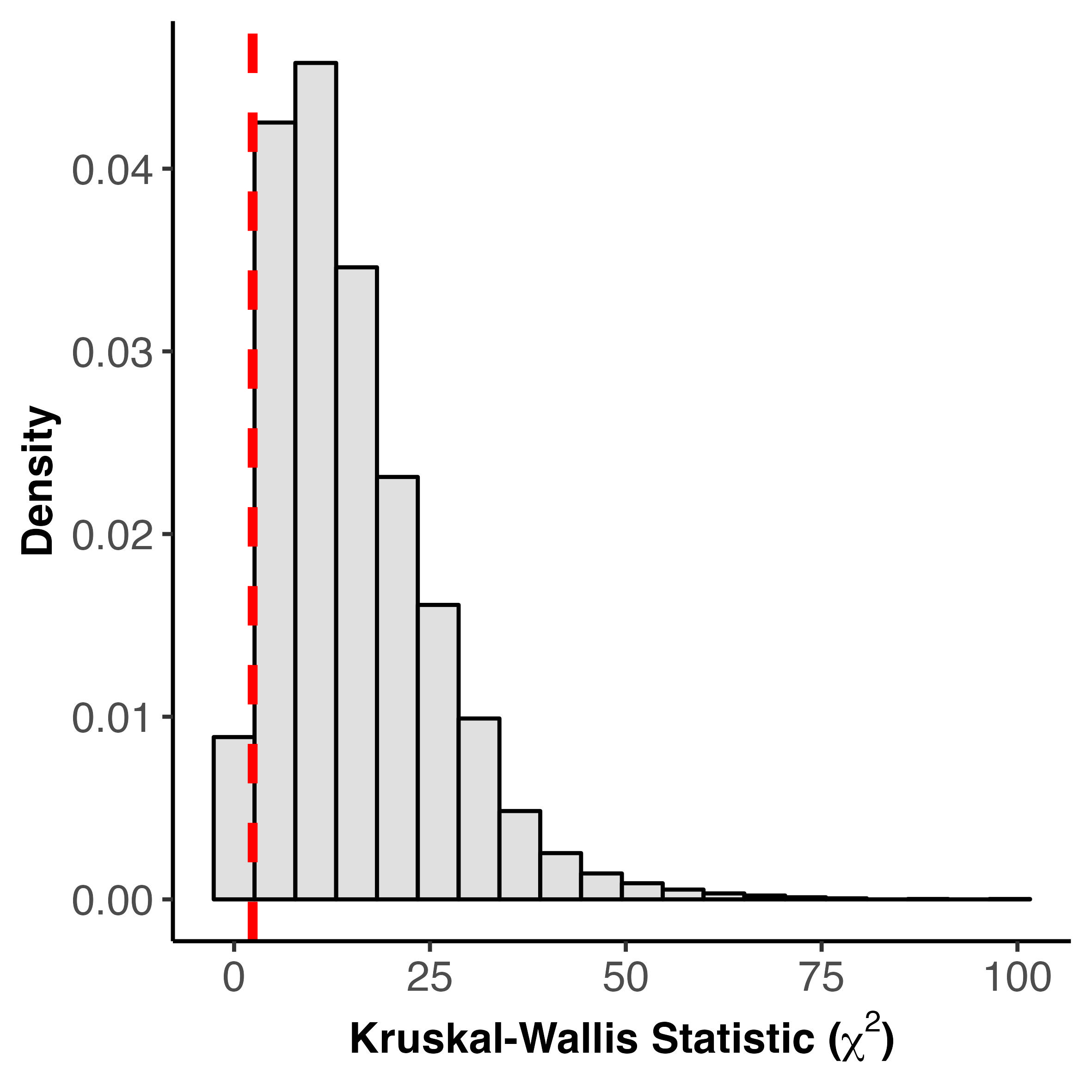

### simulated_chi_sq_distribution_total_count_water_system.jpeg

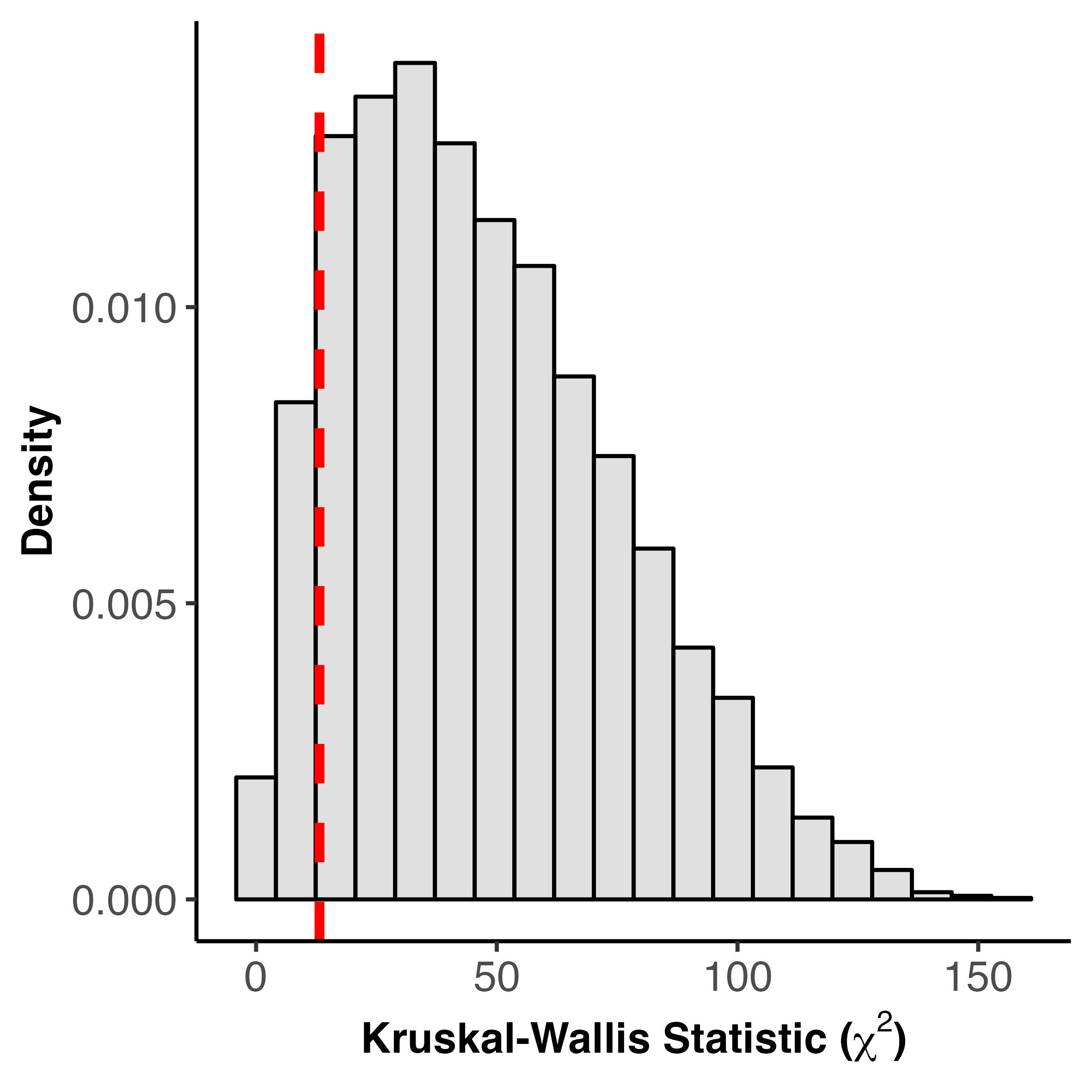

### slopes_distribution_metaanalysis_ci.jpeg

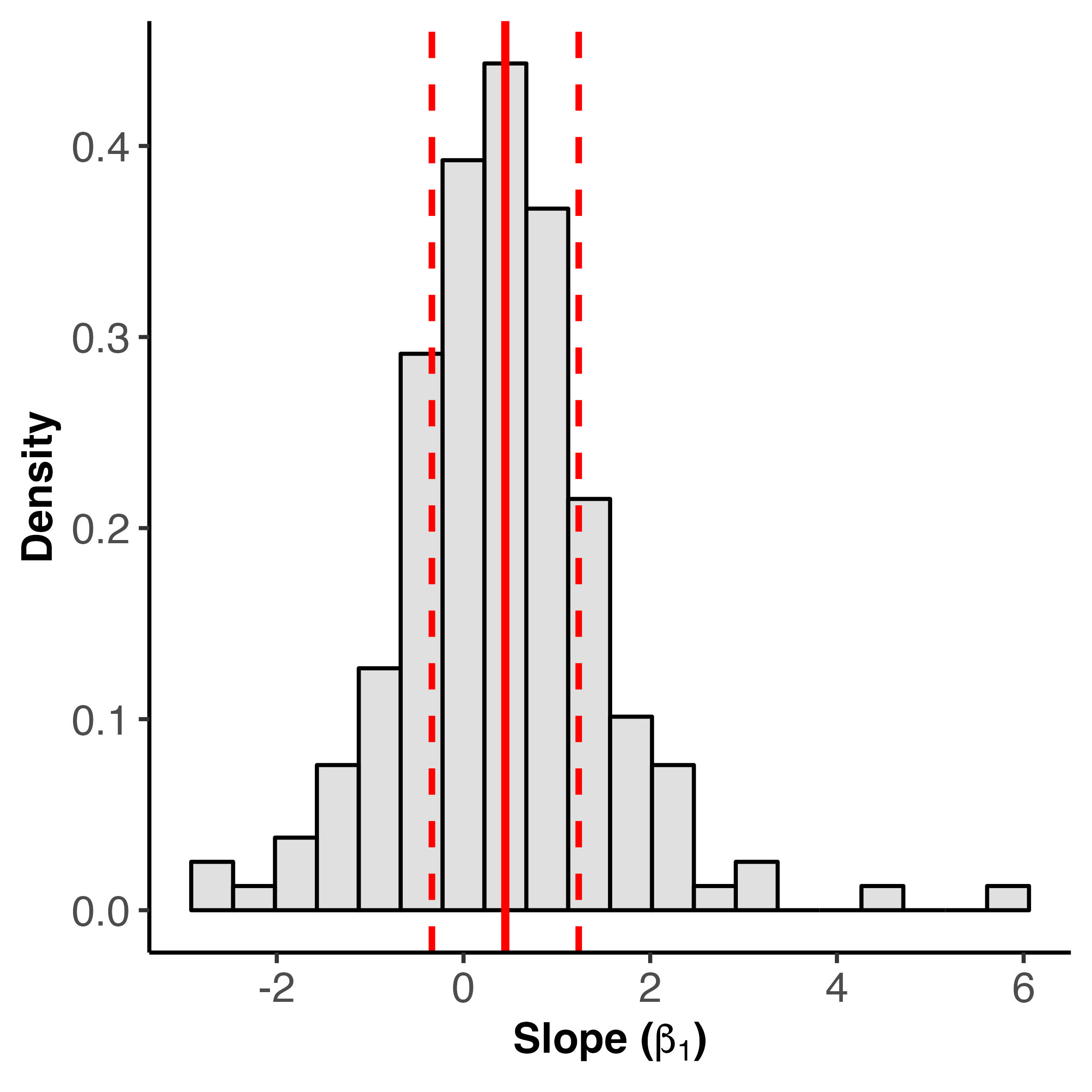

### slopes_versus_slope_sampling_variance.jpeg

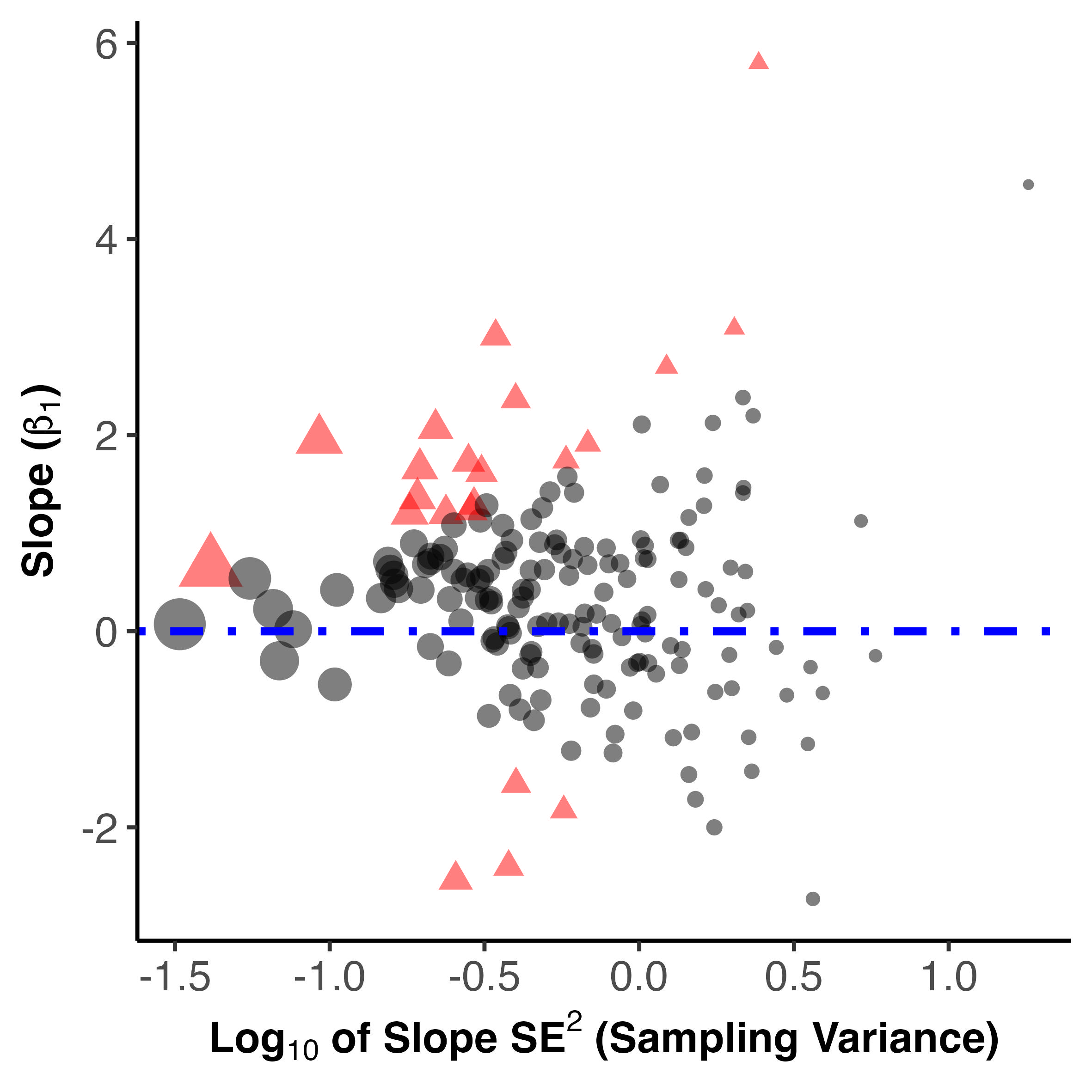

### slopes_versus_slope_sampling_variance_with_legend.jpeg

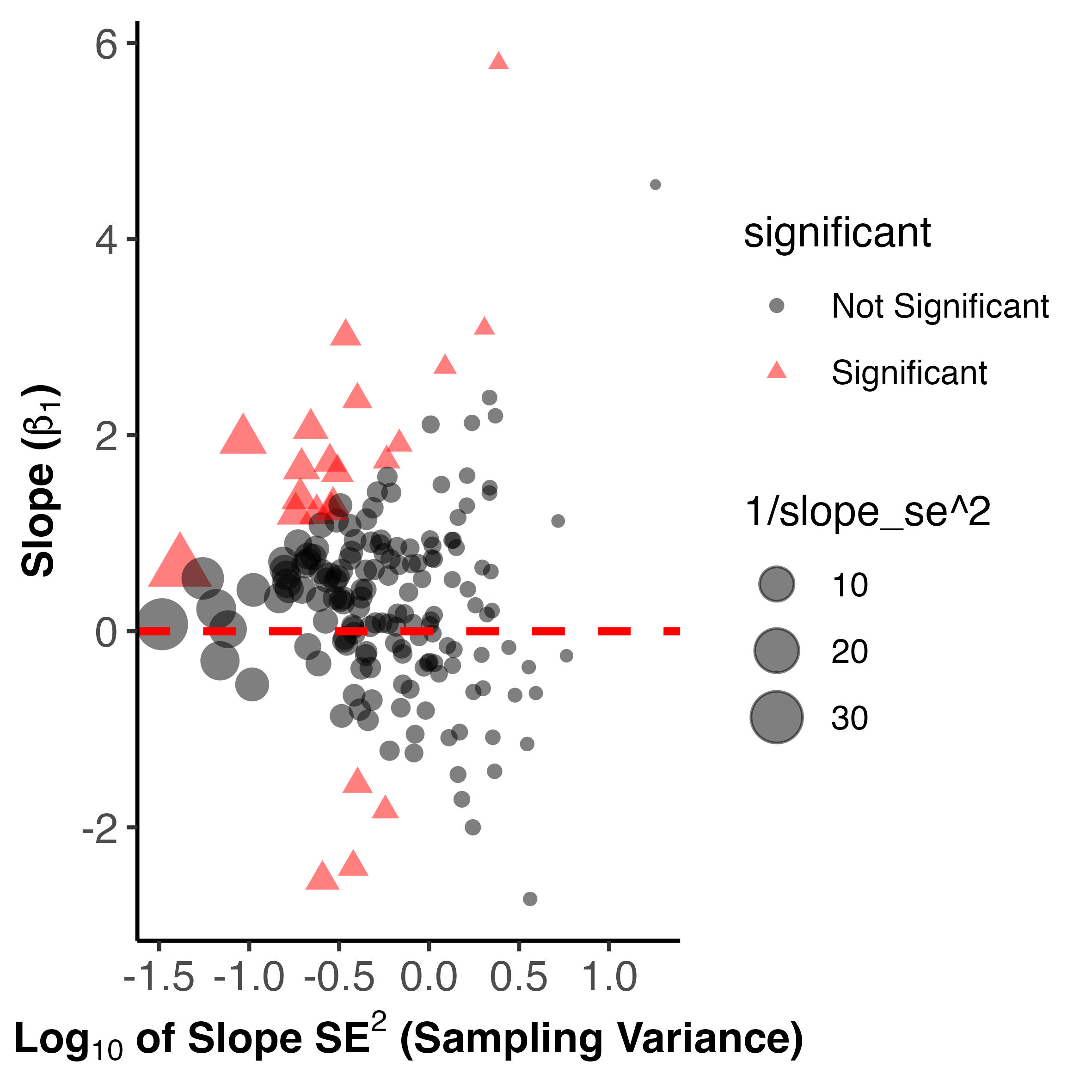

### total_count_by_aspecr_ratio_clean.pdf

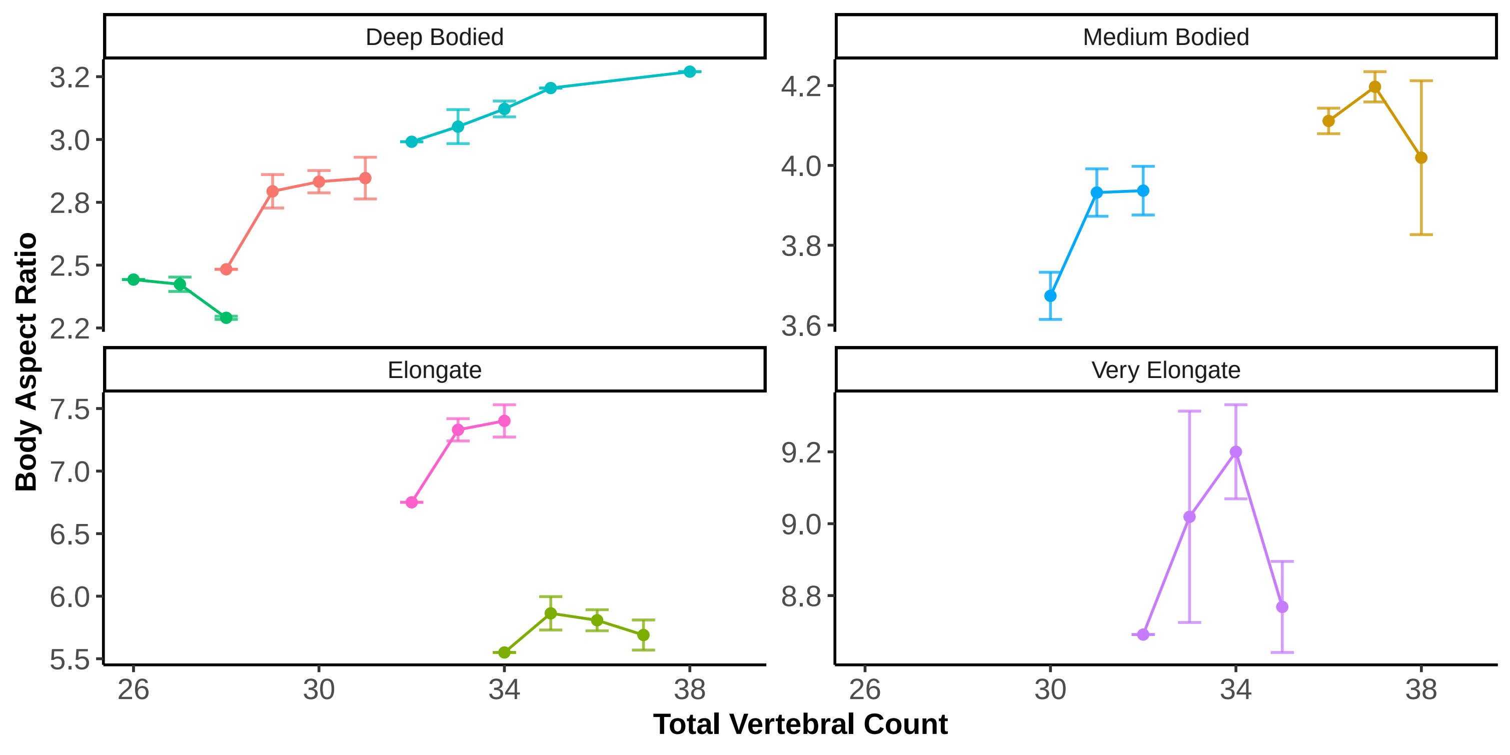

### total_count_by_aspecr_ratio_full.pdf

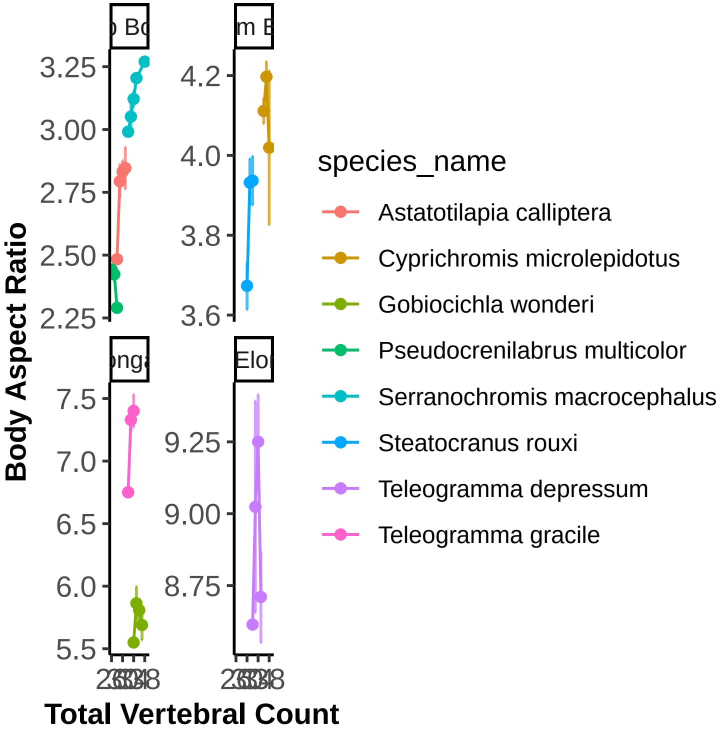

### total_count_by_aspect_ratio_clean.jpeg

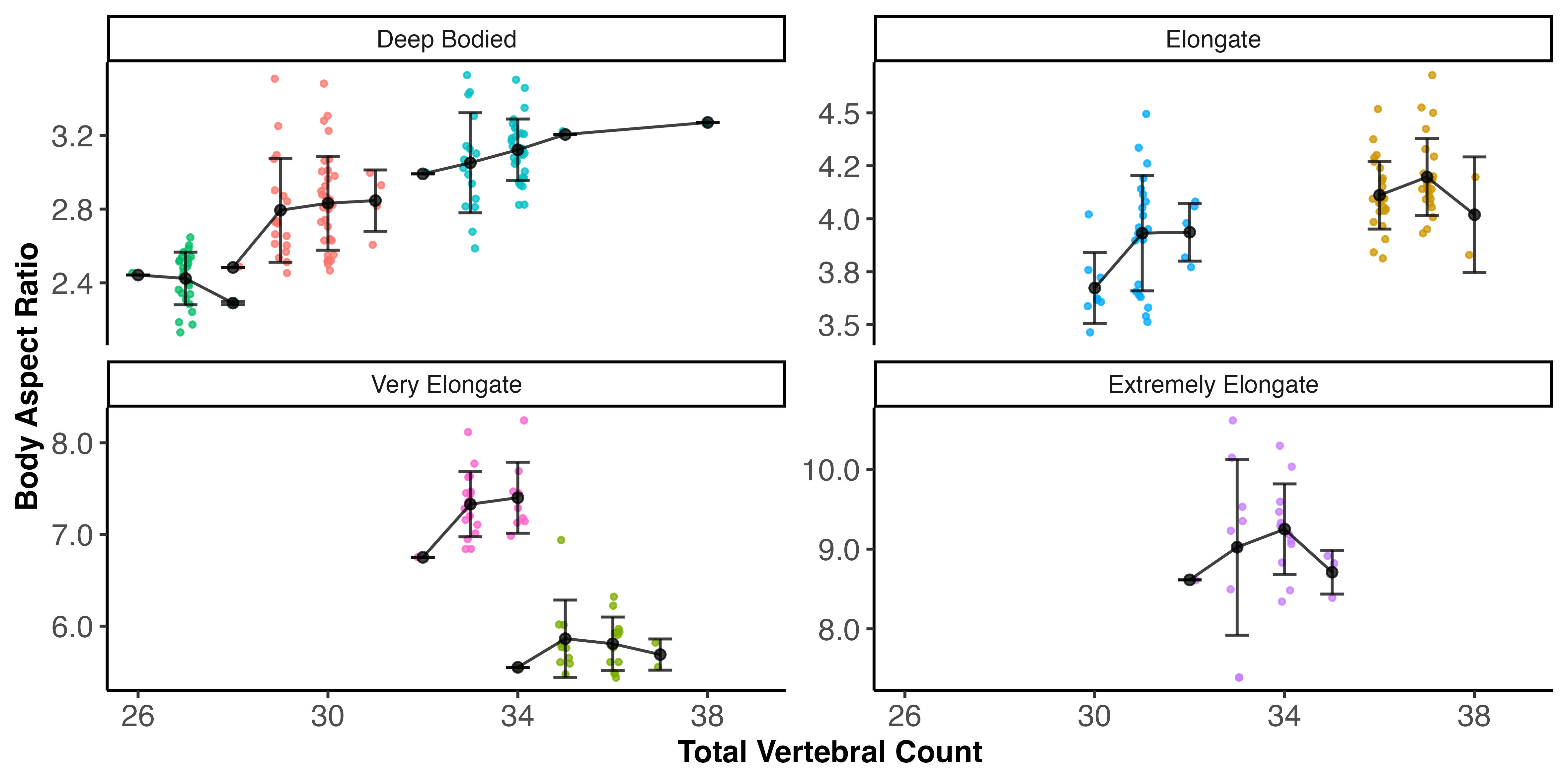

### total_count_by_aspect_ratio_clean.pdf

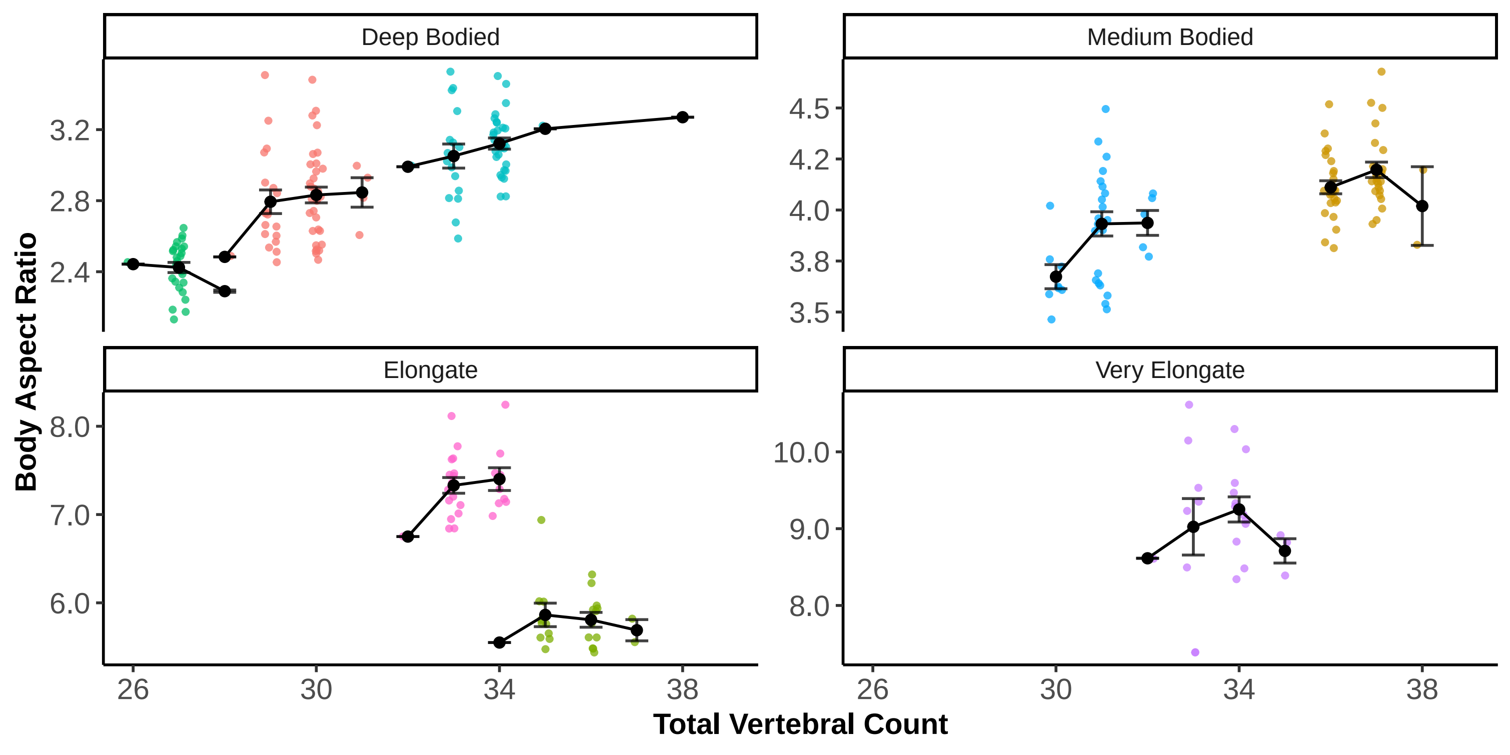

### total_count_by_aspect_ratio_full.pdf

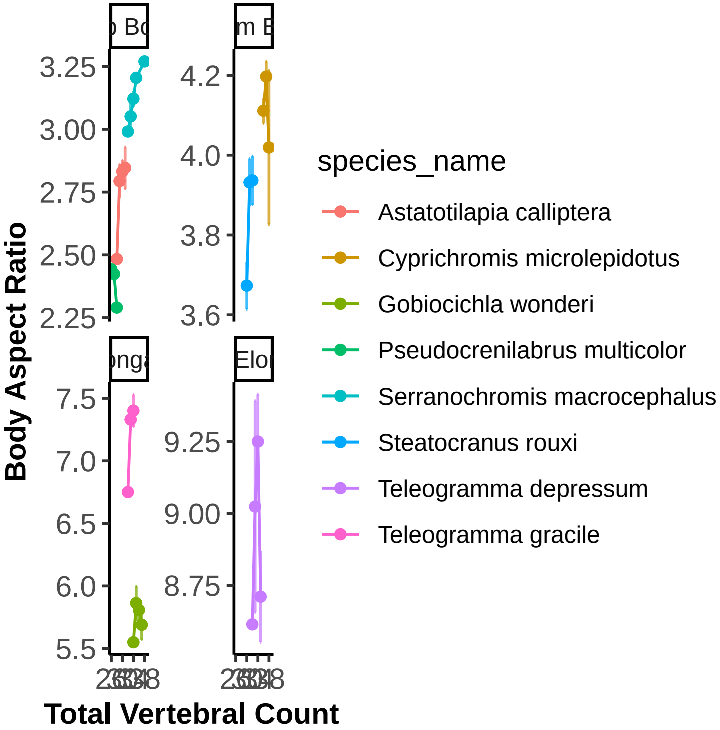

### total_count_cv_by_total_count_bin.jpeg

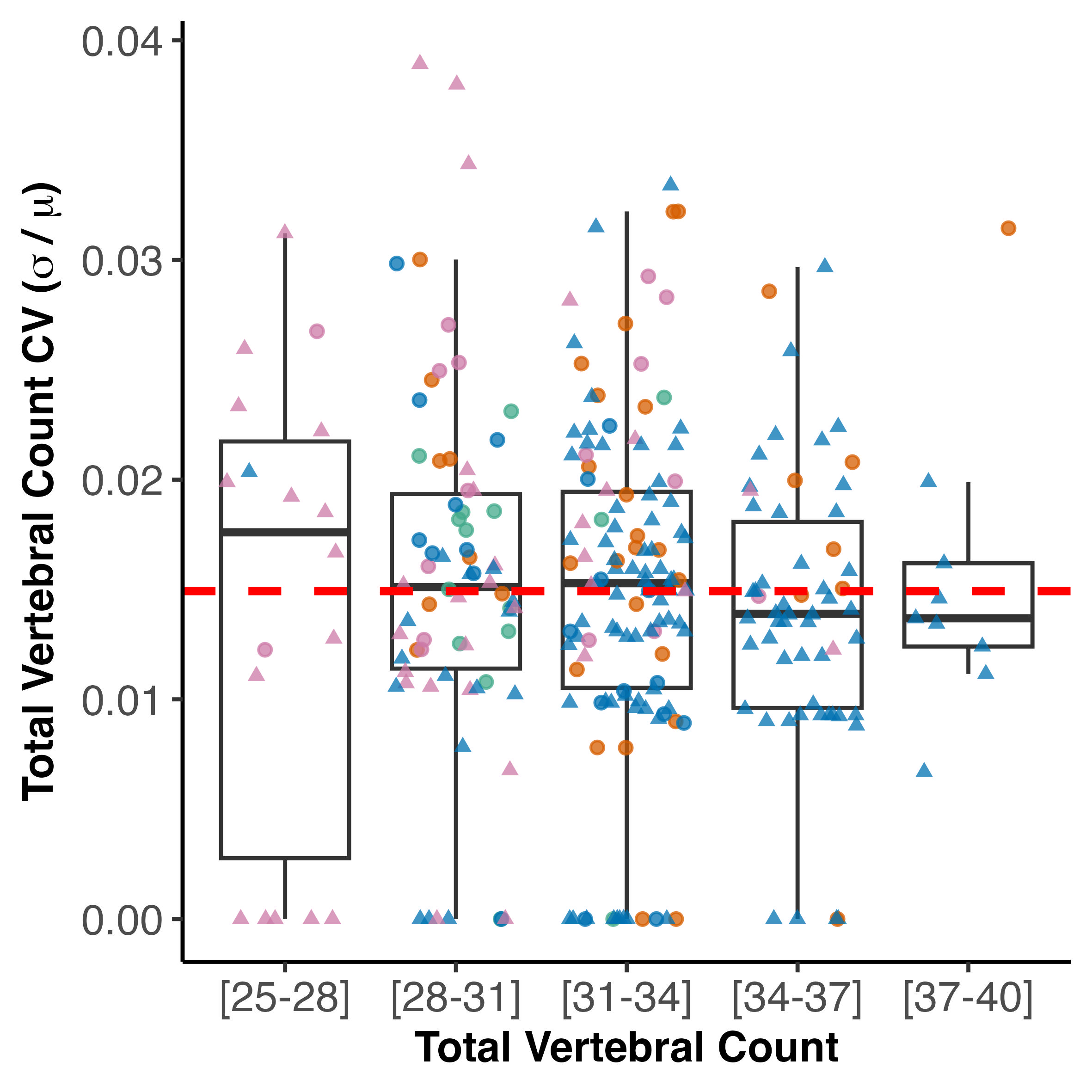

### total_count_cv_by_water_system.jpeg

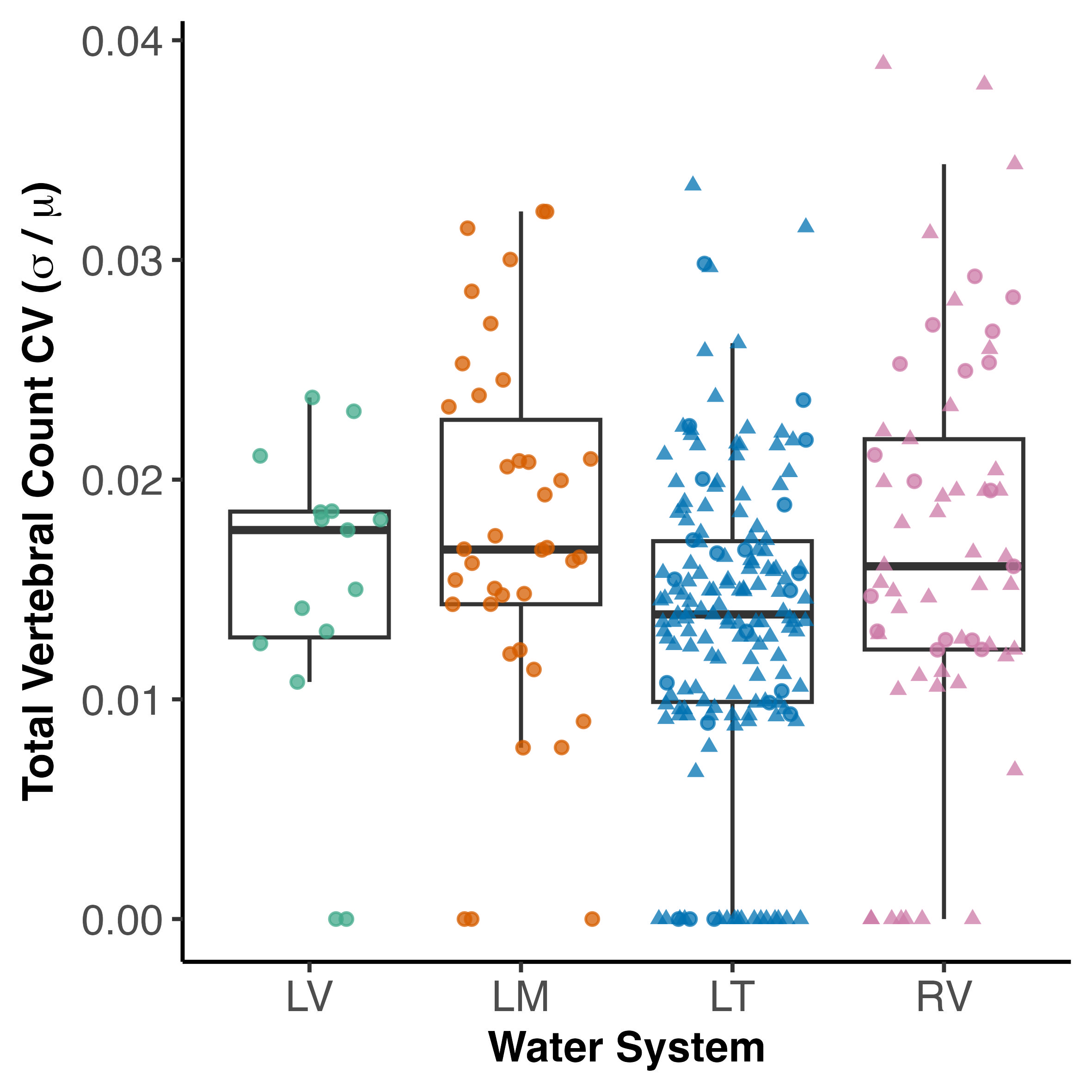

### total_count_cv_distribution.jpeg

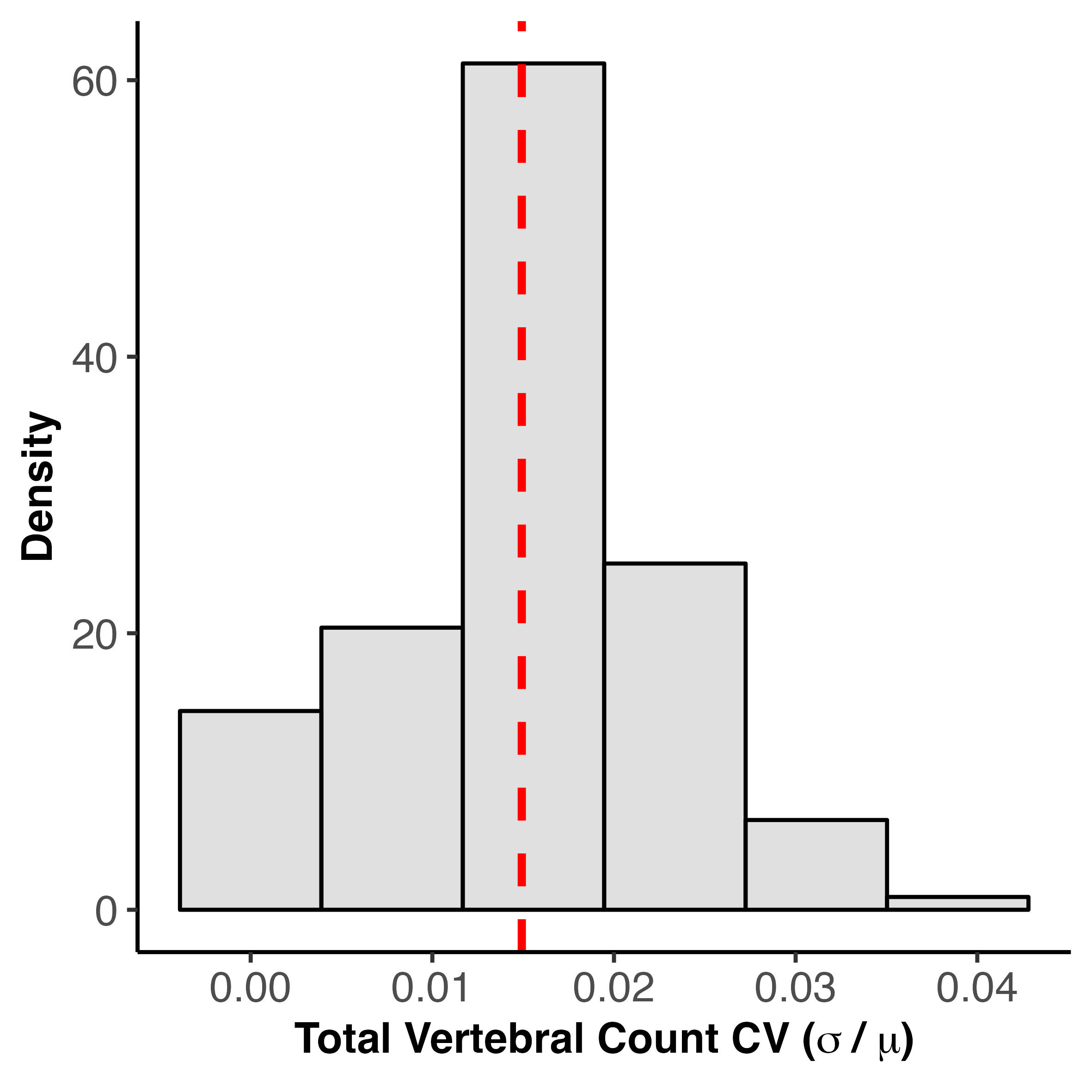

### total_count_cv_k_null.jpeg

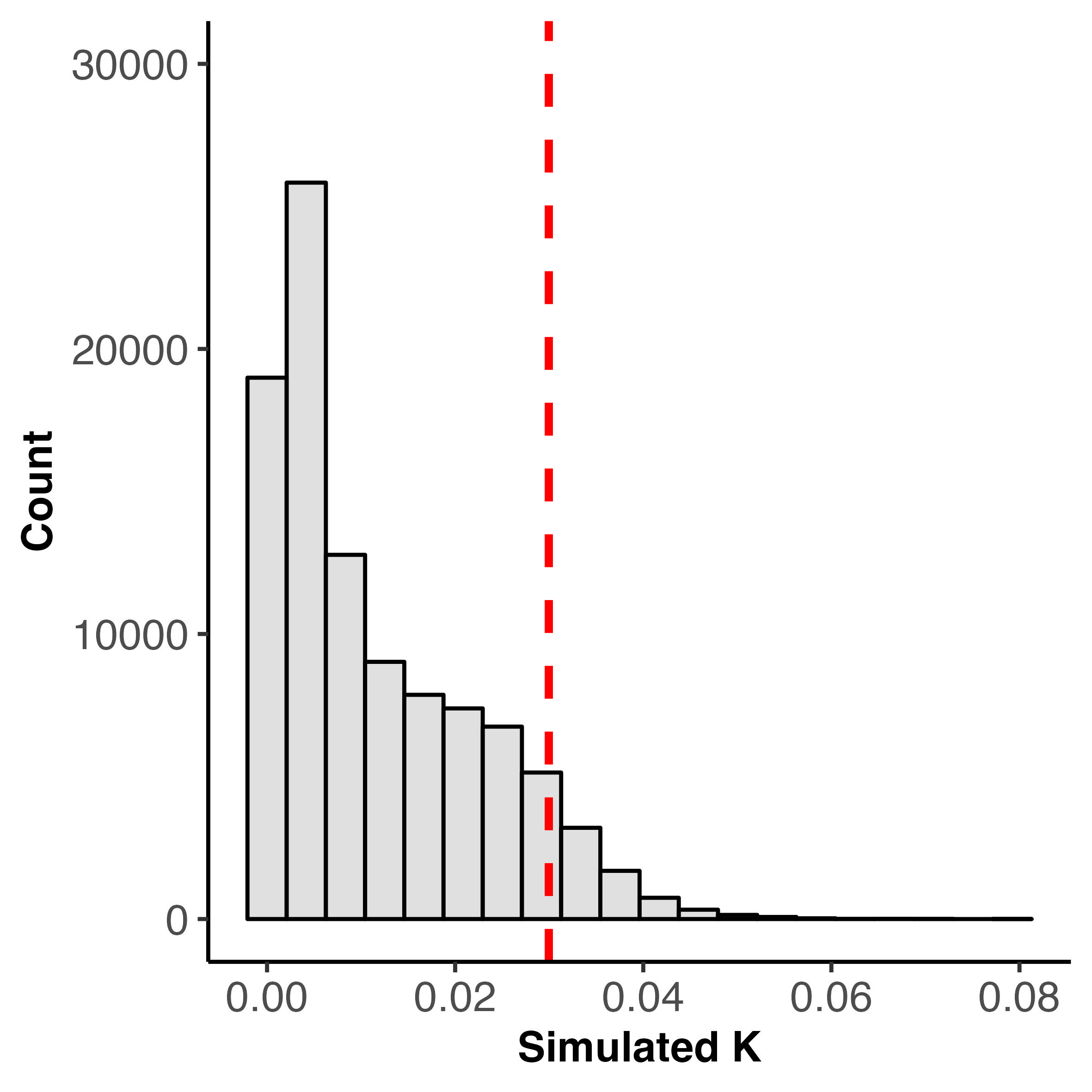

### total_count_cv_versus_total_count.pdf

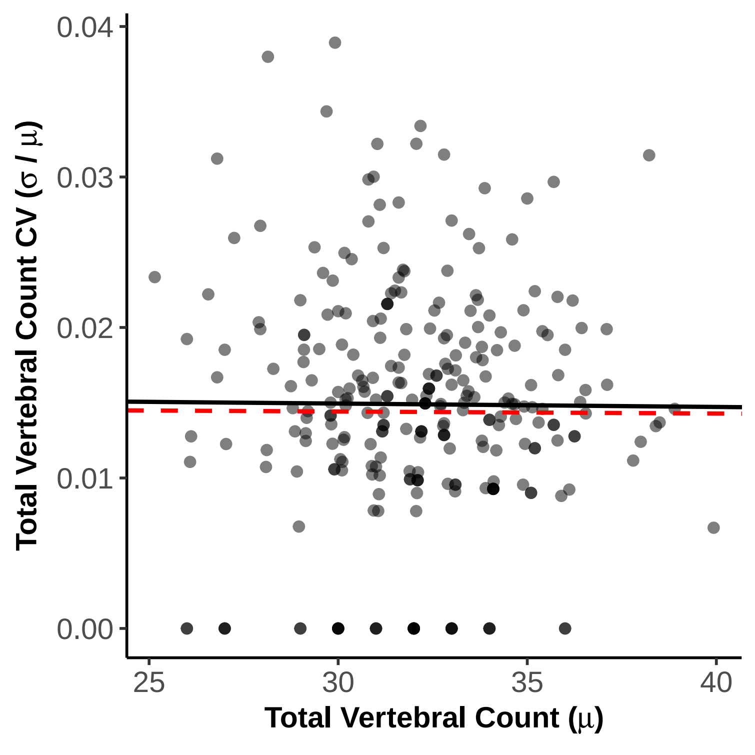

### total_count_cvlambda_likelihood_profile.jpg

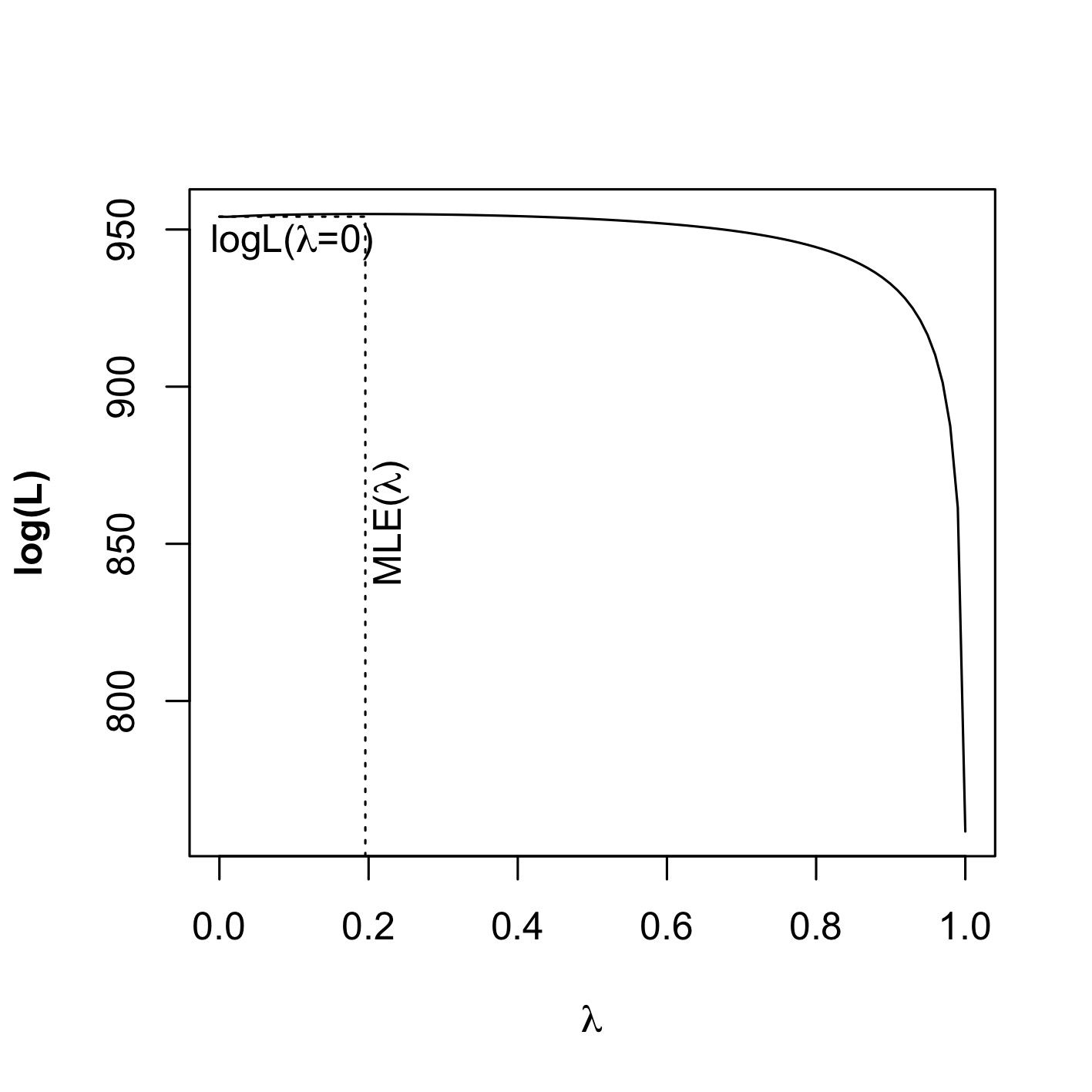

### total_count_distribution_at_least_5.jpeg

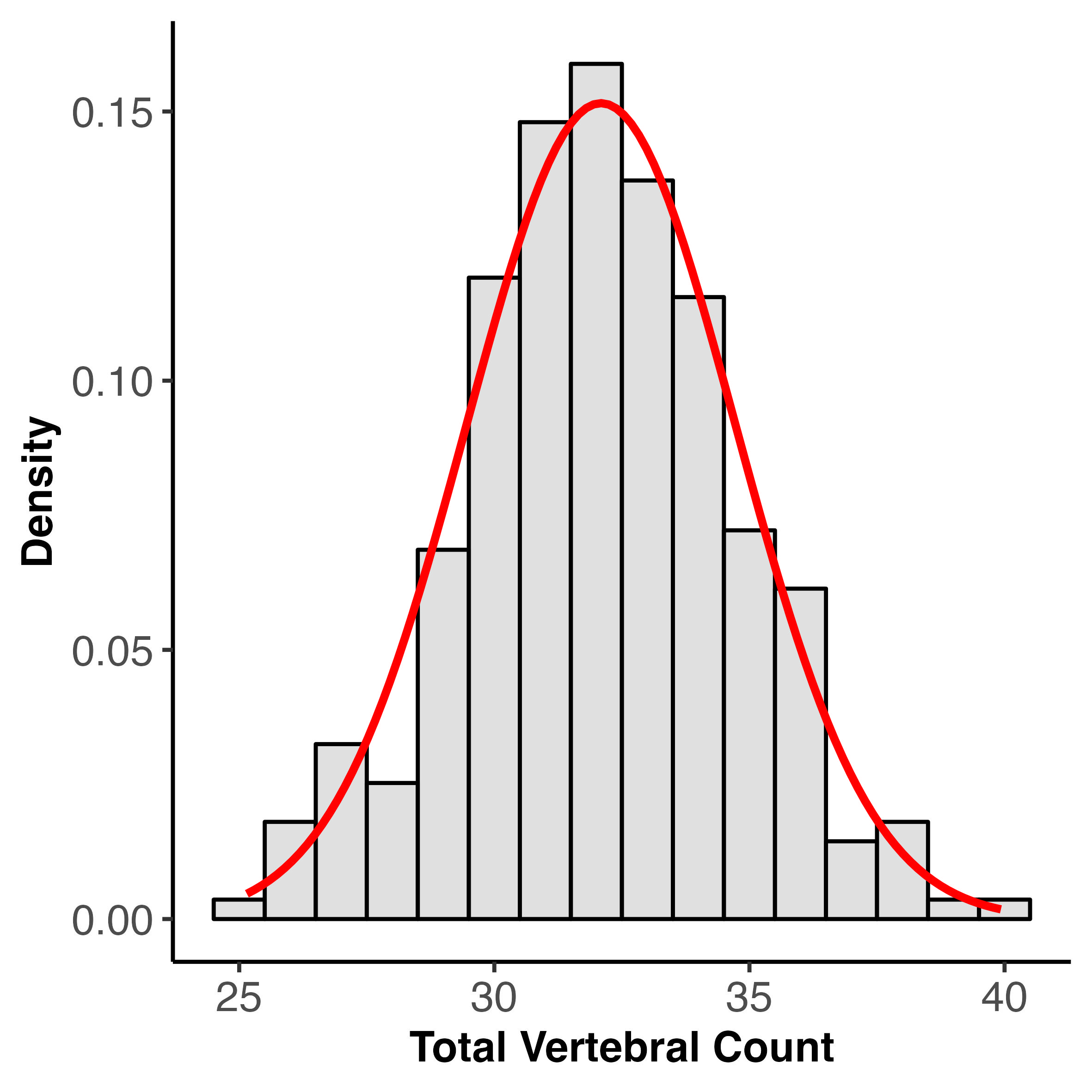

### total_count_k_null.jpeg

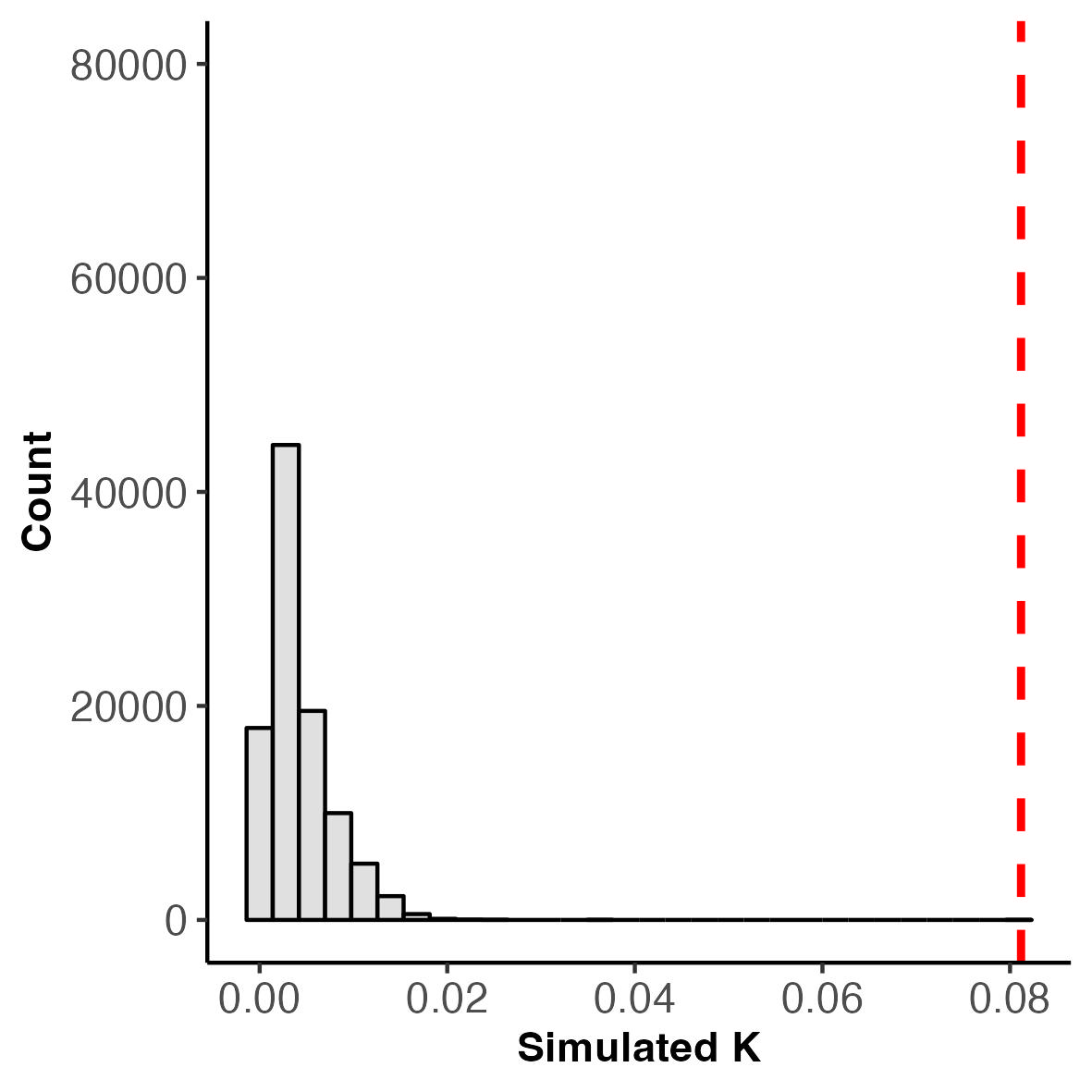

### total_count_lambda_likelihood_profile.jpg

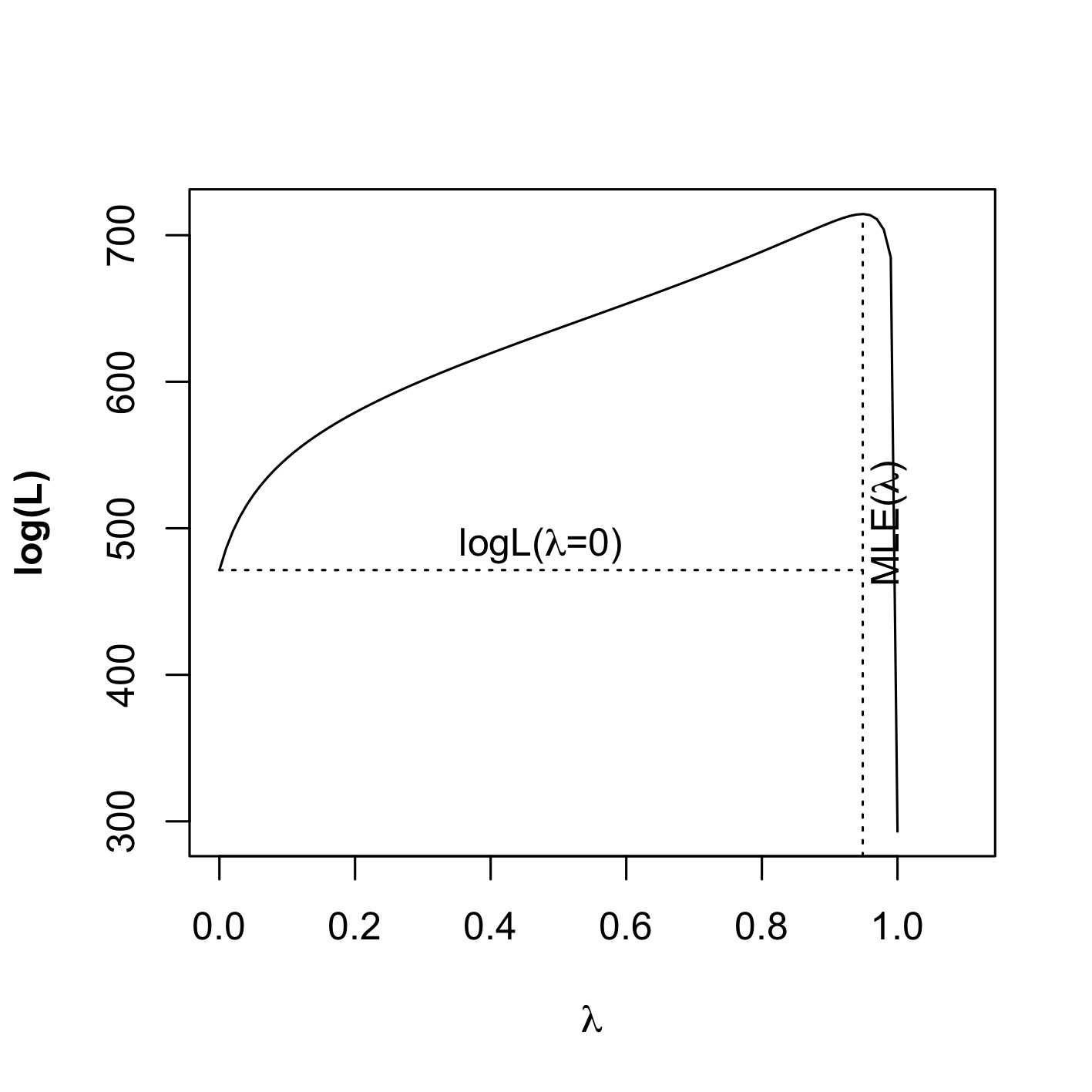

### total_count_pc_c_ratio_pgls.jpeg

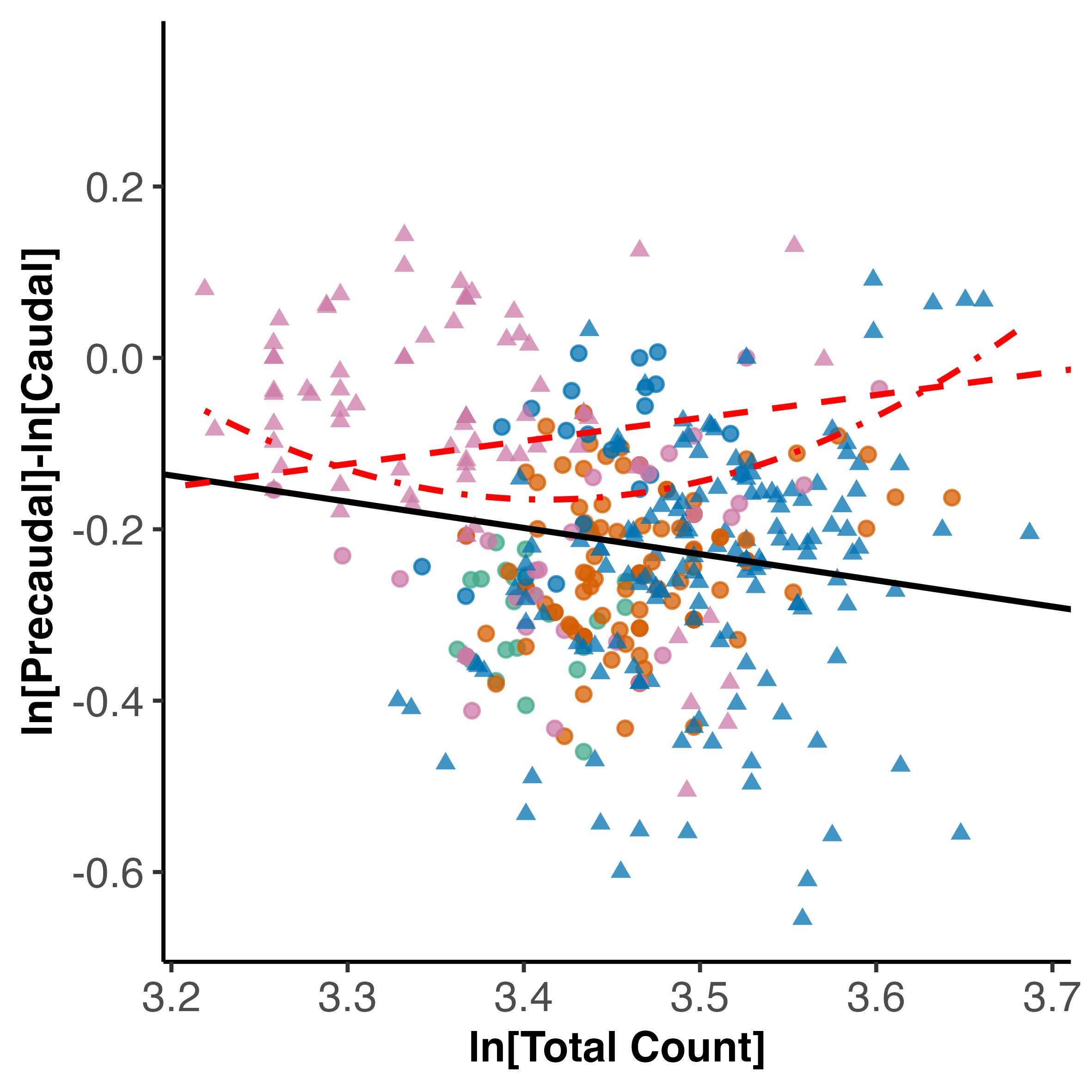

### total_count_pc_c_ratio_pgls.pdf

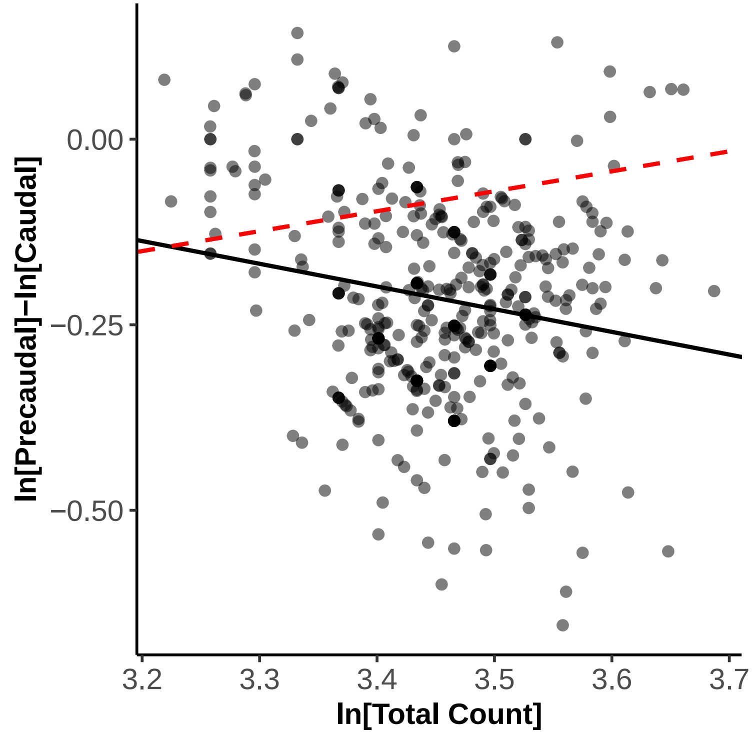

### tree_co_occuring_somitic_homeoetic_effects.jpg

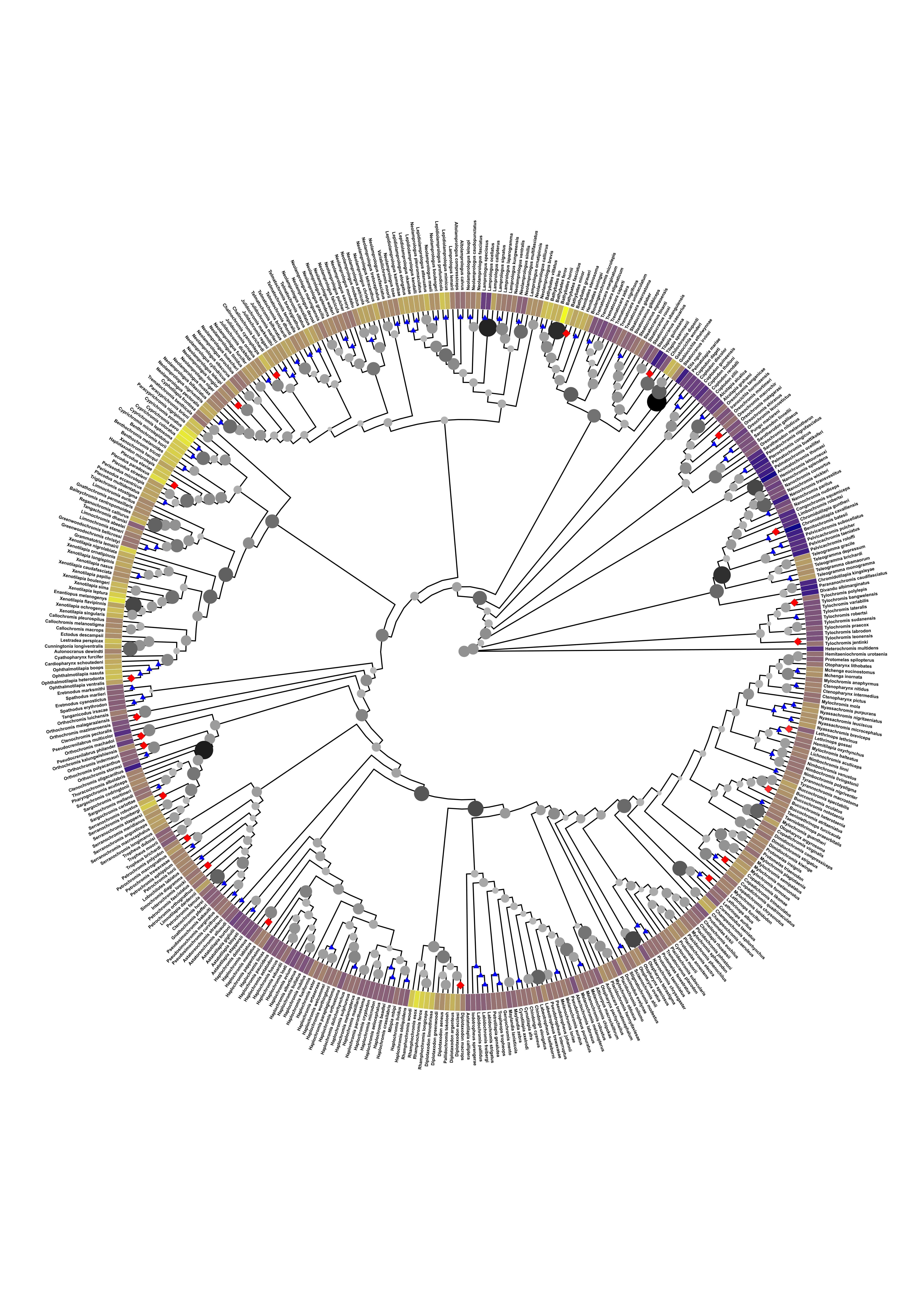

### tree_co_occuring_somitic_homeoetic_effects.pdf

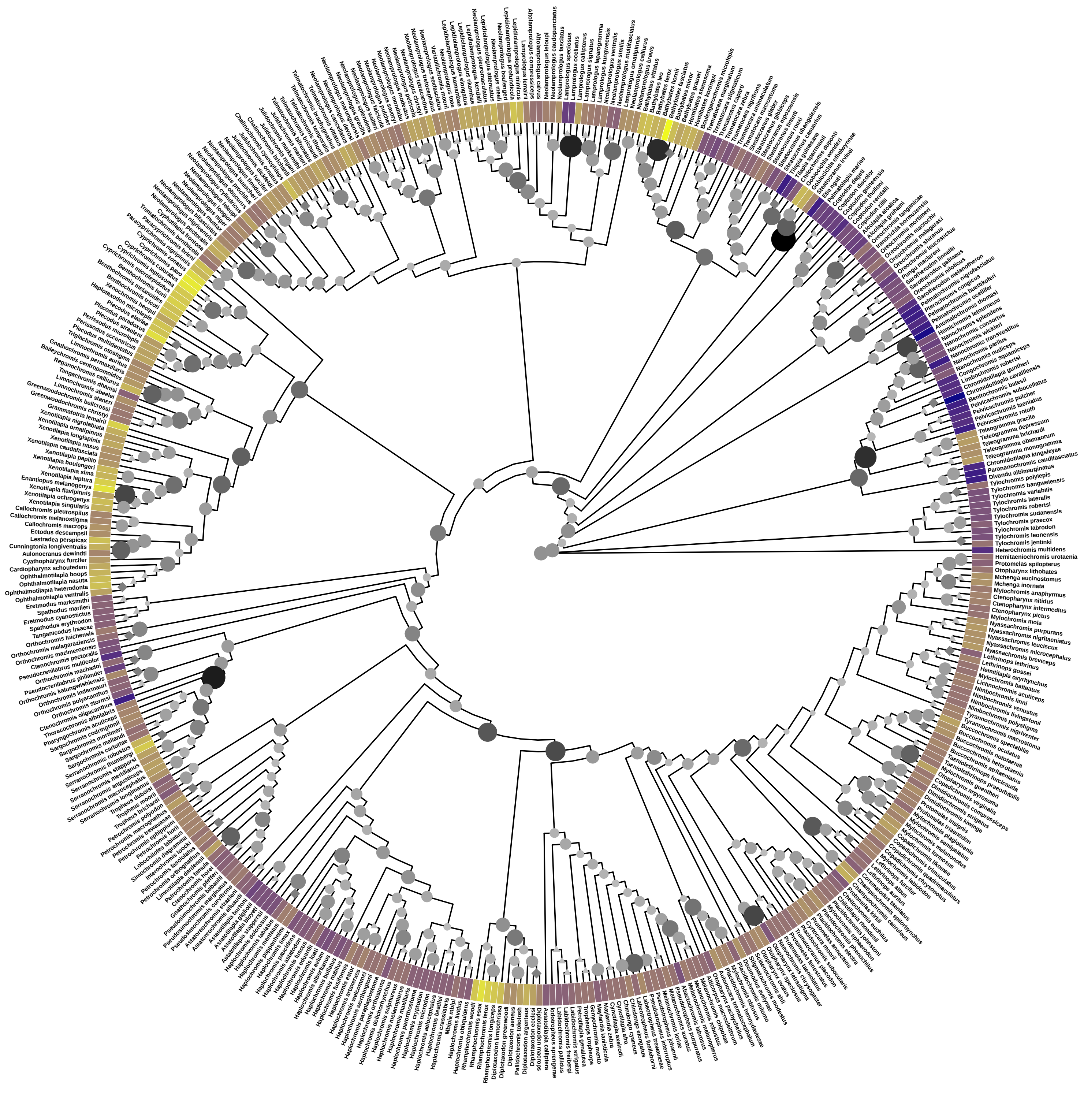

### tree_co_occuring_somitic_homeoetic_effects_time.jpg

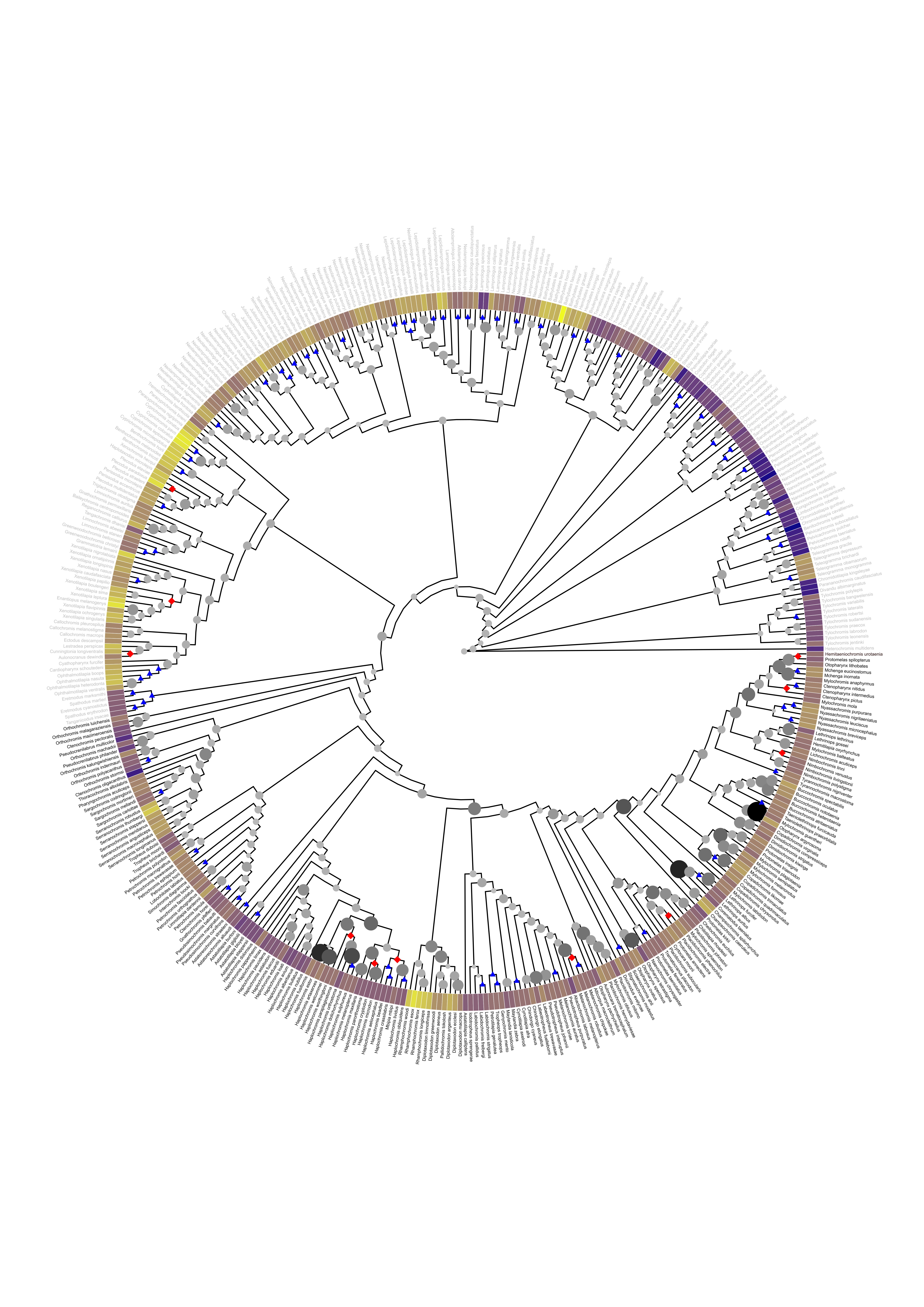

### tree_co_occuring_somitic_homeoetic_effects_time_scale.pdf

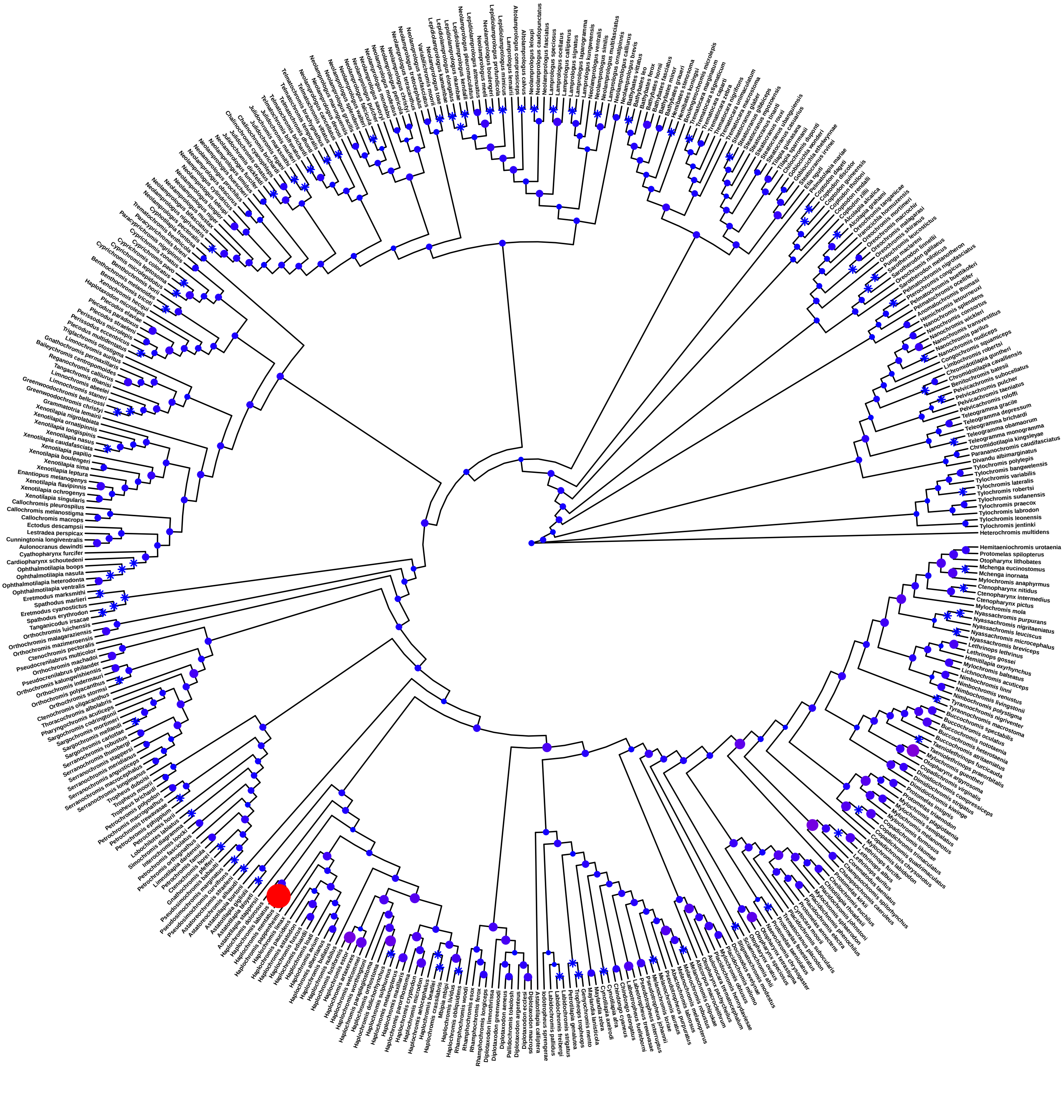

### tree_co_occuring_somitic_homeoetic_effects_with_time.pdf

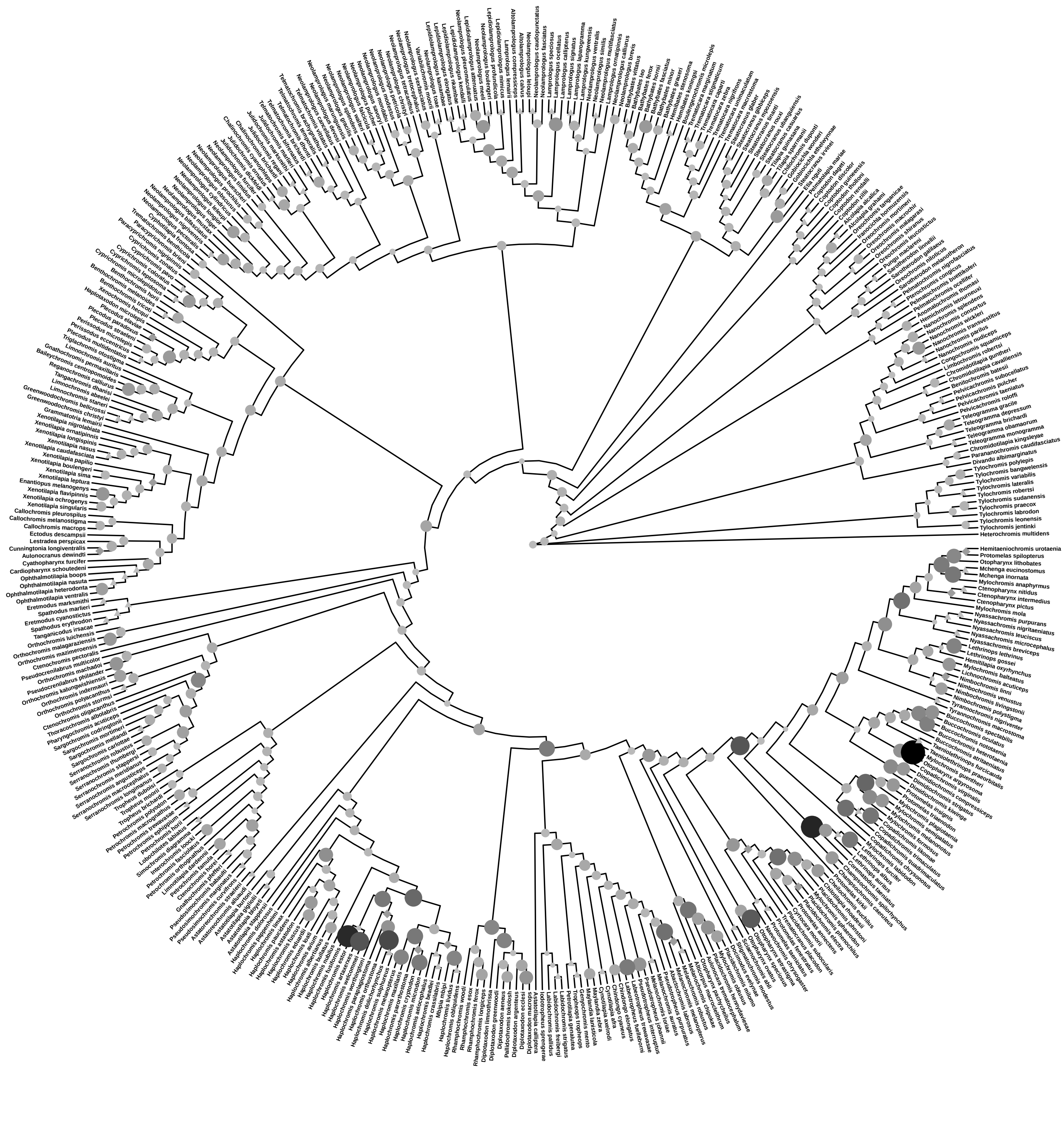

### tree_for_map.pdf

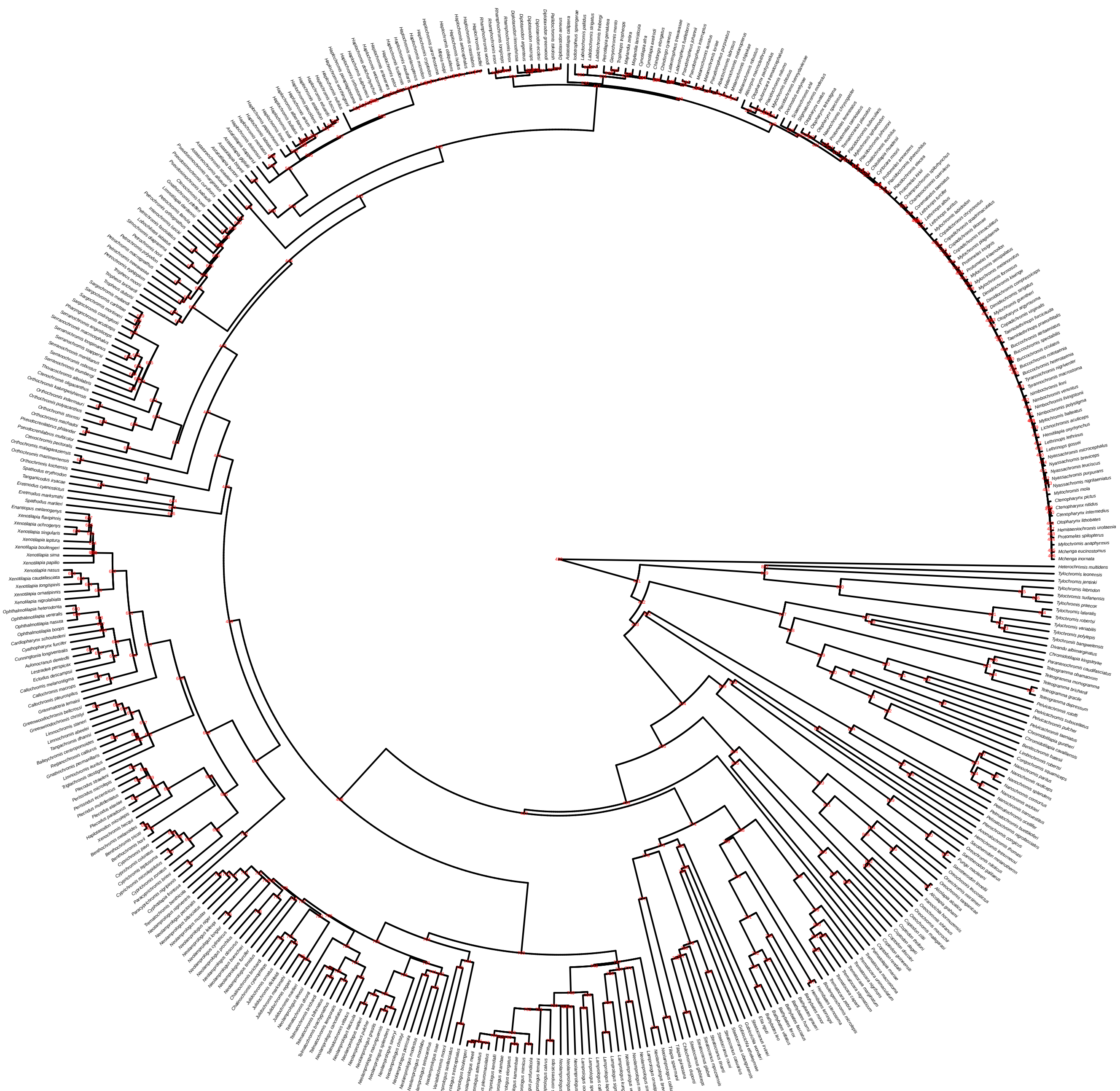
